# Memory erasure by dopamine-gated retrospective learning

**DOI:** 10.64898/2026.01.17.700100

**Authors:** Huijeong Jeong, Leo Zsembik, Farah Farouq, Jennifer A. DiSanto, Chennan Jin, Risha Chakraborty, Nishita Belur, Mingkang Zhou, Andrea D. Sanders, Styra X. Wang, Ananya Srinivasan, Sylvia M. L. Cox, Eric Garr, Sara Brooke, Patricia H. Janak, Marco Leyton, Ritchie Chen, Vijay Mohan K. Namboodiri

## Abstract

Erasing outdated memories is crucial for adaptive behavior. Yet once a cue–outcome association is learned, repeated cue exposure without outcome suppresses conditioned behavior without erasing the underlying memory. This allows rapid behavioral recovery when outcomes are reintroduced. Here, we confirm this limitation for standard “prospective extinction” protocols that present cues without the associated outcome, but show that behavioral and dopaminergic signatures of true memory erasure are achieved by inverting the paradigm: presenting outcomes without associated cues, i.e., “retrospective extinction”. We demonstrate that that mesolimbic dopamine activity at outcome is necessary for retrospective learning and that orbitofrontal cortex activity at outcome is necessary for the rapid behavioral recovery following prospective extinction. These findings reconceptualize extinction mechanisms and identify complementary routes to erase maladaptive memories or limit their relapse-like expression.

## Introduction

Unlearning outdated or harmful memories is as critical as forming new ones, enabling adaptive behavior when cues no longer predict outcomes. For instance, when trauma-associated cues no longer predict danger (referred to as extinction of cue-trauma pairing), persistent cue-trauma memories can still provoke distress. Similarly, persistent cue-drug memories allow drug-associated environmental cues to trigger craving despite abstinence. These enduring memories drive chronic relapse cycles (*1–3*). Standard extinction—repeated cue presentation without the outcome—suppresses conditioned behavior but leaves the underlying memory intact (*4–7*), as shown by rapid reacquisition with minimal cue-outcome re-exposure (**Fig. 1A**) (*8–10*) or spontaneous recovery of conditioned behavior (*11–13*). Although modified extinction procedures have been reported to reduce behavioral recovery compared to standard extinction (*14–25*), results remain variable across studies (*26–35*), leaving unclear what conditions genuinely induce unlearning. Identifying concise, reproducible conditions that truly erase memories could transform interventions for post-traumatic stress disorder (PTSD), addiction, and related disorders involving maladaptive memories.

**Fig. 1.**
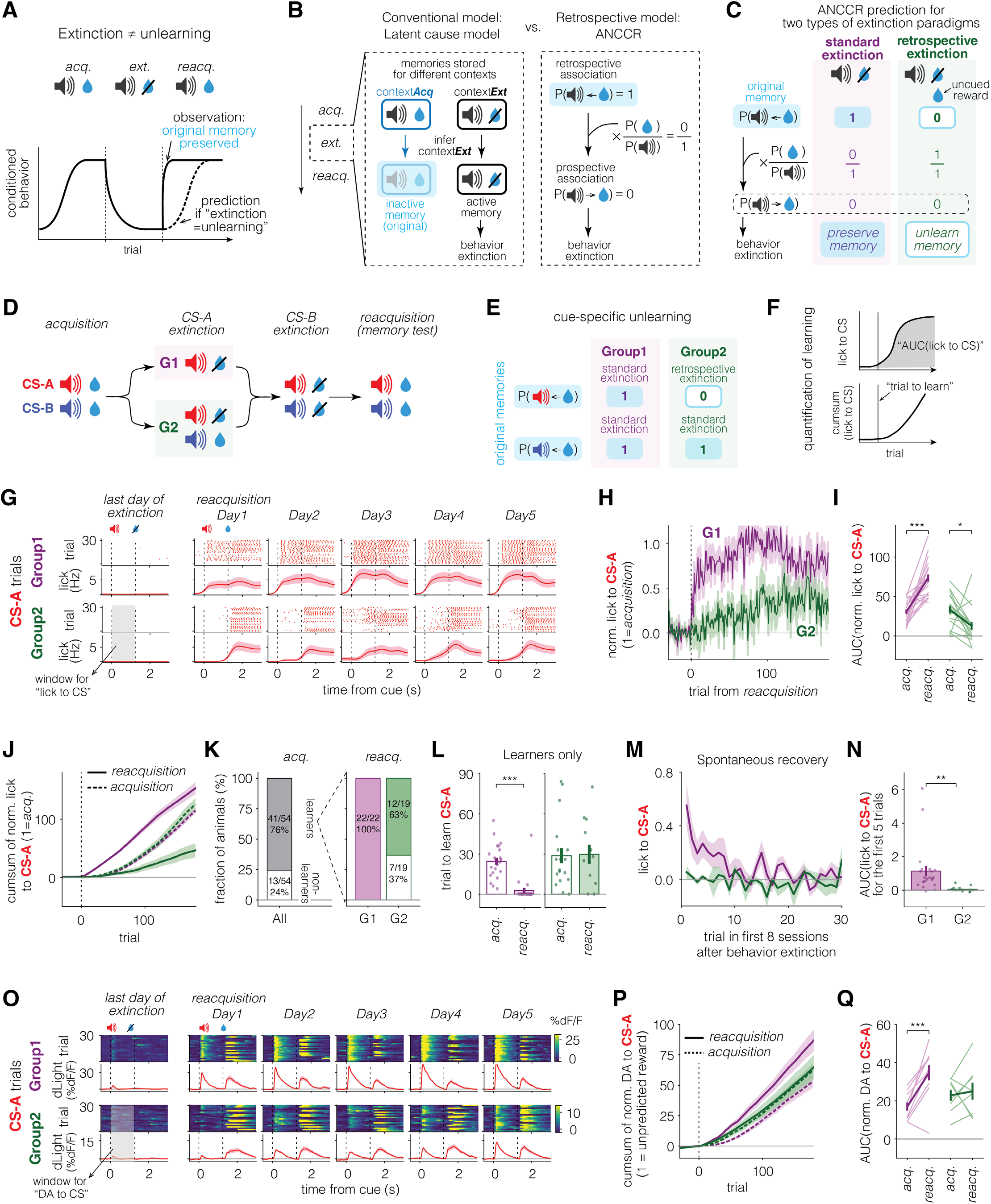
Retrospective extinction degrades original memory. **(A)** Rapid reacquisition compared to original acquisition demonstrates that extinction does not erase the original memory. This raises two questions: how is the memory preserved, and if the memory is preserved, why does behavior extinguish? **(B)** Hypotheses for how memory is preserved during extinction, proposed by two models: Latent cause model and Adjusted net contingency of causal relation (ANCCR). Latent cause models propose that distinct memories of the same cue can be stored in different inferred contexts (e.g., a cue-reward memory in contextAcq and a cue-no reward memory in contextExt). A shift in inferred context from contextAcq to contextExt enables behavior extinction despite preservation of the original cue-reward memory in contextAcq, whereas a shift back from contextExt to contextAcq enables rapid reacquisition. In ANCCR, the prospective probability P(cue→reward) is the product (indicated by ×) of the retrospective probability P(cue←reward) and the normalization factor P(reward)/P(cue). Because the retrospective probability updates only upon reward delivery, it serves as a stable memory that is preserved throughout extinction. However, a decrease in P(reward) during extinction and its rapid restoration during reacquisition drive corresponding decrease and rapid recovery of the prospective probability, and thus behavior. **(C)** ANCCR predicts the same behavior extinction, but with a different underlying mechanism, and motivates two types of extinction paradigms. In standard extinction, the original memory P(cue←reward) is preserved, but a reduction in P(reward) drives behavior extinction (as shown in B). In retrospective extinction, reward delivery dissociated from the cue (either preceded by another cue or delivered without any cue) degrades the retrospective association, resulting in unlearning. **(D)** Task schematic. Mice first learned to associate both CS-A and CS-B with reward. This was followed by CS-A extinction (differential across groups), CS-B extinction (identical), and then reacquisition for both cues (identical). **(E)** Task design for testing two types of extinction. During CS-A (red cue) extinction, Group1 (G1) underwent standard extinction, while Group2 (G2) underwent retrospective extinction. CS-B (blue cue) underwent standard extinction for both groups, allowing both between and within animal comparison of memory persistence. **(F)** Behavior was quantified as the change in lick count during the 1.25 s after cue onset (“lick to CS”). Learning was assessed via the area under the curve (AUC) of learning curve or by identifying the inflection point (‘trial to learn’, denoted by vertical line) of the cumulative learning curve. **(G)** Example lick raster plots (top row) and PSTHs (bottom row) for one representative mouse per each group aligned to CS-A trials (top, G1; bottom, G2). The first column shows data from the last day of CS-B extinction, and remaining columns (Day 1 to 5) show reacquisition. Graphs are aligned to the onset of cue, denoted by the first vertical black dotted line. The second black dotted line indicates delivery or omission of reward. G1 animal shows an increase in anticipatory licking before reward on Day 1 of reacquisition, while G2 animal only develops it on Day 4. Grey shaded area indicates the anticipatory window used to calculate “lick to CS” **(H)** Group-averaged normalized lick to CS-A across trials during reacquisition showing G1 shows faster reacquisition than G2. Trial 0 marks the first CS-A trial in reacquisition. Each animal’s lick to CS-A was normalized to its average during the last 3 days of acquisition. Lines represent mean across animals and shadings represent SEM (same as other panels). **(I)** Reacquisition is more rapid than acquisition only in G1, not in G2. AUC of normalized lick to CS-A over the first 6 days of acquisition and reacquisition. Error bar represents SEM across animals and light-colored lines represent individual animals. **(J)** Group-averaged cumulative sum of normalized lick to CS-A during acquisition (dotted) and reacquisition (solid). Trial 0 marks the first trial of each respective phase. For reacquisition, CS-B extinction trials are shown at trial numbers <0. **(K)** Fraction of learners (filled bar) and non-learners (empty bar) for each group during acquisition and reacquisition. Animals from G1 and G2 went through acquisition similarly. Only animals that learned during acquisition (n = 41) were divided into groups and included in subsequent experiments. **(L)** Reacquisition was faster than acquisition in G1 but similarly slow in G2 when non-learners were excluded (7 of 19 G2 animals during reacquisition). Error bar represents SEM across animals and individual animals are shown as scatter. **(M)** G1 shows stronger spontaneous recovery than G2. Lick to CS-A across trials within an extinction session, averaged across the first 8 sessions after behavior is extinguished (See Methods). Each data point represents the average lick response at a given trial number across sessions, then averaged across animals. **(N)** AUC across first 5 trials of the trace in M, quantifying the magnitude of spontaneous recovery. **(O)** Example heatmaps (top) and PSTHs (bottom) of dopamine release (dF/F) in the nucleus accumbens in response to CS-A, from a representative G1 and G2 mouse. Dopamine response to cue appears faster in G1 than G2 consistent with behavior. Grey shaded area indicates the anticipatory window used to calculate “DA to CS” **(P)** Reacquisition of dopamine response is more rapid than acquisition only in G1, not in G2. Group-averaged cumulative sum of normalized dopamine (DA) to CS-A during acquisition (dotted) and reacquisition (solid). DA to CS-A was calculated as AUC of dF/F over the 1.25 s cue-reward window, normalized to the maximum reward response (AUC over the 1.25 s window after reward) during the first 3 days of acquisition. **(Q)** AUC of normalized DA to CS-A during the first 6 days of acquisition and reacquisition. *: p<0.05, **: p<0.01, ***: p<0.001, n.s: non-significant in all figures. <u>See</u> **Supplementary table 2** <u>for statistical results from all figures.</u>

The maintenance of memory despite behavior extinction raises two questions: how is the original memory maintained during behavior extinction, and how is its behavioral impact suppressed during extinction? The widely accepted explanation is that extinction forms a new memory that competes with, instead of replacing, the original memory (*5–7*, *11*, *36*) (**Fig. 1B**, Conventional model). Latent cause models formalize this idea by proposing that animals learn cue-outcome (e.g., cue-reward) relationships for inferred hidden “contexts” even when physical contexts do not change (*37–41*). These models need to assume hidden contexts to explain memory maintenance during extinction, because cue-outcome relationships in these models are prospective, storing the prediction of what follows the cue, i.e., P(cue→reward). To account for behavior extinction while preserving the original cue-reward relationship, different cue-outcome contingencies are associated with different distributions over inferred latent contexts. In the simplest case where each contingency is associated with a single latent context, the acquisition context predicts [P(cue→reward | contextAcq) = 100%] during acquisition, whereas during extinction a new extinction context predicts [P(cue→reward | contextExt) = 0%]. Therefore, an inferred switch from contextAcq to contextExt produces behavioral extinction despite the original memory being preserved, while a switch back to contextAcq through rapid inference during reacquisition enables fast restoration of conditioned behavior. Although latent cause models differ in how latent causes are represented and inferred (e.g., discrete vs. probabilistic Bayesian formulations), they share the central logic that behavior following extinction depends on inference over latent causes rather than modification of a single cue-reward memory.

An alternate view comes from retrospective learning models, i.e., models learning that cues preferentially *precede* meaningful outcomes, as exemplified by Adjusted Net Contingency of Causal Relations (ANCCR) (**Fig. 1B**, Retrospective model) (*42–44*). Rather than positing multiple hidden contexts, ANCCR assumes that animals only learn and store a single retrospective P(cue←reward) association indexed by physical context. Thus, indexing memories by context is shared by latent cause models and ANCCR; the key difference is that in ANCCR, the context is a physical context defined by sensory cues instead of latent causes inferred mostly from changes in strength of association. After learning a context-indexed retrospective association P(cue←reward), animals infer a prospective P(cue→reward) prediction to drive cue-induced behavior using Bayes’ rule: P(cue→reward) = P(cue←reward)×P(reward)/P(cue). Although prospective and retrospective associations may seem similar, they differ substantially. For example, if a reward is always preceded by a cue but the cue is followed by reward only 10% of the time, the prospective association is weak [P(cue→reward) = 10%] while the retrospective association is perfect [P(cue← reward) = 100%]. This asymmetry means that lowering reward probability all the way to zero during extinction can abolish prospective P(cue→reward) association while leaving the retrospective P(cue←reward) association intact. The retrospective association therefore forms a persistent memory of the previous cue-reward pairing. Behavior, which is guided by the prospective prediction of reward following the cue, is extinguished due to the overall reward rate (i.e., P(reward)/P(cue)) becoming zero during extinction (**Fig. 1C**, standard prospective extinction). The preserved retrospective memory enables rapid restoration of behavior when estimated reward rate rapidly recovers during reacquisition. Note that ANCCR formally computes long-run discounted prospective or retrospective associations. However, for simplicity, we use probabilities to describe prospective and retrospective associations as this simpler notation is sufficient to motivate all experimental design in this paper (see **Supplementary Note 1** and Jeong et al 2022 (*42*) for formal mathematical details).

Intuitively, the retrospective framework helps explain why abstinence from drugs, or extinction-based approaches such as exposure therapy for aversive outcomes, fail to entirely erase maladaptive drug or trauma memories. Repeated cue exposure without outcome teaches that the outcome is unlikely to happen, suppressing conditioned responses to the cue, but leaves intact the memory that whenever the outcome occurs, it is preceded by the cue.

By contrast, cue-reward memory erasure requires abolishing the retrospective P(cue←reward) association. This quantity measures the fraction of rewards that are preceded by the cue and, by definition, is updated only at the time of reward (“whenever a reward occurs, check whether it was preceded by the cue”). Therefore, ANCCR predicts that true unlearning occurs only when rewards are experienced without the associated cue preceding them (they may be fully uncued like in contingency degradation experiments, or be preceded by another cue, i.e., part of another trial type). This behavioral paradigm, which we term retrospective extinction, degrades the retrospective P(cue←reward) association and induces true memory erasure (**Fig. 1C**, retrospective extinction).

In sum, in ANCCR, the primary stored memory is a single retrospective memory indexed by the physical context (**Supplementary Note 2**), rather than multiple prospective memories indexed by hidden contexts. This distinction is the central conceptual difference between ANCCR and latent-cause RPE frameworks. Distinguishing between these frameworks is crucial. Beyond advancing our understanding of learning, establishing conditions for memory erasure could inform the design of more durable treatments for disorders in which maladaptive memories drive persistent harmful behavior.

## Results

### Retrospective extinction erases the original memory

We began by testing whether retrospective extinction can, in fact, erase a cue-reward memory. We assessed memory erasure by comparing learning speeds of behavior and dopamine cue response during reacquisition with those during initial acquisition, and also by measuring spontaneous recovery of behavior during extinction. Rapid reacquisition indicates preservation of the cue-reward memory. In contrast, if memory is lost, the speed of reacquisition in the animals that do reacquire should be comparable to initial acquisition; further, spontaneous recovery should be minimal.

We employed Pavlovian conditioning in head-fixed mice that randomly interleaves two distinct tones (CS-A, CS-B). During the *Acquisition* phase, both cues predicted sucrose reward with 50% probability, denoted [A→^50%^sucrose, B→^50%^sucrose], and conditioned behavior was measured using anticipatory licking (**Fig. 1D-F**). Thereafter, mice were divided into two groups to perform either standard or retrospective extinction of CS-A, while performing standard extinction of CS-B in both groups. CS-B trials therefore provide a within-animal control for comparing memory strength to CS-A (see **fig. S1** for detailed rationale of the experimental design). To achieve such cue-specific extinction, in the phase after *Acquisition,* “*CS-A extinction*”, Group1 (G1, standard extinction, purple), received only CS-A followed by reward omission [A→omit]; here ANCCR predicts that the lack of rewards should prevent erasure of the CS-A←reward association. In contrast, Group2 (G2, retrospective extinction, green), experienced CS-A omission trials randomly interleaved with CS-B trials [A→omit, B→^50%^sucrose]; because animals continue to receive rewards in CS-B trials, but not in CS-A trials, ANCCR predicts that CS-A←reward association should be degraded. For both groups, we continued the protocol until responding to *CS-A* was eliminated. Thereafter, CS-B underwent standard extinction (“*CS-B extinction*”) in which both CS-A and CS-B were unrewarded [A→omit, B→omit]. Memory was then tested in the *Reacquisition* phase, which resumed the original rule [A→^50%^sucrose, B→^50%^sucrose] (**Fig. 1D**), by determining whether behavioral reacquisition was faster than the original acquisition (**Fig. 1F**). In short, this paradigm selectively degrades the retrospective CS-A←reward association in G2, but not in G1 (**Fig. 1E**).

We first focused on CS-A trials. As predicted, G1 animals reacquired anticipatory licking to CS-A within a few trials, learning significantly faster than during initial acquisition, and demonstrating memory preservation under standard extinction. In contrast, G2 animals reacquired CS-A behavior much more slowly than G1, at a rate matching their own acquisition, and consistent with erasure of original memory (**Fig. 1G-L**). Interestingly, ∼40% of animals in G2 (compared to 0% in G1) never reacquired conditioned behavior, demonstrating the power of retrospective extinction (**Fig. 1K**, **fig. S2A,B**). G2 also showed reduced spontaneous recovery of licking during extinction sessions, measured roughly two weeks after CS-A responding had ceased on average (**Fig. 1M,N**), further indicating memory erasure.

Simultaneous dopamine release measurement in a subset of animals revealed parallel group differences (**Fig. 1O-Q**): rapid reacquisition of dopamine responses to CS-A in G1, but reacquisition as slow as acquisition in G2. Critically, this held even in the ∼40% of animals that never reacquired anticipatory licking (non-learners, **Fig. 1K**; 50% of dopamine-recorded G2 animals, **fig. S2N-Q**). Cue-evoked dopamine responses track the learned strength of the cue-reward association while behavior may require a downstream threshold crossing(*45*). Therefore, the dopamine response provides the most direct evidence that the underlying cue-reward memory was degraded rather than merely suppressed: had retrospective extinction left the memory intact and only raised the threshold for behavioral expression, dopamine responses should have still been reacquired faster in G2, similar to G1. They were not.

The dissociation between dopaminergic and behavioral reacquisition in G2 indicates a second, separable effect. Previous work has shown that animals can develop cue-evoked dopamine responses over training without expressing conditioned behavior(*45*), suggesting that cue-induced behavior expression requires threshold crossing of cue-induced dopamine (“is the evidence sufficient to believe that the cue causes the reward?”). Retrospective extinction therefore appears to have additionally raised this threshold, which may also account for the modest slowing of CS-B behavioral reacquisition in G2 relative to G1 (**fig. S2I-K**) and for the similarly rapid reacquisition of dopamine responses to CS-B across groups (**fig. S2L,M**). This expression effect is layered on top of memory erasure rather than an alternative to it: CS-B reacquisition in G2 remained substantially faster than both CS-A reacquisition and initial acquisition (**fig. S2I,J**), confirming that the CS-B memory was preserved despite the reduction in behavioral expression.

We next asked whether the group difference in reacquisition could be attributed to other differences between groups. G1 and G2 did not differ in anticipatory behavior immediately before reacquisition (**fig. S2C**) or before spontaneous recovery (**fig. S2D**). Reacquisition was consistently faster in G1 than G2 regardless of the number of matching extinction sessions (**fig. S2F-H**). G2 showed slower CS-A extinction than G1 (**fig. S3A-D**), consistent with a protective effect of cross-cue generalization (**Supplementary Note 3**) (**fig. S3E-H**). However, extinction rate did not correlate with reacquisition rate within either group (**fig. S2E**). Therefore, reacquisition results are not explained by these differences. Additionally, for *CS-B extinction*, ANCCR predicts slower extinction in G2 than G1, because the reward rate remained elevated during the *CS-A extinction* phase in G2 but was degraded to zero in G1 (**fig. S4A,B**). Behavior and dopamine responses supported this prediction (**fig. S4C-H**). ANCCR also predicts faster behavioral extinction of CS-B than CS-A within G1, which we likewise observed (**fig. S4I-O**). Together, these analyses confirm that the impact of retrospective extinction on memory erasure cannot be explained by differences in baseline behavior or extinction dynamics.

### Suppression of dopamine response at reward during retrospective extinction prevents memory erasure

Because retrospective extinction induced unlearning as predicted by ANCCR, we next tested ANCCR’s additional postulate that dopamine gates retrospective learning at rewards. In ANCCR, memory erasure is counterintuitively hypothesized to occur not due to reward omission, but instead by receipt of rewards themselves, specifically those not preceded by CS-A. To test whether unlearning of CS-A←reward memory in G2 depends on dopamine gating at reward on CS-B trials, we optogenetically suppressed dopamine neuron activity at reward, while leaving dopamine transients unmanipulated during interleaved CS-A→omission trials (**Fig. 2A**, G2-opto-Exp). In the experimental group, an inhibitory opsin (stGtACR2) was expressed in VTA dopamine neurons and laser (473nm, 8-11mW) was delivered for 5 s following reward delivery on CS-B trials during *CS-A extinction*. The 5 s window was selected to ensure complete suppression of reward response (**Fig. 2C**). Control animals received the same laser stimulation but without opsin expression (G2-opto-Ctrl). Thus, G2-opto-Ctrl was predicted to replicate the CS-A←reward memory erasure as in the original G2, whereas G2-opto-Exp should block this erasure, making the result more similar to the G1 (**Fig. 2B**). Reacquisition to CS-B was expected to remain rapid in both groups (see **fig. S5** to find CS-B responses).

**Fig. 2.**
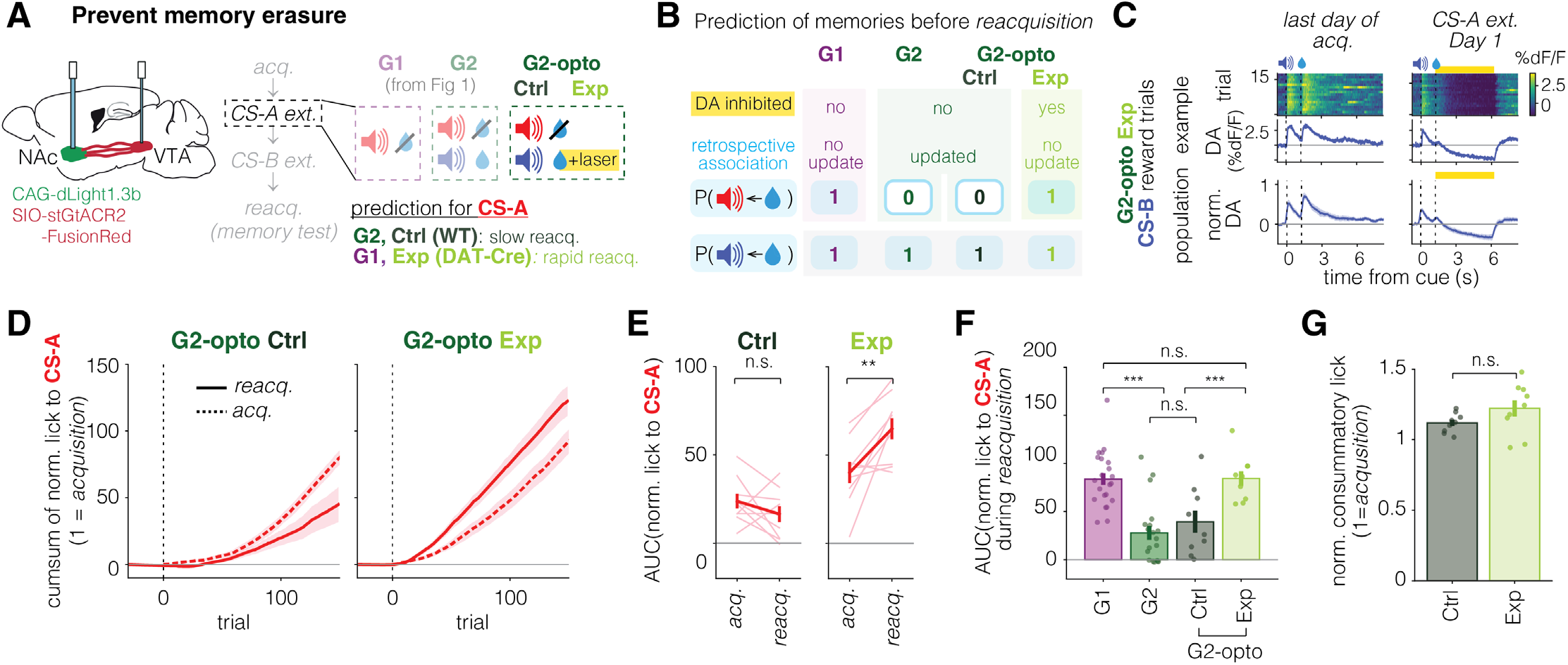
Dopamine response at reward gates memory erasure via retrospective extinction. **(A)** Experimental schematic. In the same CS-A extinction paradigm used for G2, dopamine neurons in VTA were optogenetically suppressed for 5 s following reward delivery in CS-B trials in experimental group (Exp). The control group (Ctrl) underwent the same procedure without expression of inhibitory opsin. **(B)** Prediction. Dopamine is hypothesized to gate updates of retrospective association occurring at the reward. In Exp, suppressing dopamine response during CS-B rewards prevents degradation of CS-A ← reward memory, resulting in fast reacquisition to CS-A (like G1). Ctrl is expected to resemble G2, showing unlearning and slow CS-A reacquisition. **(C)** Example heatmaps and PSTHs of dopamine release (dF/F) in the nucleus accumbens during CS-B rewarded trials from an example Exp animal (top) and group-averaged PSTHs across all Exp animals (bottom). The left column shows data from the last day of acquisition with no laser, and the right column shows the first day of CS-A extinction with laser presented on CS-B reward trials. Yellow bars indicate laser-on periods. Dopamine response to reward after CS-B was strongly suppressed during the CS-A extinction phase compared to the last day of acquisition. **(D)** Cumulative sum of normalized lick to CS-A across trials during acquisition (dotted) and reacquisition (solid). In the Ctrl group (left), reacquisition is slow, resembling the pattern observed in G2 without optogenetic manipulation (see Fig. 1J). In contrast, the Exp group (right) shows rapid reacquisition. Lines represent mean across animals and shadings indicate SEM. **(E)** Comparison of AUC of normalized lick between acquisition and reacquisition for CS-A. Light lines represent individual animals. **(F)** Comparison of AUC of normalized lick to CS-A during reacquisition to the original groups 1 and 2 without optogenetic manipulation (Fig. 1), showing similarity of Ctrl to G2, and of Exp to G1. Error bars represent SEM across animals. G1 and G2 results are from the same animals shown in Fig. 1. **(G)** Mean normalized consummatory lick during CS-A extinction, showing no difference between groups. Error bars represent SEM across animals and 1 indicates the average consummatory lick over last 3 days of acquisition.

Simultaneous recording of dopamine release in the nucleus accumbens confirmed effective suppression of reward-evoked dopamine, with net responses reduced below baseline (**Fig. 2C, fig. S5L**). In ANCCR, such below-baseline suppression at the time of reward reduces behavior to both cues by lowering estimated reward magnitude (as observed, **fig. S5M**) without degrading retrospective association. Consistent with prevention of memory erasure, the experimental group reacquired CS-A behavior rapidly, whereas the control group showed slower reacquisition than acquisition, similar to the original G2 (**Fig. 2D-F, fig. S5B,C**). Likewise, in the experimental group, reacquisition speed of dopaminergic response to CS-A was as rapid as that to CS-B, further supporting a blockade of retrospective learning (**fig. S5D-I**). Notably, consummatory licking was not affected during optogenetic manipulation, indicating that suppression of dopamine response at reward did not impair reward consumption by inducing aversion, but selectively disrupted retrospective learning triggered by rewards (**Fig. 2G**). Importantly, the reacquisition test was performed >6 days after the final optogenetic manipulation, making it unlikely that differences in reacquisition speed between groups can be attributed to artifacts of the optogenetic manipulation. Together, these findings support that behavior extinction in the experimental group did not arise from the degradation of stored cue-reward memory like in the control group or the original G2, but by suppressing behavioral output variables like in G1 (estimated reward magnitude in the experimental group and reward rate in G1). They further support a causal role for dopamine activity at rewards themselves, on trials lacking CS-A, in driving CS-A←reward memory erasure. This gating role of dopamine is consistent with the emerging consensus among alternative dopamine models proposing learning rate control (*42*, *44*, *46*, *47*), and supports the core postulate of reward-triggered retrospective learning in ANCCR.

### Latent cause models do not explain memory erasure by retrospective extinction

Results from G1 and G2 (**Fig. 1**) and from causal dopamine inhibition in G2-opto (**Fig. 2**) are consistent with simulations of ANCCR (**Fig. 3A-C**). In latent cause models, whether the original cue-reward relationship is erased or maintained depends on which latent context is inferred during extinction. This inference is implemented in many ways by distinct versions of latent cause models (**fig. S6A**). For example, the model by Redish et al. 2007 (*40*) infers a single discrete context on each trial based on the similarity between current observations—such as sensory cues and recent reward history—and the learned context prototypes. In this framework, persistent negative RPEs promote the inference of a new state by increasing sensitivity to the current observations (which likely contain the clues regarding the identity of the new state; **fig. S6B**). By contrast, Bayesian latent cause models (*37*, *38*), formalizing prior ideas (*48–50*), represent context inference probabilistically, maintaining a graded belief over multiple contexts on each trial and updating these beliefs via posterior inference. In this setting, both positive and negative prediction errors can favor inference of a new context by reducing the posterior probability of existing contexts (**fig. S6C-D**). In addition to latent cause models, a recent RPE-based model named “value RNN” (*51*) learns to identify and encode task structure implicitly through the dynamics of a recurrent neural network, and learns the value of cues in different contexts without explicit context inference.

**Fig. 3.**
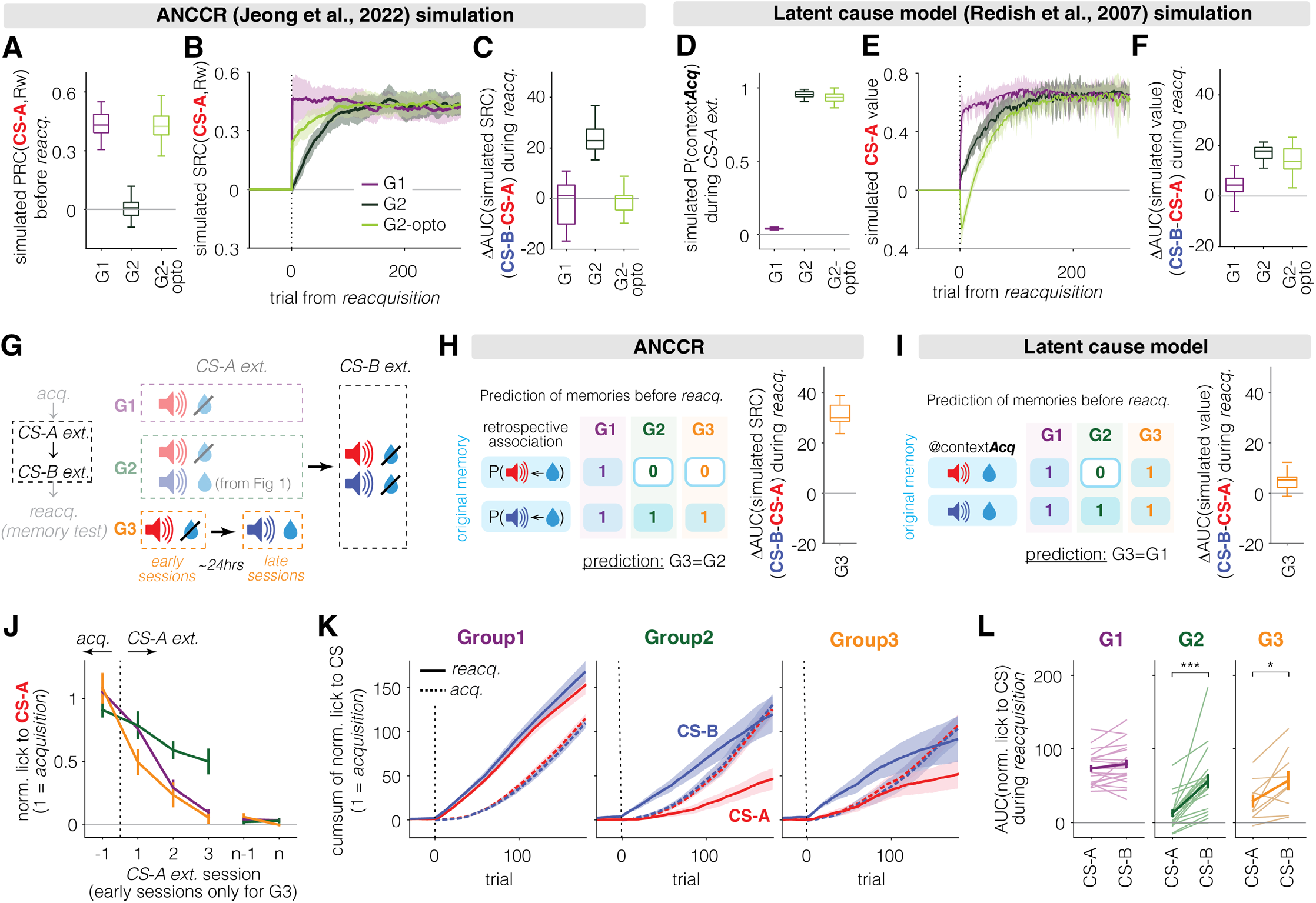
Latent cause model cannot explain slower reacquisition following retrospective extinction. **(A-C)** ANCCR simulations for G1, G2, and G2-opto. **(A)** Simulated retrospective association (predecessor representation contingency, PRC) between CS-A and reward after CS-A extinction. Higher value implies preserved memory, while lower value reflects unlearning. **(B)** Simulated prospective association (successor representation contingency, SRC) between CS-A and reward during reacquisition. It captures the behavioral results. ANCCR predicts that dopamine inhibition at reward following CS-B (G2-opto) would cause G2 to behave like G1, consistent with the empirical results (Fig. 2). The lower rise of SRC (and predicted behavior) in G2-opto is because the previously experienced non-zero reward rate in G2 (which was during *CS-A extinction*) is half that of G1 (which was during *Acquisition*). **(C)** Difference in AUC of simulated SRC between CS-A and CS-B during reacquisition. **(D-F)** Latent cause model (Redish et al., 2007; see **fig. S6-7** for other latent cause models) simulations for G1, G2 and G2-opto. **(D)** Simulated probability of inferring the acquisition context (contextAcq) during CS-A extinction. Closer to zero indicates switching to a new context (memory preserved), while higher values indicate inference of contextAcq (unlearning). In box plot, the central line indicates the median of iterations; box edges mark the 25^th^ and 75^th^ percentiles; whiskers extend to the most extreme data points within 1.5× the interquartile range. **(E)** Simulated CS-A value across trials during reacquisition can capture behavioral results of G1 and G2. However, latent cause model predicts that dopamine inhibition at reward following CS-B (G2-opto) would not speed up behavior in G2, which contradicts the empirical results (Fig. 2). Lines indicate mean and shadings represent STD across iterations. Negative CS-A value early in *Reacquisition* is due to the negative RPE from dopamine inhibition in G2-opto. **(F)** Difference in AUC of simulated values between CS-A and CS-B during reacquisition, reflecting the predicted cue-dependent difference in reacquisition speed. **(G)** Task schematic for CS-A extinction and CS-B extinction. Group3 (G3) is similar to G2, but CS-A and CS-B trials were separated into distinct sessions across days during CS-A extinction. CS-B extinction, consisting of omission trials for both CS-A and CS-B, was identical across groups. **(H)** ANCCR predicts G3 would behave like G2, since CS-B rewarded trials degrade the retrospective association between CS-A and reward, leading to unlearning, even though CS-B reward trials happen more than two days after behavior extinction for CS-A. **(I)** Latent cause model predicts that G3 would behave like G1, since animals infer a new context during CS-A omission trials (early sessions of CS-A extinction), preserving original memory in contextAcq. **(J-L)** Behavioral results support ANCCR. G1 and G2 results are from the same animals shown in Fig. 1. **(J)** Average lick to CS-A in each session during CS-A extinction phase. Day n indicates the last day of CS-A extinction. For G3, only early sessions, those identical to CS-A extinction phase in G1, were included in analysis. **(K)** Group-averaged cumulative sum of normalized lick to CS-A (red) and CS-B (blue) during acquisition (dotted) and reacquisition (solid). **(L)** AUC of normalized lick to CS-A and CS-B during reacquisition. CS-B reacquisition is faster than acquisition in all groups, but CS-A reacquisition is faster only in G1 and not in G2 or G3.

Can any of these models capture the above experimental results anticipated by retrospective learning? Bayesian versions of the latent cause model (*37*, *38*) and value RNN (*51*) failed to account for the results from G1 and G2 (**fig. S6N-U, 7**), and thus we did not further evaluate their predictions on G2-opto. By contrast, the Redish et al. model (*40*) was compatible with the results from G1 and G2. This model correctly predicted that CS-A memory will be lost in G2 but not in G1 (**Fig. 3D-F, fig. S6E-J**). In this model, the presence of CS-B trials during the *CS-A extinction* in G2—resembling *Acquisition*—increases the probability of inferring the acquisition context (contextAcq) instead of the extinction context (contextExt). In G1, where extinction occurs without CS-B trials, the model infers contextExt, preserving memory. This difference in context inference produces memory erasure in G2 but not in G1, matching behavioral outcomes (**Fig. 3D-F, fig. S6E-J**). However, this model fails to explain the results from G2-opto (**Fig. 3D-F**, **fig. S6M**). Nevertheless, because this discrepancy could stem from implementation details rather than core model principles (**Supplementary Note 4**), we next designed experiments to directly discriminate ANCCR from the core computations underlying all versions of latent cause models.

The core tenet of all latent cause models is that the hidden context inferred *at the time of behavior extinction* determines whether to maintain or degrade memory. In contrast, degradation of memory occurs *at the time of rewards* in ANCCR. To fully distinguish between these update principles for ANCCR and all latent cause models, we designed Group 3 (G3). In G3, *CS-A extinction* phase was temporally separated across sessions: the first 5 daily sessions (‘early sessions’) consisted solely of CS-A omission trials (like G1) [A→omit], whereas the subsequent daily sessions (‘late sessions’) included only CS-B trials with 50% reward probability [B→^50%^sucrose] (**Fig. 3G**). Latent cause models predict that G3 will behave like G1, because early CS-A-only extinction sessions, which are identical to G1, should trigger context switching and preserve CS-A memory (**Fig. 3I, fig. S8A**). Because G3 undergoes the exact same CS-A extinction as G1 during early sessions, subsequent late sessions of CS-B trials should not thereafter degrade the original CS-A-reward memory. Conversely, ANCCR predicts that subsequent rewarded CS-B trials provide opportunities for retrospective extinction of the CS-A←reward, causing memory erasure as in G2 (**Fig. 3H, fig. S8B**). In sum, latent cause models predict that G3 will behave like G1 while ANCCR instead predicts that G3 will behave like G2. Thus, this design discriminates the theoretical frameworks by testing whether rewards delivered >2 days after complete behavior extinction of CS-A (**Fig. 3J**) retrospectively erase the preserved CS-A memory. This is therefore a test of the central principle of reward-triggered retrospective learning.

Behavioral results strongly favor ANCCR over latent cause models. Within each group, we compared CS-A and CS-B reacquisition speeds, using CS-B as an internal control since it underwent identical standard extinction across all groups and should show uniformly rapid reacquisition under both theoretical frameworks (**Fig. 3K,L**). We compared behavioral results from G3 to those from G1 and G2. G1 animals reacquired both cues rapidly with no speed difference, consistent with preserved memory for both cue-reward associations. In contrast, both G2 and G3 exhibited selective impairment. CS-B reacquisition remained rapid, whereas CS-A reacquisition was significantly slower. Accordingly, the difference between CS-A and CS-B reacquisition was significantly larger in G2 and G3 than in G1 (see **Supplementary Table 2** for stats). Importantly, no cue differences were found during initial acquisition across any group (**fig. S8C**), confirming that reacquisition differences are due to extinction manipulations instead of variations in baseline cue salience. Together, these results contradict the core tenet of latent-cause models that persistence of the original memory after extinction is determined by the latent context inferred at the time of extinction. Instead, they support the ANCCR prediction that rewards, even if experienced more than 2 days after behavioral extinction, can retrospectively erase a preserved cue–reward memory.

We next tested whether uncued rewards induce retrospective update of all reward memories in a non-cue-selective manner. To this end, we next tested Group4 (G4), in which the second phase of CS-A extinction (like in G3) included uncued rewards instead of CS-B trials. According to ANCCR, this manipulation should non-selectively degrade all reward-associated memories (**fig. S8D,E**). Consistent with this, G4 animals showed slow reacquisition for both CS-A and CS-B on average, supporting ANCCR’s core idea that reward triggers unlearning (**fig. S8F-K**). G4, however, showed greater individual variability than G2 or G3 (**fig. S8J,K**). As mentioned above, ANCCR also assumes that memories are indexed by contexts. Critically, however, these contexts are assumed to be physical contexts instead of latent contexts inferred mostly based on changes in strength of associations. Therefore, the increased individual variability in G4 likely reflects whether animals treated the reward-only phase as occurring in the same physical context as other phases with auditory cues, or in a novel context. Because a physical context is nothing more than a set of neutral cues, the absence of auditory cues during the reward-only sessions may create ambiguity in inference of the physical context (discussed later in **fig. S14** and **Supplementary Note 2**). Nevertheless, animals with slow CS-A reacquisition also showed slow CS-B reacquisition, demonstrating that uncued rewards degraded all reward-associated memories in these animals, exactly as predicted by ANCCR. Physical context inference therefore constrains retrospective learning: in a design where animals experienced CS-A trials alone, subsequent CS-B reward trials did not degrade CS-A memory, consistent with neutral cues defining physical contexts (**fig. S9A-C**). In contrast, delivering uncued rewards during inter-trial interval of standard extinction, with rewards occurring in the same physical context, induced degradation of memory (**fig. S9D-F**). Together, these results demonstrate that retrospective memory updating requires rewards to be experienced in the same physical context as in acquisition, while remaining unassociated with the original cue (either signaled by another cue as in **Figs. 1-3** or occur without any cue as in **Fig. 4, fig. S9D-F and 10**). This interpretation may also explain why CS-A memory degradation was slightly weaker in G3 than G2 (**Fig. 3K,L**). In G3, CS-B-reward trials during *CS-A extinction* occurred after CS-A sessions and may have increased the likelihood of CS-B trials being assigned to a distinct physical context. Additionally, learning non-associative aspects of the task, such as efficient reward collection in freely moving animals (**fig. S9D-J**), can obscure comparison of reacquisition speed to acquisition to assess memory erasure. Collectively, these results suggest that physical context and non-associative task learning should be carefully considered when designing experiments to evaluate memory erasure via retrospective extinction.

**Fig. 4.**
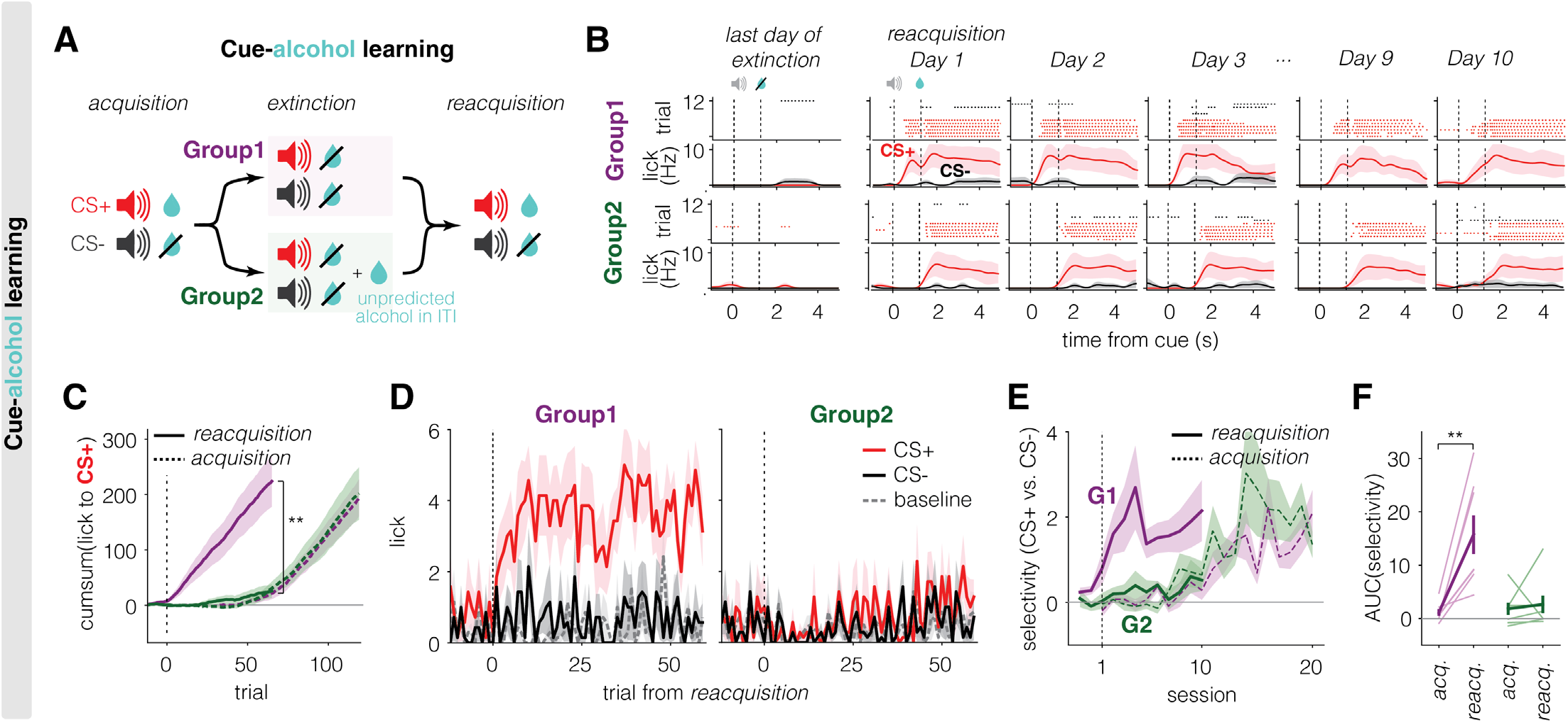
Retrospective extinction erases memory in cue-alcohol learning. **(A)** Schematic of cue-alcohol learning. Animals were trained to associate CS+ with alcohol and CS-with no alcohol. During extinction, alcohol following CS+ was omitted without (G1; standard extinction) or with (G2; retrospective extinction) additional uncued alcohol during ITI. This paradigm predicts that the original memory is preserved in G1, while unlearned in G2. **(B)** Example lick raster plots (top row) and PSTHs (bottom row) for a representative mouse for each group (top, G1; bottom, G2). The first column shows data from the last day (Day -1) of extinction, and the remaining columns show reacquisition. G1 reacquires anticipatory lick for CS+ (red) immediately on Day 1, while G2 did not show anticipatory lick until Day 10. Note that lick to CS-(black) remained for both groups throughout days. In PSTHs, lines indicate mean across trials and shadings represent SEM. **(C)** Cumulative sum of lick to CS+ across trials during acquisition (dotted) and reacquisition (solid). Trials <0 in reacquisition indicate extinction trials. G1 shows rapid reacquisition, while reacquisition is as slow as sacquisition in G2. **(D)** Group-averaged lick count during a 1.25 s window after (red, CS+; black, CS-) or before (dashed gray, baseline) cue onset across trials. Trial zero is the first trial of reacquisition. G1, but not G2, rapidly reacquires anticipatory lick to CS+. Lines indicate mean across animals and shadings represent SEM. **(E)** Group-averaged selectivity (CS+ vs. CS-) based on anticipatory lick during acquisition (dotted) and reacquisition (solid). G1, but not G2, shows faster reacquisition than acquisition. Lines indicate mean across animals and shadings represent SEM. **(F)** AUC of selectivity over first 10 days of acquisition or reacquisition. Light lines represent individual animals. Error bars indicate SEM across animals.

Despite these constraints on retrospective learning, experiments in freely moving rats confirmed that retrospective degradation is reward identity specific and persistent. When different reward types (sucrose vs. food pellet) were associated with different cues, retrospective degradation selectively impaired recovery only for the cue associated with the degraded reward identity (**fig. S10A,B**). Moreover, retrospective extinction effects persisted even when followed by standard extinction, with no evidence of behavioral recovery (**fig. S10C-E**)—a persistence not expected of transient behavioral suppression, contrasting earlier pigeon autoshaping results from a similar experiment (*52*, *53*).

Altogether, we demonstrate cue-specific retrospective memory erasure in G2 and G3, where CS-B-reward trials degrade CS-A retrospective association. By contrast, reacquisition rate was similar for both cues when mesolimbic dopamine activity was suppressed at rewards in G2-opto. Collectively, these findings show that retrospective extinction is a robust and effective procedure for memory erasure via reward-driven updates, providing strong empirical support for ANCCR over latent cause models.

### Retrospective extinction produces unlearning in a simplified design across outcome modalities

The two-cue design in Groups 1-3 enabled rigorous within-animal comparisons while maximizing inference of the same physical context to distinguish retrospective learning from latent-cause models. However, a simpler design in which unpredicted rewards are delivered during inter-trial intervals (i.e., contingency degradation; **fig. S9D-F**) may be more relevant for potential clinical applications because uncued rewards are expected to degrade all reward-associated memories, making it a potentially useful strategy for modifying maladaptive memories in clinical settings. We therefore asked whether memory erasure observed in this simpler retrospective extinction paradigm extends beyond sucrose memories to also be relevant for alcohol memories. We used two auditory cues: a conditioned stimulus (CS+) paired with 20% alcohol, and a control cue (CS-) never associated with outcome [CS+→alcohol; CS-→nothing]. During extinction, G1 underwent standard extinction with omission of outcome after CS+ [CS+→omit; CS-→nothing], whereas G2 experienced retrospective extinction, receiving unpredicted outcomes during inter-trial intervals to trigger retrospective degradation of cue-outcome association [CS+→omit; CS-→nothing; ITI→alcohol] (**Fig. 4A**). We included CS-trials to account for changes in licking that are not specific to CS+, particularly any potential increase induced by uncued alcohol deliveries during the ITI. Retrospective extinction remained highly effective at erasing memory in this design. G1 animals rapidly reacquired conditioned responding to CS+, learning substantially faster than during initial acquisition. In contrast, G2 animals showed markedly slower reacquisition, similar to their original acquisition rate (**Fig. 4B,C**). This pattern remained the same when conditioned responses were quantified using selectivity for CS+ over CS- to control baseline differences across animals (**Fig. 4D-F**). This suggests that retrospective extinction can also induce unlearning for memories associated with drug rewards. Comparable trends were observed in cue-shock learning, although the effect size was smaller (**fig. S11A-H**). Finally, a preliminary human cocaine-use experiment revealed patterns broadly consistent with the possibility that cocaine exposure in treatment settings outside of drug-associated contexts may reduce subsequent self-reported cocaine use (**fig. S11I**). Altogether, these preliminary experiments suggest translational potential for retrospective extinction and motivate the development of rigorously controlled clinical trials in the future.

### Mesolimbic dopamine signaling in contingency degradation and extinction is consistent with ANCCR

Our findings further shed light on ongoing debates regarding dopamine’s role in learning, contrasting prospective reward prediction error (RPE)-based frameworks (including latent cause models) and retrospective learning models such as ANCCR. Here, we focus on Qian et al. (*54*) as a representative example of this broader debate, because their task was specifically designed to test between prospective and retrospective computations during contingency degradation. Qian et al. (*54*) reported that contingency degradation due to uncued rewards is stronger than that due to cued rewards, arguing for prospective computations over ANCCR’s retrospective reward-driven updating (**fig. S12I**).

Though the task design is elegant, this interpretation is premature as Qian et al. do not consider the possibility of cross-cue generalization (**Supplementary Note 3**). When multiple cues precede the same reward, the cues can acquire a shared representation (cross-cue generalization or “acquired equivalence”) (*55–57*). Cross-cue generalization can protect an established cue–reward association from degradation, because when every reward remains preceded by some cue (the “cued-reward” condition), a generalized cue←reward association persists and slows degradation of the original cue. In contrast, uncued rewards degrade both cue-specific and generalized associations, accelerating contingency degradation (**fig. S12J**). Incorporating cross-cue generalization into ANCCR reconciles the findings of Qian et al. (*54*). Evidence for such generalization is present in their own data—specifically, the rapid acquisition of responses to the second cue—and also in our experiments, where it manifests as slower extinction of CS-A behavior and dopamine responses in G2 relative to G1 (**fig. S3, fig. S12A–C**). Nevertheless, animals in G2 ultimately overcome this generalization, extinguishing conditioned behavior to CS-A while maintaining it to CS-B, producing distinct cue-specific reacquisition rates. Thus, cross-cue generalization transiently influences extinction but does not prevent the cue-specific memory erasure predicted by ANCCR.

Finally, RPE-based models including Qian et al. advocated the use of an “unbiased” value RNN to infer latent states without hand-crafted features. In our dataset, however, the same value-RNN framework fails to capture the CS-A reacquisition-rate difference between G1 and G2 (**fig. S7F–H**) and their extinction-rate difference (**fig. S12F–H**), although incorporating cross-cue generalization could recover the latter effect. These results demonstrate that neither ANCCR nor RPE-based models naturally account for known cognitive processes such as cross-cue generalization (*55–57*). Once cross-cue generalization is incorporated, the set of results presented here and in Qian et al. is better explained by retrospective learning.

Additionally, our framework addresses recent counterintuitive findings by Burwell et al. (*58*) that elimination of dopamine omission dip accelerates extinction, contrary to RPE accounts (**fig. S12K**). Classical or latent cause models using RPE predict that large omission-evoked dopamine dips hasten extinction by driving either value reduction or context switching (**fig. S12L**, left/middle). In ANCCR, positive dopamine transients promote learning of an event’s cause, whereas negative dopamine transients suppress it. Therefore, ANCCR instead predicts that the omission dip suppresses formation of a cue←omission (i.e., cue←frustration) association, acting as a compensatory brake that slows behavioral extinction. Artificially attenuating this brake therefore accelerates behavioral extinction, aligning with Burwell et al.’s observations (**fig. S12L**, right; **fig. S12M**). A fundamental prediction of this account is that behavioral extinction can reflect two separable mechanisms: cue←reward erasure is driven by rewards not preceded by the original cue, whereas cue←omission learning can be driven by omitted rewards preceded by the original cue if dopamine dips to omissions are eliminated. This dual-mechanism account is consistent with Burwell et al.’s finding that their perturbation accelerated behavioral extinction without detectably accelerating extinction of the acquired cue-evoked dopamine response. This account is also consistent with Mwenda et al.’s(*59*) demonstration that enhancing dopamine dips during unrewarded actions counterintuitively enhances natural and punishment-resistant reward seeking.

Further, we observed that G2 showed slower extinction despite a persistent dopamine omission dip (**fig. S12D,E**). In addition, larger omission dips were associated with slower extinction across individual animals (**fig. S12N**). Moreover, animals with smaller dopamine dips showed weaker asymptotic behavior after reacquisition, consistent with competition between preserved cue←reward memory and a newly formed cue←omision memory (**fig. S12O**). Together, these results support ANCCR’s interpretation that dopamine dips act as a protective mechanism, preventing formation of conflicting or excessive associations (**fig. S12P**). To summarize, within ANCCR, dopamine release at reward acts as a gate that determines when retrospective cue←reward associations are updated, whereas omission-evoked dopamine dips serve as a protective brake that suppresses formation of competing cue←omission memories. This contrasts with RPE-based views in which both reward and omission signals drive value updating or context switching. Burwell et al.’s findings challenge RPE and latent cause accounts in which omission-evoked dopamine dips serves as necessary teaching signals for extinction of cue-reward memory, and are instead consistent with ANCCR’s proposal that cue←omission learning and cue←reward erasure are distinct computational processes. Overall, behavior and dopamine dynamics during and after extinction are best understood within a retrospective learning framework (**Supplementary Note 5**).

### Orbitofrontal cortex is a potential therapeutic target to prevent relapse following standard extinction

Although retrospective extinction provides a robust framework for memory erasure in experimental settings, it is constrained by a central limitation: its effectiveness depends on exposure to the outcome. In clinical contexts, however, there can be ethical and safety considerations that preclude re-exposing individuals to traumatic events or addictive substances in attempts to erase maladaptive associations. This practical barrier motivates the need for alternative approaches capable of limiting relapse without requiring outcome re-exposure as during retrospective extinction. In other words, this motivates the need for therapeutic approaches that limit reacquisition of conditioned behavior even after prospective extinction.

To address this, we investigated whether broad activity disruption in line with potential human neuromodulation therapies could prevent relapse. Specifically, we focused on the orbitofrontal cortex (OFC), a region driving maladaptive persistence of behavior (*60*, *61*), particularly in addiction (*62–66*). As OFC is functionally heterogeneous along its medial-lateral axis (*67–69*), we focused on the ventral/medial OFC because a neuronal subpopulation in the ventral/medial OFC potentially encodes retrospective associations in its cue response (*43*, *70*), and encodes reward salience in its reward response (*71*). To test whether suppressing ventral/medial OFC during either of these periods can prevent rapid reacquisition, we trained animals with *Acquisition* of a single cue-reward association followed by standard prospective *Extinction* and then selectively inhibited OFC during *Reacquisition* either in the cue-reward period or a 5 s post-reward period (**Fig. 5A**). Optogenetic inhibition of OFC during the reward period markedly slowed reacquisition, with behavioral suppression persisting until the intervention ended (**Fig. 5C,D**; **fig. S13L**). Importantly, consummatory licking was unaffected during laser stimulation (**fig. S13M**), indicating that slowed reacquisition reflected impaired learning, not reduced motivation or motor capacity. In contrast, suppressing OFC during cue-reward delay did not slow reacquisition (**Extended Data fig. 13E, F**). Instead, the laser itself acted as a novel, reward-predictive cue (**Extended Data fig. 13D-K**), thereby preventing a clean assessment of degraded cue-outcome memory in this condition. In a separate cohort, inhibition of OFC during the outcome period of *Acquisition* slowed initial learning (**fig. S13N**), consistent with the idea that OFC activity during reward delivery contributes to updating reward-related information required for learning.

**Fig. 5.**
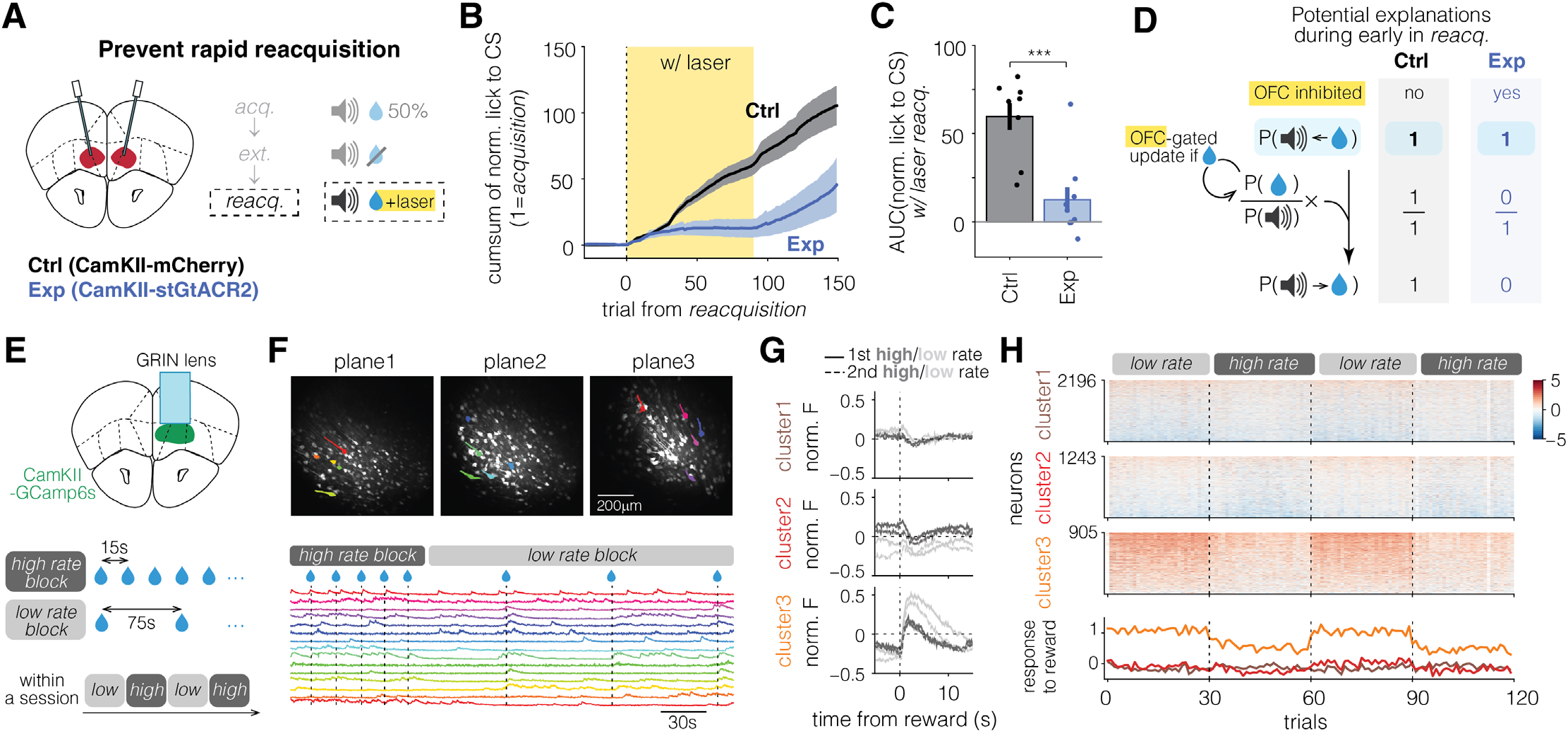
Reacquisition speed can be modulated by manipulating orbitofrontal cortical activity. **(A-D)** Suppressing orbitofrontal cortex (OFC) can prevent rapid reacquisition. **(A)** Experimental schematic. After acquisition of single cue-reward association, standard extinction and reacquisition were followed in a sequence. During reacquisition following standard extinction, OFC CamKIIα expressing neurons were optogenetically suppressed for 5 s following reward delivery in experimental group (Exp). The control group (Ctrl) underwent the same procedure without expression of inhibitory opsin. Note that behavioral outcome (anticipatory licks) is measured during a non-overlapping period with the laser (post reward). **(B)** Group-averaged cumulative sum of normalized lick to CS across trials. Yellow shading marks the laser-on trials. **(C)** AUC of normalized lick to CS during laser-on trials of reacquisition. Learning is suppressed in Exp. **(D)** Potential explanation of the results. If OFC controls the update of P(reward) during reacquisition, its suppression in Exp should block the update and delay reacquisition, while Ctrl should reacquire rapidly. **(E-H)** OFC encodes estimated reward rate. **(E)** Experimental schematic. Uncued rewards were delivered at either short (mean 15 s, high rate block) or long (mean 75s, low rate block) inter-reward intervals, with reward rate switching three times within a session. Activity of OFC CamKIIα-expressing neurons were recorded through a GRIN lens. **(F)** Example fields of view across three simultaneously recorded imaging planes (top) and fluorescence traces of representative neurons (bottom). Vertical dashed lines indicate reward delivery. **(G)** PSTHs of clustered neurons (see Methods) averaged across the last three trials of each block. Each reward rate includes two blocks (1st block: solid, 2nd block: dashed). PSTHs show averaged normalized fluorescence across all neurons within each cluster. Cluster 2 primarily modulates baseline activity with reward rate, whereas cluster 3 modulates reward-evoked phasic response. Note that the higher baseline activity of cluster 2 during high reward rates cannot be trivially explained by slowly decaying reward responses measured by GCaMP6s fluorescence because phasic reward responses are negative in this cluster. **(H)** Phasic reward responses of individual neurons across trials (top) and averaged response across neurons within each cluster (bottom). Phasic response to reward was quantified as the difference in area under curve of fluorescence traces 5 s before and after the first lick following reward delivery.

These experiments were designed purely to test whether OFC is a promising target for neuromodulation therapy and not to test between different theoretical frameworks as explanations of OFC computation. Nevertheless, we tested whether these results are consistent with ANCCR. A slowing of reacquisition in ANCCR could arise through two possible mechanisms during the *Reacquisition* phase, either by suppressing the stored retrospective memory (**fig. S13A, B**), or by impairing rapid reinstatement of the estimated reward rate needed for behavioral expression (**Fig. 5A, B**). Because reinstatement of the estimated reward rate is expected to occur upon reward delivery, whereas retrieval of the stored retrospective memory would be expected during cue presentation, the finding that OFC inactivation during the reward period slowed reacquisition is more consistent with a role in reward-rate reinstatement than in retrospective memory retrieval. More broadly, however, this result does not distinguish ANCCR from other frameworks including prospective learning models, as the result can also be explained by known OFC’s role in encoding of cognitive maps, outcomes, and value (*71–79*), and the encoding of reward salience in ventral/medial OFC (*71*). Although discriminating between different computational accounts will require future experiments specifically designed for that purpose, the present findings demonstrate that transient disruption of OFC activity during outcome processing substantially reduces rapid reacquisition following standard extinction, highlighting the OFC as a promising therapeutic target for limiting relapse.

Having identified OFC as a potential therapeutic target, we next sought to determine what reward-related information is represented during the outcome period, the time window in which OFC inhibition suppressed rapid reacquisition. Because testing such reward rate encoding is difficult during reacquisition itself (e.g., G1 and G2 from Fig. 1 have similar reward rates during reacquisition), we designed an experiment in which animals experienced different (uncued) reward rates. During this condition, we performed two-photon calcium imaging of OFC neurons to directly test whether OFC neurons encode reward rate immediately after reward delivery (i.e., the trial period during which OFC inhibition abolished behavioral reacquisition). Each session contained four alternating blocks of high rate (mean interval of 15 s) or a low rate (mean interval of 75 s) of rewards, with 30 rewards per block (**Fig. 5E**). The experiment was repeated across two sessions with counterbalanced block orders, yielding consistent results across days (**Fig 5G,H** and **fig. S14E-G** from Day 2; Day 1 shown in **fig. S14H-J**). Across three imaging planes in each of four animals, we recorded activity from ∼4,000 OFC neurons (**Fig. 5F**; **fig. S14A**). Licking behavior was comparable across blocks (**fig. S14D**, left), yet many neurons differentiated reward rate either through changes in baseline activity or phasic reward response (**fig. S14D**, right). Spectral clustering (*80*) identified three distinct neuronal subpopulations (**fig. S14B,C**) including one whose phasic reward responses rapidly adjusted to the current reward rate (Cluster3; **Fig. 5G,H**; **fig. S14E-J**). This population may encode a direct estimate of reward rate or, alternatively, a memory of reward rate given reward (i.e., P(reward|reward))—a statistic capturing how likely rewards are to recur once one is encountered. Such a prior would allow animals to rapidly restore their internal estimate of reward rate, and thus behavior, when they return to the reward-rich environment after prolonged reward absence (e.g., extinction→reacquisition; home cage→training). Together with the optogenetic experiments, these results demonstrate that OFC activity during outcome delivery is essential for reinstating extinguished behaviors, suggesting that targeted neuromodulation of OFC activity could curb cue-induced relapse without requiring direct outcome re-exposure during extinction.

## Discussion

Our findings identify a fundamental principle governing whether extinction suppresses behavior versus erases memory. By comparing standard, cue-based (“prospective”) extinction with outcome-based (“retrospective”) extinction, we show that unlearning occurs only when outcomes are experienced without their predictive cues. This retrospective extinction selectively degrades cue←outcome associations and is gated by dopamine release at outcome. Orbitofrontal cortex activity is necessary for rapid reacquisition after prospective extinction. Together, these results unify behavioral, circuit, and computational perspectives to reveal how dopamine-gated retrospective learning enables genuine memory erasure—reconceptualizing unlearning as a process governed by outcome-triggered, rather than cue-driven, updating of associative memories.

Because conditioned behavior can decline for reasons unrelated to memory loss(*12*), a claim of erasure requires convergent evidence. Seven lines of evidence converge here. First, behavioral reacquisition in G2 animals that did reacquire proceeded similarly to acquisition, and ∼40% of animals never reacquired (**Fig. 1G-L**). Second, independent of any behavioral criterion, cue-evoked dopamine responses in G2 were reacquired as slowly as during acquisition, including in the animals that never resumed licking (**Fig. 1O-Q, fig. S2P,Q**). This excludes the possibility that slow behavioral reacquisition reflected only a raised threshold for behavioral expression despite rapid neural reacquisition. Third, even after CS-A responding had ceased for roughly two weeks, G1 continued to show spontaneous recovery, whereas this recovery was largely absent in G2 (**Fig. 1M,N**). Fourth, the effect was specific: CS-B, which underwent identical standard extinction in every group, reacquired rapidly in the same animals (**Fig. 3K,L**), and in rats, degradation was confined to the cue associated with the degraded reward identity (**fig. S10A,B**). Fifth, the same effects were observed even when the degrading rewards occurred in separate sessions more than two days after CS-A responding had ceased (**Fig 3, fig. S8**). Sixth, the effect persisted through subsequent standard extinction without behavioral recovery (**fig. S10C-E**), arguing against mere transient suppression. Seventh, erasure was abolished by suppressing dopamine at reward while leaving reward delivery and every other task variable intact (**Fig. 2**)—establishing that degradation is caused by reward-triggered retrospective updating rather than by reward exposure or differences in reward history between groups. Furthermore, the same pattern was reproduced in a simpler retrospective extinction paradigm using uncued outcomes—for sucrose and alcohol, and, for shock with a smaller but significant effect (**Fig. 4, fig. S9D-F, 11**).

These assays test for faster reacquisition, time-dependent recovery, and cue- and outcome-specific effects. They do not establish that no trace whatsoever remains; a memory inaccessible under all these conditions could in principle be revealed by others. We therefore use “erasure” to denote degradation of the retrospective cue←outcome association to the point where it confers no measurable advantage during reacquisition, supports no spontaneous recovery, and leaves no cue-specific residue—the operational sense in which unlearning has been sought, and hard to find, in prior extinction work.

Within the ANCCR framework, diminished conditioned behavior can arise from degradation of three computational variables: cue←outcome memory, estimated outcome rate, and estimated outcome magnitude (**fig. S15A,B**). Only degradation of the former produces unlearning. Thus, findings traditionally interpreted as unlearning may instead reflect distinct mechanisms. For example, weakened relapse following gradual reduction of outcome frequency (*14*, *15*, *22*) (**fig. S15C,D**) or magnitude (*16*, *81*) (**fig. S15E,F**) can be explained by a lowered prior of outcome rate or magnitude (**Supplementary Note 2**). Specifically, learning that outcome rate is lower or that its magnitude is reduced may be sufficient to suppress behavior even though the original memory of the cue-outcome association is still intact (**fig. S15C-F**). Similarly, optogenetic suppression of dopamine at rewards may have caused a reduction in estimated outcome magnitude and resulted in reduction of conditioned behavior despite intact original memory (*82*, *83*). Additionally, strengthening a competing memory, such as cue-omission association, can also reduce conditioned behavior as observed by Burwell et al. (**fig. S12K-P**).

CS-retrieval extinction, in which the cue is presented by itself (and believed to open a reconsolidation window) before repeated extinction reduces relapse (*17*, *20*, *21*, *23*, *24*) (but see (*26–32*, *35*)). Though not a core prediction of ANCCR, this can be understood as altered inference of a physical context rather than memory degradation. A physical context is nothing more than a set of neutral cues: altering the distribution of neutral cues may lead to inference of a new physical context. Therefore, introducing a temporal gap between the retrieval cue and extinction (often much longer than acquisition inter-cue intervals) and thereby altering the inter-cue intervals expected for a physical context, may cause animals to infer a new context, thereby reducing relapse by limiting access to the original memory (**fig. S16A,B**). This view preserves the role of context-indexed learning and memory like the latent cause theory, though suggesting that statistics of neutral cues, instead of their outcome associations, dominate context inference (**Supplementary Note 2**). Reduced relapse in a paradigm with reversed retrieval and extinction trials (*84*, *85*) and with varied ITI during extinction (*86*, *87*) support this interpretation. By contrast, outcome-retrieval extinction represents a rare case consistent with genuine memory degradation. An uncued outcome triggers retrospective updating that weakens all cue←shock (*18*) or cue←drug (*19*) associations, consistent with the experimentally observed reduction in relapse across all cues linked to that outcome (**fig. S16C-E**). This is also consistent with memory erasure-like phenotypes with contingency degradation in instrumental learning(*25*). Overall, relapses are best understood as shaped by interactions between associative memory persistence and modulation of behavioral output variables. This nuanced view broadens the implications for developing therapeutic strategies.

Beyond clarifying when extinction produces true unlearning, our results also identify a neural mechanism gating this computational outcome. In doing so, they refine current views of mesolimbic dopamine function. Prior work showed that mesolimbic dopamine conveys causal associations (*42*, *88*) but left open whether and how these signals control memory persistence versus erasure. By directly studying standard (prospective) extinction, retrospective extinction, and selective manipulations of dopamine at reward, we find that reward-evoked dopamine transients gate updates of retrospective cue←reward associations. By contrast, our findings, along with those of Burwell et al. (*58*), suggest that omission-evoked dips primarily act as a brake on forming competing cue←omission memories. Thus, both positive and negative dopamine transients participate in the same gating process predicted by ANCCR: positive transients promote learning of an event’s cause, whereas negative transients suppress it. This division of labor explains how dopamine can regulate whether extinction leads to true unlearning or mere behavioral suppression, providing a mechanistic link between computational processes and therapeutic opportunities.

Our findings revise widely held RPE accounts. As explained above, our findings are hard to reconcile with established RPE-based latent cause models or principled RPE-based accounts such as the value-RNN framework (**fig. S7H**). Our results show that rewards, even those delivered two days after behavior extinction, can retrospectively degrade a cue←reward memory (G3 and G4 in **Fig. 3 and fig. S8**). This has similarities to retrospective revaluation (*89*). A well-known example of this is backward blocking. After initially learning CS-A+CS-B→reward, subsequent CS-A→reward training degrades conditioned behavior to CS-B at test (*90*, *91*). Prior RPE based models such as Kalman temporal-difference learning (Kalman-TD) explain this finding (*92*). However, such models cannot explain our G3 and G4 results. This is because Kalman-TD operates by learning cross-covariance between simultaneously experienced cues. In our experiments, on the other hand, the cues were experienced on independent trials and have no cross-covariance. In this sense, our experiment setting is more related to retroactive interference (cues experienced on independent trials) than the commonly studied cue competition setting (cues experienced simultaneously) of retrospective revaluation (*93*). Thus, there is no natural mechanism for models like Kalman TD, applicable to the cue competition setting, to capture the effects seen in G3 and G4. Other models of retrospective revaluation such as the comparator hypothesis do not account for rapid reacquisition after standard extinction (*94*), and as such, also do not apply to our data.

An alternative explanation for the slower reacquisition observed in G2 is the acquisition of an inhibitory association by CS-A, i.e., conditioned inhibition. In principle, slower reacquisition could arise if extinction caused CS-A to acquire an inhibitory association with reward rather than erasing the original positive cue←reward memory. However, several observations argue against this interpretation. First, modified versions of Wagner’s Standard Operating Procedure (SOP) model (*95–97*), a prominent associative account of conditioned inhibition, can under certain parameter regimes, predict slower reacquisition in G2 by allowing inhibitory association between CS-A and reward. However, they cannot account for results from G3 (**fig. S17**). Second, both conditioned behavior in learners (**Fig. 1L**) and cue-evoked dopamine responses (**Fig. 1Q**) were reacquired at a comparable speed to initial acquisition, whereas conditioned inhibitors are expected to show retardation of subsequent excitatory learning (*98*). Notably, even animals that did not recover conditioned behavior (non-learners) exhibited full recovery of cue-evoked dopamine response at the same speed as in acquisition (**fig. S2P,Q**), suggesting that their lack of behavior recovery reflects suppressed behavior expression rather than impaired cue-reward learning. Finally, we observed no evidence of negative dopamine response to CS-A at the start of reacquisition (**Fig. 1P**), whereas conditioned inhibitors have previously been shown to evoke negative dopamine response on average (*99*). Together, these findings argue against conditioned inhibition and instead support retrospective memory erasure.

Our results highlight a key distinction for therapeutic interventions in maladaptive memory-driven conditions such as addiction or PTSD: true degradation of the underlying cue–outcome memory versus suppression of its behavioral expression. Genuine erasure of a retrospective association requires exposure to the outcome itself, as in retrospective extinction. By contrast, although modulation of behavioral variables downstream of associative memory suppresses relapse, relapse risk remains so long as the underlying maladaptive association is preserved. For instance, while drug-free therapies leave drug-associations intact, supervised administration of drugs such as opioid agonists may degrade associations between opioid effects and drug-associated cues experienced in street settings (*100–104*). However, the efficacy of clinical implementations of retrospective extinction through drug exposure depends on overcoming one fundamental limitation. This is that treatment itself introduces novel contextual and social cues that are repeatedly experienced in the treatment setting. These novel cues, once paired with drug administration during treatment, can become potent relapse triggers, simply shifting the drug-related retrospective association from the original street cues to treatment-related cues. Thus, a retrospective association related to drug or trauma can only shift from one cue to another, thereby suggesting that shifting to cues rarely encountered in life might provide the best therapeutic approach to minimize relapse. Several key questions remain untested, including how many outcome exposures are required to achieve memory erasure and how inter-exposure interval influences this process. ANCCR predicts that learning per reward experience is greater when the interval between rewards is longer. We previously found empirical evidence consistent with this prediction during initial acquisition (*45*), but whether the same rule applies during retrospective extinction remains to be empirically tested. Our findings therefore motivate systematic testing of strategies that deliberately shift maladaptive associations onto rarely encountered cues to minimize relapse risk, while carefully evaluating their practicality and safety in real-world clinical settings.

Though behavioral therapies to degrade maladaptive cue←outcome memories require outcome exposure, neuromodulation therapies that modulate behavioral-expression variables such as perceived outcome magnitude or outcome rate may provide an alternate approach. Additionally, it may also be possible to simulate experience of uncued outcomes through neuromodulation. Here we show that inactivation of OFC suppresses rate of reacquisition, potentially through modulating behavioral-expression variables. This raises the possibility that if OFC is suppressed, the drug may still be consumed after prolonged abstinence without inducing drug-seeking driven by cues. Thus, neuromodulation approaches provide complementary strategies to limit relapses when direct outcome re-exposure is not safe or feasible.

## Acknowledgments

We thank M. Tadross, A. D. Redish, P. Dayan, G. Quirk, E. Harkin, B. Sabatini, S. Mihalas, L. Kusmierz, M. Kheirbek, M. Stryker, V. Sohal, K. Moussawi, J. Berke, H. Fields, D. Ron, and members of the Namboodiri laboratory for helpful discussions regarding the manuscript. We thank A. L. Cochran for discussions related to her latent cause model. We used LLMs to aid in editing for clarity. This project was supported by NIH R01MH129582, R01AA029661, R01DA062018, the Scott Alan Myers Endowed Professorship, Alfred P Sloan Fellowship, Pew Biomedical Scholarship, Klingenstein-Simons Fellowship (VMKN), CIHR MOP–36429, PJT–175296 (ML), Klingenstein-Simons Fellowship, The David and Lucile Packard Foundation, and Shurl and Kay Curci Foundation (RC2). We used LLMs to aid in editing for clarity. The authors have no competing interests.

## Author contributions

Conceptualization: HJ, VMKN; Methodology: HJ, LZ, FF, ADS, SMLC, EG, PHJ, ML, RC^2^, VMKN; Investigation: HJ, LZ, FF, JAD, CJ, RC^1^, NB, MZ, ADS, SXW, AS, SMLC, EG, SB; Visualization: HJ, EG, SMLC, VMKN; Funding acquisition: ML, RC^2^, VMKN; Project administration: HJ, VMKN; Supervision: VMKN; Writing – original draft: HJ, VMKN; Writing – review & editing: HJ, LZ, FF, JAD, CJ, RC^1^, NB, MZ, ADS, SW, AS, SMLC, EG, SB, PHJ, ML, RC^2^, VMKN; ^1^Risha Chakraborty; ^2^Ritchie Chen

## Supplementary Notes

### Supplementary Note 1

Here, we simplify the calculation of prospective and retrospective associations by denoting them as P(cue→reward) and P(cue←reward) respectively. This involves two main simplifications. One is that the relevant association is better measured by the prospective and retrospective contingencies: specifically, P(cue→reward)-P(reward) and P(cue←reward)-P(cue) respectively. These conditional minus marginal probabilities measure how much more likely a reward is to follow a cue than its baseline probability, and how much more likely a cue is to precede a reward than its baseline probability. The second simplification is in describing the associations in terms of probability, which assumes a “trial period”, i.e., a time period over which either pairing is calculated. In reality, ANCCR does not make such a trial-based assumption. It is instead based on the computation of event-based successor and predecessor representation contingencies from a continuous timeline of events (successor and predecessor contingencies are related to each other in a form that looks exactly like Bayes’ rule though this relationship is not akin to full Bayesian inference). Because these considerations are not necessary to understand and motivate the experimental designs presented here, they are omitted in the presentation of the theory.

### Supplementary Note 2

ANCCR, like latent cause models, assumes that learned associations are context conditioned. The key difference is that ANCCR conditions on *physical* context, which is defined by sensory cues present in the environment, while latent cause models condition on latent contexts even when physical context remains unchanged. The context conditioning in ANCCR results in some nontrivial predictions. First, we have already shown both in terms of theoretical predictions and empirical observations that the rate of learning of a retrospective A←B association is proportional to the inter-B interval (*45*). Thus, ANCCR predicts that animals store the inter-event intervals for different types of stimuli experienced in a context (e.g., inter-reward interval, inter-CS-A interval, etc.). On transitioning from extinction to reacquisition, the original memory of inter-reward interval during conditioning provides a prior for the rapid restoration of estimated reward rate. Such rapid initialization of reward rate produces rapid reacquisition (during standard extinction when the original retrospective association is intact). Second, related to this, if the reward rate is gradually reduced during conditioning like in the gradual extinction paradigm (**fig. S13C-D**), the reward rate prior will be correspondingly lower during reacquisition and it will appear slower than after non-gradual extinction. Third, context conditioning also has a protective role in memory storage. When context conditioned, a retrospective CS-A←reward association is learned as P(CS-A←reward|context), i.e., it is the conditional probability of cue preceding a reward in that context. Therefore, obtaining the same reward in an entirely new context (e.g., homecage versus experimental chamber) would not degrade a previously learned context-conditioned cue-reward memory. For example, P(CS-A←reward | context_1_) can be maintained while learning P(CS-B←reward | context_2_). This captures the everyday intuition: you can learn that restaurant A in city X and restaurant B in city Y both serve good pizza, without “forgetting” about A when you discover B. Even if B is discovered in the same city X, assuming A keeps serving good pizza, A’s retrospective association will only be driven to zero if you completely stop visiting A, so that rewards in that context are now preceded only by B. In this limiting case, the retrospective association P(A←reward) goes to zero, but the marginal probability P(A) also goes to zero, so the prospective association P(A→reward), defined as the conditional probability of reward given A, can remain high. This is the simplest “retrospective extinction” scenario shown in **fig. S1F-J**. Despite the unlearning of retrospective association though, “behavioral reacquisition” will be rapid once A again precedes rewards, as discussed in **fig. S1F-J**. Therefore, when A entirely stops occurring, the prospective association is effectively “remembered”. This fits the definition of the prospective conditional probability P(A→reward), which, by definition, is conditioned on the occurrence of A; if A no longer occurs, the probability remains unchanged, even if it is accessed via retrospective Bayes’ inference when A reappears.

### Supplementary Note 3

The results from **fig. S3** are consistent with cross-cue generalization providing a protective effect against contingency degradation. In the cue-alcohol experiment, which followed a similar paradigm as cue-sucrose experiment but employed only one CS+ (unlike the two CS+s in the cue-sucrose experiment), retrospective extinction impaired reacquisition, but without slowing extinction, in contrast to the results from Group 2 (**fig. S3E-H**). This pattern suggests that, in the cue-sucrose experiment, cross-cue generalization contributed to slowed extinction in G2. Because both CS-A and CS-B were rewarded during acquisition, continued delivery of rewards with CS-B during retrospective extinction in G2 maintained a generalized cue-reward association, which slowed extinction of CS-A. Cross-cue generalization is a well-established phenomenon in associative learning (*55–57*), but for simplicity, many formal models including ANCCR and TDRPE have not yet explicitly modeled it, leaving its precise functional effects empirically uncertain. In contrast, as CS-A extinction in G1 was driven primarily by a decrease in estimated reward rate, it is less affected by cue generalization. This interpretation is supported by the absence of extinction-speed difference in the cue-alcohol experiment, where only a single cue was rewarded. Thus, the observed difference in CS-A extinction speed between groups 1 and 2 likely reflects cross-cue generalization.

### Supplementary Note 4

We did not simulate the value RNN model (*51*) or latent cause models (*37*, *38*) other than Redish et al. (*40*) on the dopamine optogenetic inhibition experiment in **Fig. 2** because they did not capture the behavioral results from groups 1 and 2 in **Fig. 1** (**fig. S6,7**). The Redish et al 2007 latent cause model, while capturing the behavioral difference between groups 1 and 2, does not capture the results of the optogenetic inhibition experiment for Group 2 in **Fig. 2**. This is because the artificially induced negative RPE at reward delivery on CS-B trials makes the model attempt to improve its hidden context inference. This is achieved by making the model more sensitive to the external inputs, based on the reasoning that if the current inferred context is wrong, the external inputs likely contain the clues to better context inference. In control animals with no optogenetic inhibition, the model infers that the original acquisition context is active during the CS-A extinction phase. This is because the model identifies the partial similarity between CS-A extinction phase and the acquisition phase in G2 due to the presence of CS-B trials in both phases. In experimental animals, however, the optogenetic inhibition artificially produces negative RPEs even though the external world has not otherwise changed. Although this makes the model more strongly attuned to the sensory inputs, the inferred context still remains the acquisition context, because the sensory evidence itself is unchanged. Further, as the assumed negative RPE induced by the optogenetic inhibition gets stronger, the value of both cues decreases in the acquisition context. Due to these combined effects, subsequent reacquisition becomes slower as the magnitude of simulated negative RPE increases (**fig. S6M**). Collectively, the observed results that optogenetic inhibition of dopamine reward responses on CS-B trials prevents CS-A memory erasure is inconsistent with this model.

Nevertheless, it is possible that this discrepancy is due to an implementation detail in Redish et al.’s model. This is because the intent of the model is to induce state splitting on encountering consistent negative RPEs. However, because of the particular implementation increasing the attention to external cues, it produces exactly the opposite effect as intended when applied to the optogenetic inhibition experiments performed here. Other implementations reducing the width of the radial basis function for stimulus categorization would be more likely to induce state splitting and capture our results. However, Redish et al. caution that this model is highly unstable. We did not therefore simulate this implementation.

### Supplementary Note 5

ANCCR was proposed as an alternative model of associative learning and dopamine signaling. A common criticism is that ANCCR is “complex” (*54*, *115*) whereas reward prediction error (RPE) models are “simple,” because RPE is often introduced didactically as received minus predicted reward (*116*). In practice, however, the didactic RPE (essentially Rescorla–Wagner) cannot account for crucial phenomena such as persistent memory after extinction (*4*, *7*, *11*) or mesolimbic dopamine ramps (*117–122*), leading to the development of latent-cause (*37–40*) and hidden-state RPE models (*123*, *124*). These model families are substantially more complex, and neither family individually captures both persistent memory and ramps. An RPE-based account of both would need to combine their complexities and further exploit flexible choices of state space and reward function (*125–131*). In contrast, ANCCR naturally captures both persistent memory and dopamine ramps. Thus, relative to the full RPE model family, ANCCR is not unusually complex or overparameterized. Notably, Qian et al. misstate in their main text that ANCCR has 12 free parameters instead of the 6 correctly listed in their Methods (*54*). Their value-RNN framework does offer a principled way to discover latent states and eliminates the degrees of freedom afforded by hand-picked assumptions. The experiments here were designed specifically to test hidden-state inference, and the results disprove the model as it has no additional degrees of freedom that could account for the current findings. Overall, ANCCR is simpler than existing RPE-based models while explaining the same key findings.

In addition, the core computations in ANCCR yield several counterintuitive, easily falsifiable predictions. These include: (i) the persistence of memory after standard extinction and its loss after retrospective extinction; (ii) the exact quantitative dependence of cue–reward learning rate on the inter-reward interval (*45*); and (iii) the dependence of mesolimbic dopamine ramps on inter-trial intervals (*117*). Each prediction is empirically supported. Together, these results demonstrate that ANCCR provides a parsimonious account of well-established phenomena in associative learning and mesolimbic dopamine signaling.

## Supplementary Figures

**fig. S1.**
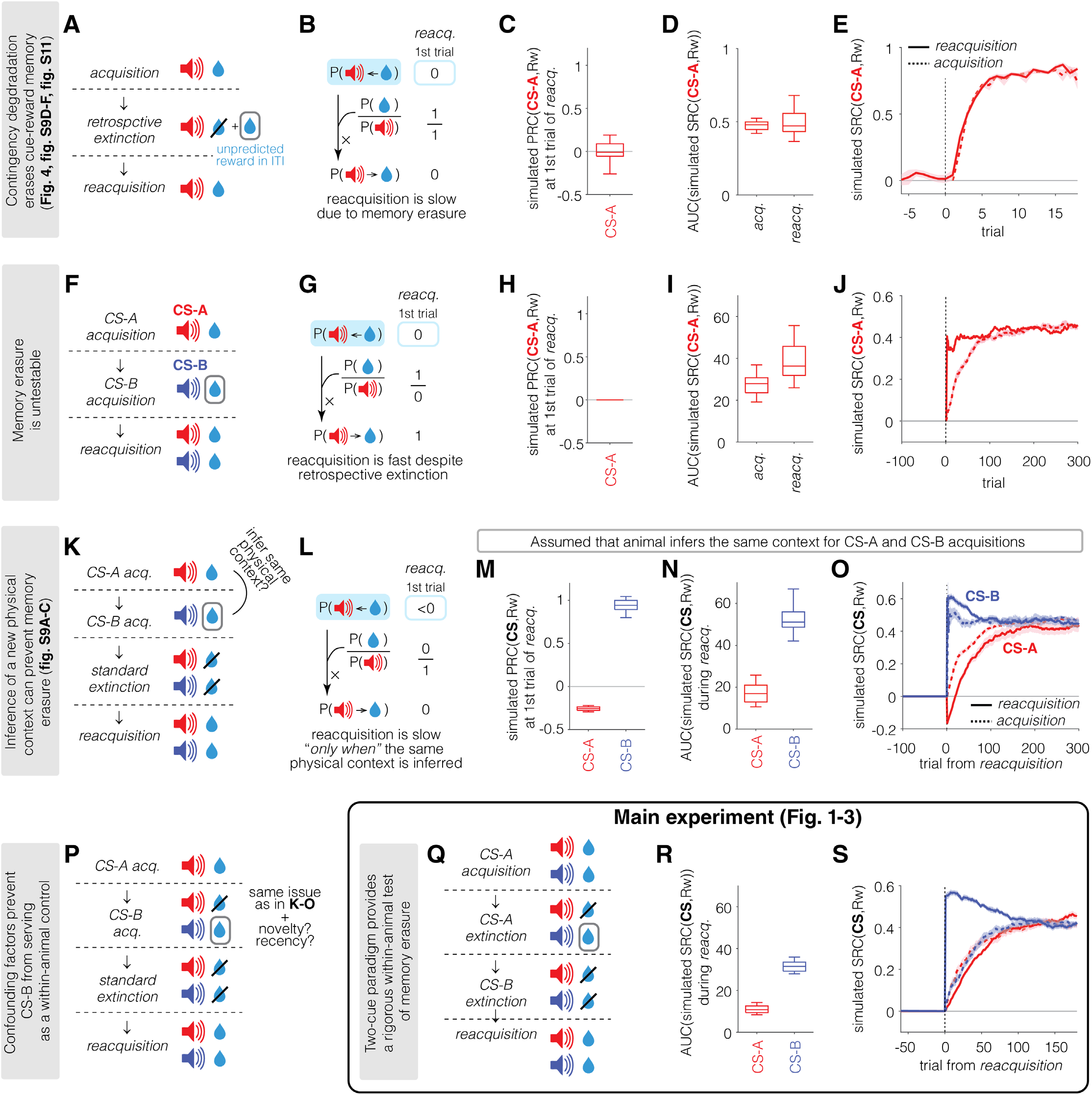
Rationale for the design of the two-cue paradigm to assay memory erasure. Although simpler retrospective extinction paradigms can be envisioned, we reasoned that the two-cue paradigm offers the most rigorous test of memory erasure by enabling within-animal comparisons while avoiding confounds that may blur interpretation in simpler designs. These simulations were included to motivate the experimental design by illustrating why simpler paradigms may not be well suited for assessing changes in cue←reward memory. They are intended as design rationale rather than evidence supporting the ANCCR model. **(A-E) (A)** Contingency degradation paradigm while omitting predicted reward is one of the simplest forms of retrospective extinction. Although this paradigm produced robust memory erasure experimentally (Fig. 4 and fig**. S9D–F** and **11**), it lacks a within-animal control, limiting its ability to account for individual variability. **(B)** Delivering unpredicted rewards during inter-trial interval while omitting the expected reward after the cue degrades the retrospective association between cue and reward. Because this association must be relearned, reacquisition proceeds at a speed similar to initial acquisition. **(C)** Simulation results confirm that retrospective association (predecessor representation contingency, PRC) between CS-A and reward decreases to zero during extinction, resulting in zero PRC at the first trial of reacquisition. **(D, E)** Because the underlying retrospective memory needs to be re-learned during reacquisition, the prospective association (successor representation contingency, SRC), hypothesized to drive conditioned behavior, develops at a similar speed to that during acquisition. **(F-J) (F)** Introducing a novel cue CS-B as a predictor of reward in a subsequent phase after learning CS-A-reward association is another means to degrade the CS-A←reward memory through retrospective extinction. **(G)** However, when reacquisition follows CS-B acquisition immediately, the estimated rate of CS-A is near zero. Under ANCCR, this zero rate compensates for degraded CS-A←reward memory in the Bayes’ rule computation, as both numerator and denominator are near zero. Consequently, despite the erased retrospective memory, the model predicts that an animal would infer a high prospective association (CS-A→reward) and acquire behavior rapidly. This caveat makes the paradigm unsuitable for testing memory degradation using reacquisition speed. **(H)** Simulation results confirm that the PRC between CS-A and reward decreases to zero during CS-B acquisition, while that for CS-B remains high. **(I, J)** Nonetheless, the SRC between CS-A and reward during reacquisition rapidly increases, recovering to a level comparable to that of CS-B, whose SRC remained intact throughout the experiment. **(K-P) (K, L)** Introducing standard extinction before reacquisition makes reacquisition speed depend on the degraded retrospective association, provided animals infer the same physical context across CS-A acquisition and CS-B acquisition. During standard extinction, ANCCR predicts that the estimated rate of CS-A recovers to its original level from CS-A acquisition. This recovery removes the zero-division compensation that previously enabled fast reacquisition despite degraded retrospective association, ensuring reacquisition speed reflects the strength of the remaining retrospective association. However, because ANCCR hypothesizes that associations are conditioned on physical context (see **Supplementary Note 5**), this memory erasure effect is only expected if animals treat the CS-A and CS-B acquisition phases as occurring in the same physical context. Since these phases contain entirely different sensory cues, animals may instead infer different physical contexts, thereby preventing retrospective memory erasure. Consistent with this prediction, experimental results in **fig. S9A-C** showed no evidence of memory erasure in this paradigm. **(M-O)** These simulations assume that animals infer the same physical context across all phases of the experiment, which need not be true. Under this assumption, the retrospective association between CS-A and reward is reduced during CS-B acquisition (M), resulting in slower recovery of the prospective association during reacquisition (N,O) compared with CS-B. However, because CS-B is acquired only after CS-A training, its acquisition is already facilitated by the previously learned high reward rate. Whether cross-cue generalization also contributes to the accelerated acquisition of CS-B beyond this effect remains unclear. As a result, CS-B acquisition is faster than true initial learning (i.e., CS-A acquisition) and cannot serve as an appropriate within-animal reference for assessing reacquisition of CS-B. Therefore, despite preserving its retrospective memory through standard extinction, CS-B cannot function as a valid within-animal control in this sequential design even when the same physical context was assumed across phases. Note that we are not questioning the general usefulness of such a design; we are instead just stating that comparing acquisition and reacquisition speeds in this paradigm will produce bias. **(P)** Similarly, maintaining CS-A omission trials while introducing a new CS-B reward pair during CS-B acquisition may reduce inference of a new physical context and promote erasure of CS-A reward memory. However, because CS-B acquisition remains facilitated by prior reward rate training, this paradigm suffers the same limitation, making it less suitable as a rigorous test of memory erasure. In addition to the effect from prior reward rate training, difference in novelty of cue during initial acquisition or recency from initial acquisition to reacquisition can affect the speed of acquisition and reacquisition, making it challenging to use CS-B as a within-animal control for CS-A. **(Q-S)** The two-cue paradigm used as the main test (Fig. 1**-3**) provides a rigorous within-animal test of memory erasure using reacquisition speed. Because CS-A and CS-B are trained simultaneously, it prevents differences in acquisition speed that arise in sequential designs, enabling acquisition of both cues as the reference of initial learning. In addition, all phases contain both CS-A and CS-B, minimizing the possibility that animals infer different physical contexts across phases. Further, CS-B standard extinction phase not only provides a within-animal control, but also ensures that reacquisition speed primarily reflects the strength of underlying memory, rather than other behavior control factors (F-J). In sum, although the two-cue paradigm may appear more complex than simpler retrospective extinction designs, it provides the most rigorous test of memory erasure while minimizing potential confounding factors.

**fig. S2.**
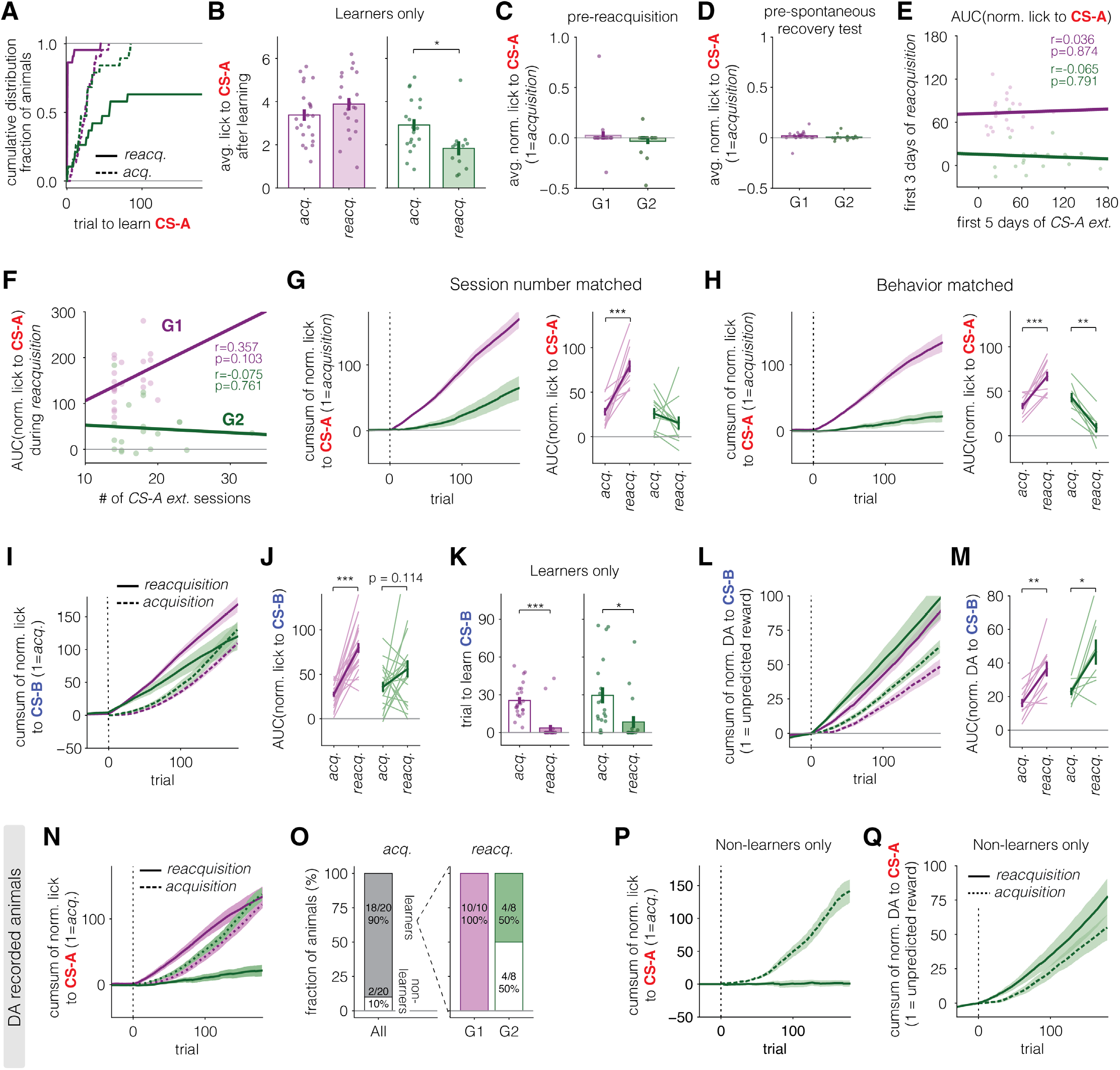
Differences in reacquisition or spontaneous recovery across groups are specific to CS-A, and not due to training history or pre-test behavior. **(A)** Cumulative distribution of animals based on trial to learn CS-A during acquisition and reacquisition. Mice were considered non-learners (trial to learn is infinity) if their post-learning average lick rate remained below 0.5 Hz and excluded from trial to learn analysis (7 of 19 G2 animals during reacquisition; see Methods). **(B)** Lick to CS-A after acquisition (empty bar) or reacquisition (filled). Animals with post-learning lick rates below 0.5 Hz were excluded from this analysis (7 of 19 G2 animals during reacquisition; same exclusion as in Fig. 1L). **(C)** No difference in pre-reacquisition licking behavior between groups. Individual animals are shown as circles. Error bar indicates SEM across animals. **(D)** No difference in pre-test (average lick on a day before the test) licking behavior between groups in the spontaneous recovery analysis. Note that pre-test lick was averaged across trials within a given session, whereas spontaneous recovery measure in Fig. 1N uses the AUC of lick vs. trial plot over the first 5 trials where spontaneous recovery is primarily observed. **(E)** No correlation between reacquisition speed and extinction speed to CS-A. **(F)** No correlation between reacquisition speed and the number of CS-A extinction sessions. Each point represents an individual animal; the line indicates linear regression within each group **(G)** Group-averaged cumulative sum of normalized lick to CS-A during acquisition (left) and AUC of normalized lick to CS-A over the first 6 days of acquisition and reacquisition (right), using animals matched for the number of CS-A extinction sessions across groups (n = 12 for G1, 11 for G2). Left, lines indicate mean across animals and shadings represent SEM. Right, light-colored lines indicate individual animals and error bar represents SEM. **(H)** Same as G but using animals matched for extinction behavior (n = 10 for G1, 8 for G2). For behavior matching, CS-A extinction phase was extended 10 days after animal reached behavioral extinction threshold (see Methods). **(I-K)** Although cross-cue generalization may have suppressed CS-B licking in a subset of G2 animals, resulting in no significant difference in AUC between acquisition and reacquisition when all animals were included (J; note that difference in AUC between acquisition and reacquisition is significant in both groups when the analysis window is restricted to the first two days, instead of three days, of acquisition/reacquisition; G1, 1.1×10-8; G2, 0.033), G2 still exhibits a similar trend of rapid reacquisition for CS-B as observed in G1 (I). When the analysis was restricted to CS-B learners (n=22/22 for reacquisition in G1, n=17/19 in G2; two non-learners in G2 also did not learn CS-A supporting the effect of cross-cue generalization), trial to learn was significantly faster during acquisition than during reacquisition in both groups with no group difference in learned trial either during acquisition (p=0.5622) or reacquisition (p=0.3883) (K). This supports that memory erasure due to retrospective extinction in G2 is largely specific to CS-A. **(L,M)** Reacquisition of DA to CS-B is more rapid than acquisition in both groups, supporting cue-specific memory erasure due to retrospective extinction. **(N,O)** The group difference in CS-A reacquisition speed (using the same analyses as in **Fig. 1J,K**) remained when the analysis was restricted to the subset of animals with recorded dopamine responses recorded (animals shown in Fig. 1O**–Q**). **(P,Q)** G2 animals that failed to reacquire conditioned behavior to CS-A (P) nevertheless reacquired dopamine responses to CS-A at a similar speed to that observed during acquisition (Q).

**fig. S3.**
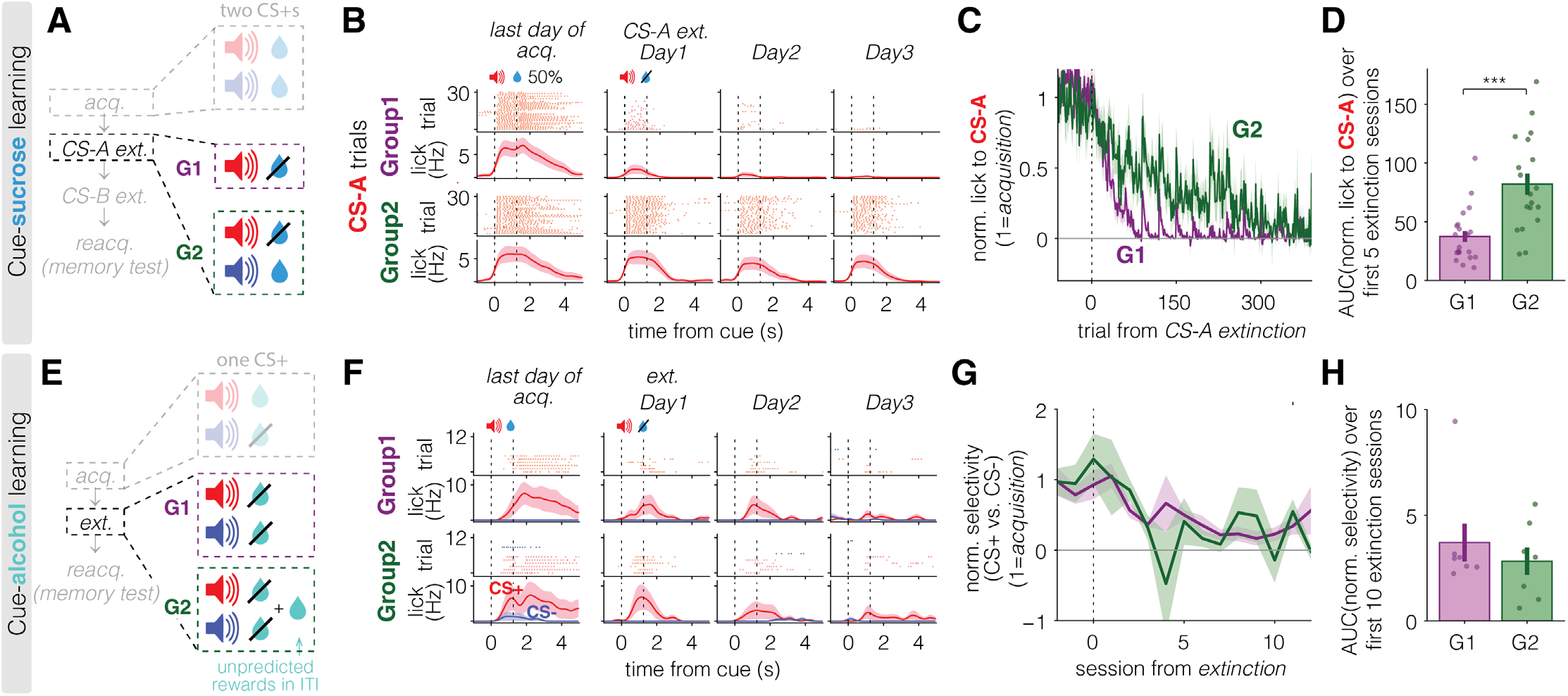
Slow CS-A extinction in Group2 may result from generalization across cues. **(A-D)** Behavioral results from cue-sucrose learning. **(A)** Task schematic for CS-A extinction in cue-sucrose learning (main task from **Fig1**). During CS-A extinction, G2 continued to experience CS-B reward trials, whereas G1 did not. Existence of CS-B rewarded trials in G2 may slow the extinction of CS-A due to generalization across cues acting as a protective mechanism against degradation of CS-A -reward association. **(B)** Example lick raster plots (top row) and PSTHs (bottom row) during CS-A extinction for one representative mouse per group in cue-sucrose learning. The first column shows data from the last day of acquisition, and remaining columns (Day 1 to 3) show CS-A extinction. **(C)** Group-averaged normalized lick to CS-A across trials during CS-A extinction. Trial 0 marks the first CS-A trial in CS-A extinction. Lines represent mean across animals and shadings represent SEM. **(D)** AUC of normalized lick to CS-A over the first 5 days of CS-A extinction is significantly larger in G2 than G1. Individual animals are shown as circles. Error bar indicates SEM across animals. **(E-H)** Behavioral results from cue-alcohol learning. **(E)** Task schematic for extinction in cue-alcohol learning. Here, only one cue (CS+) was paired with reward during acquisition. In G2, unpredicted rewards were delivered in the inter-trial interval, rather than rewards paired with another CS+. This task paradigm avoids any slowdown in the extinction of CS+ due to generalization between cues. **(F)** Same as B, but from mice in cue-alcohol learning. CS+ and CS-trials are shown in red and blue, respectively. **(G)** Group-averaged normalized selectivity (see Methods) for CS+ over CS-across extinction sessions. Each animal’s selectivity was normalized to its average during the last 5 days of acquisition. Session 0 indicates the first session of extinction. **(H)** AUC of normalized selectivity for CS+ over CS-during the first 10 days of extinction.

**fig. S4.**
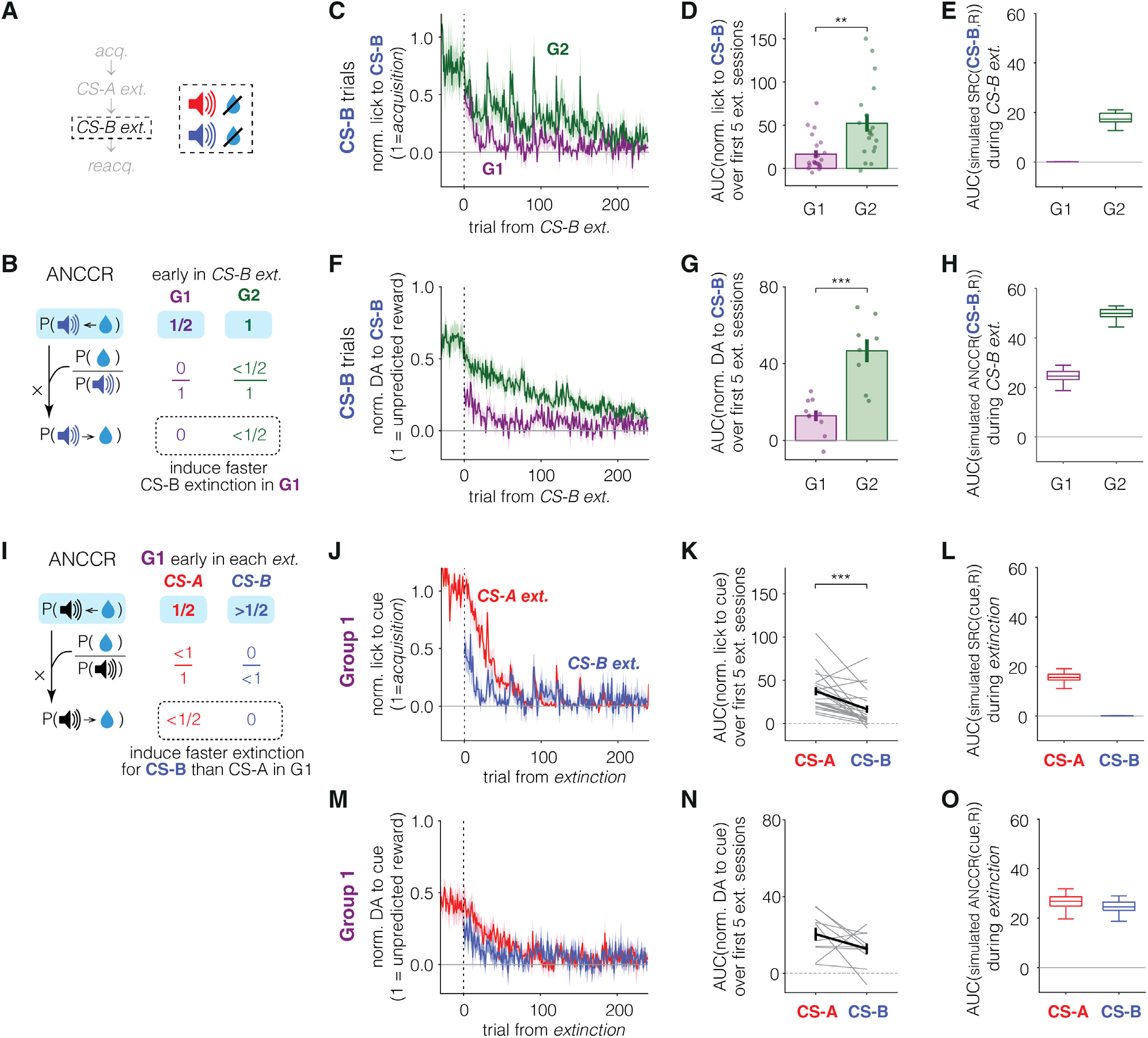
ANCCR predicts faster CS-B extinction in Group1 than in Group2 (A-H) CS-B extinction is faster in G1 than in G2. **(A)** Task schematic for CS-B extinction. **(B)** Prediction from ANCCR. Because P(reward) had already decreased to zero during CS-A extinction in G1, CS-B extinction is predicted to occur faster in G1 than in G2. **(C)** Group-averaged normalized lick to CS-B across trials during CS-B extinction. Trial 0 marks the first CS-B trial in CS-B extinction. Lines indicate mean and shadings represent SEM across animals. **(D)** AUC of normalized lick to CS-B over the first 5 days of CS-B extinction. **(E)** AUC of simulated prospective association (successor representation of contingency, SRC) between CS-B and reward, used to model behavior. **(F)** Group-averaged normalized DA to CS-B across trials during CS-B extinction. **(G)** AUC of normalized DA to CS-B over the first 5 days of CS-B extinction. **(H)** AUC of simulated ANCCR between CS-B and reward during CS-B extinction. ANCCR reflects a weighted sum of prospective and retrospective association, hypothesized to represent dopamine response. Note that this weight may itself be dynamic depending on environmental statistics, but we have chosen not to include a fit of this dynamic weight in the ANCCR simulations. **(I-O)** CS-B extinction is faster than CS-A extinction in G1. **(I)** Prediction from ANCCR. In early CS-A extinction, P(rewards) decreases from its positive acquisition value to zero, driving extinction. In contrast, because P(reward) had already reached zero during CS-A extinction, subsequent CS-B extinction is predicted to occur more rapidly. **(J)** Group-averaged normalized lick to CS-A and CS-B across extinction trials in G1. **(K)** AUC of normalized lick to CS-A and CS-B over the first 5 days of each extinction phase in G1. Gray line indicates individual animals; error bar represents mean across animals. **(L)** AUC of simulated SRC between cue (CS-A or CS-B) and reward during each extinction phase in G1. **(M)** Group-averaged normalized DA to CS-A and CS-B across extinction trials in G1. **(N)** AUC of normalized DA to CS-A and CS-B over the first 5 days of each extinction phase in G1. **(O)** AUC of simulated ANCCR between cue (CS-A or CS-B) and reward during each extinction phase in G1. Note that there is no difference between CS-A and CS-B because higher retrospective association for CS-B (due to its lower baseline rate learned during preceding CS-A extinction) compensates for its weaker prospective association.

**fig. S5.**
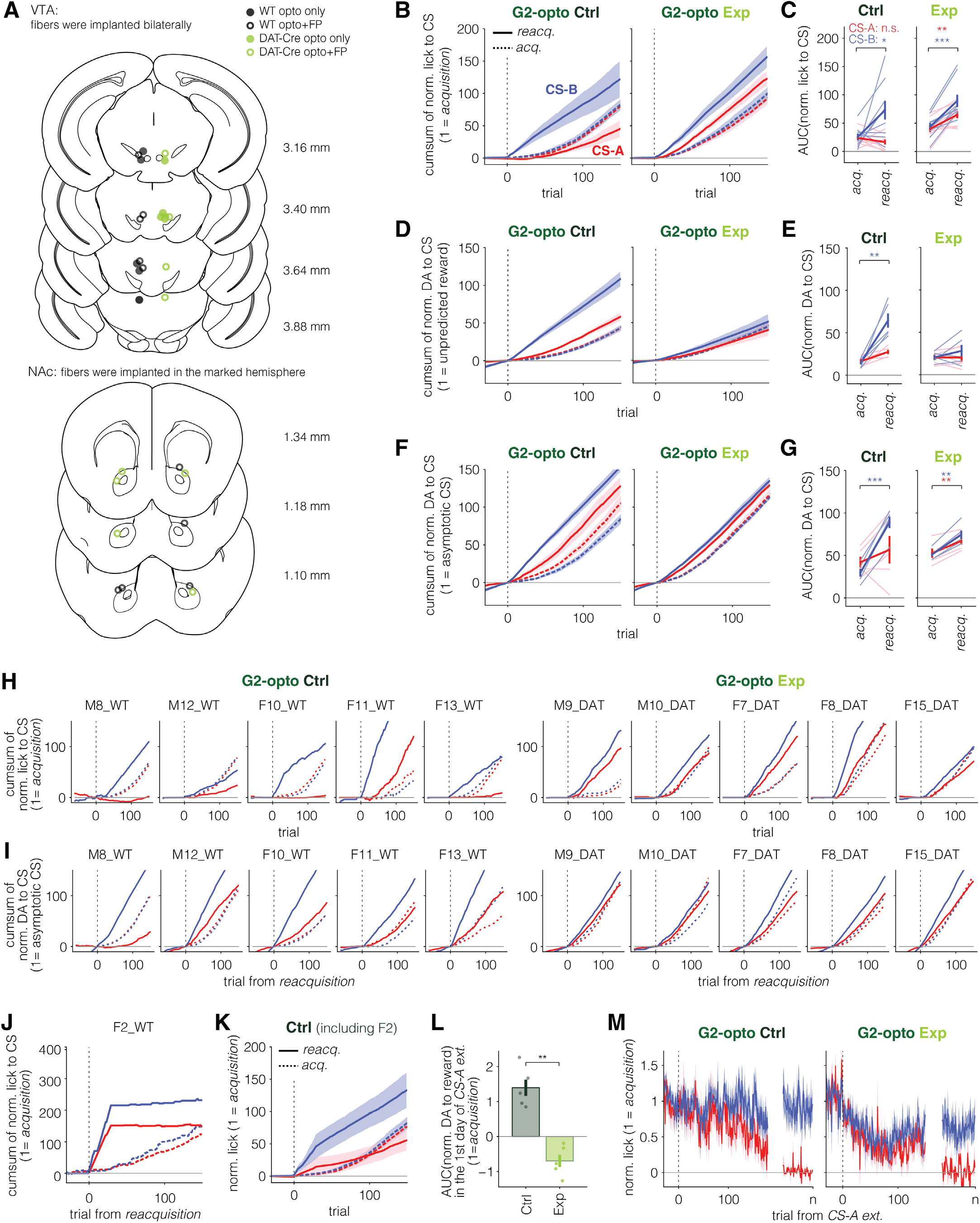
Reacquisition of dopamine response to cue in the VTA optogenetic experiment. **(A)** Histological location of optic fibers for photometry recording of dopamine release in nucleus accumbens (top) and for optogenetic manipulation of VTA dopamine neurons (bottom). Dopamine release was simultaneously measured in a subset of animals for VTA optogenetic experiment (5 of 9 for G1, 5 of 9 for G2). The fiber was mislocated in one WT and one DAT-Cre animal (whose fibers are located at AP 3.88; fiber is located dorsal to VTA). but we included these animals: WT animal was in the control group with no opsin expression, and simultaneous fiber photometry recording in NAc in DAT-Cre animal confirmed strong inhibition of dopamine release, indicating sufficient light reached the VTA despite the dorsal placement of the fiber. The results did not change depending on the inclusion of these animals. **(B-C)** Group-averaged normalized lick to CS (B) and AUC of normalized lick to CS (C) for all animals including those with and without dopamine measurements. These panels are identical to Fig. 2D**-G**, except that behavioral responses to both CS-A and CS-B are shown. **(D-G)** Group-averaged normalized DA to CS (D, F) and AUC of normalized DA to CS (E, G) for animals in which dopamine release was measured. For D and E, DA to CS was normalized to maximum DA to reward across first three acquisition sessions. To dissociate the speed of learning (inflection point from zero) and the magnitude of response (asymptotic level of DA to CS), in F and G, DA to CS was normalized to asymptotic DA to CS in last 3 days of each phase. In both cases, reacquisition was rapid selectively to CS-B in WT animals. In DAT-Cre, the comparison between acquisition and reacquisition was dependent on normalization methods. Because dopamine inhibition during extinction can reduce estimated reward magnitude, overall cue responses (both CS-A and CS-B) was generally lower during reacquisition when compared to acquisition (see H and I). However, regardless of normalization method, no group difference in reacquisition speed was observed. **(H-I)** Cumulative sum of normalized lick (H) and DA (I) to CS for individual animals using both normalization methods. **(J-K)** One control animal (F2-WT) was excluded from analysis because its behavior on the first day of reacquisition was markedly different from all subsequent days (J), making its learning uninterpretable. Including this animal did not change the results (K; same analysis as in Fig. 2E but with F2-WT included). **(L)** Normalized dopamine response to reward on the first day of CS-A extinction. Dopamine response to reward was significantly suppressed in Exp group, confirming effective optogenetic inhibition in Exp. Individual animals are shown as circles. Error bar indicates SEM across animals. **(M)** Group-averaged normalized lick to CSs during CS-A extinction. Trial n indicates the last trial of CS-A extinction phase. Omission of rewards following CS-A induced a comparable reduction in anticipatory behavior to CS-A across groups. However, Exp group also showed a reduction in anticipatory behavior to CS-B, consistent with prior reports (*132*, *133*). This generalized reduction in responding to reward-associated cues in Exp is consistent with the ANCCR prediction that below-baseline dopamine response at reward induces a reduction in the estimated reward magnitude.

**fig. S6.**
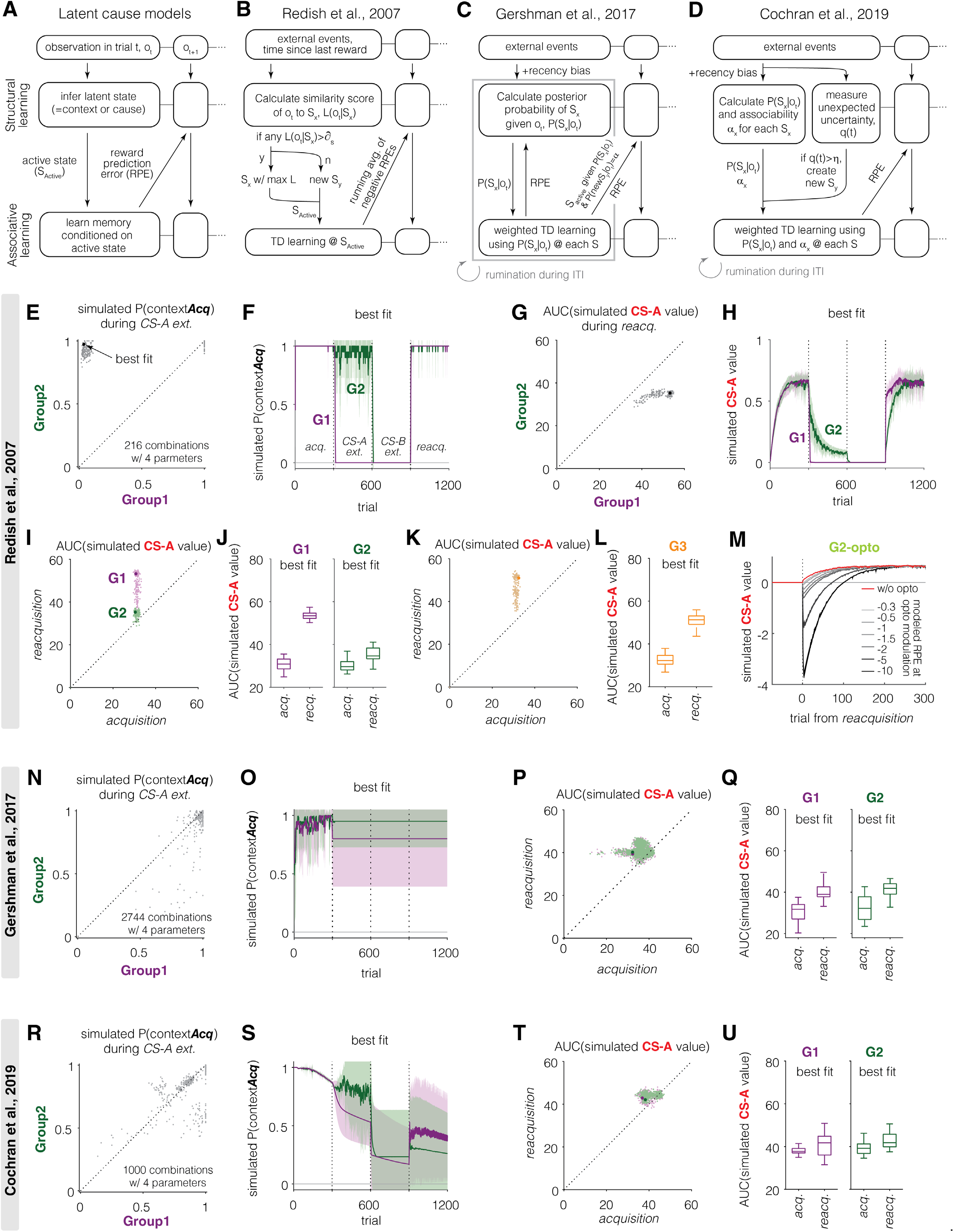
Variants of latent cause model do not capture behavioral results (A-D) Schematic of latent cause models. **(A)** General framework. An agent infers a latent state (cause or context) from observations (o_t_; structural learning), then learns cue-outcome memory conditioned on the active state (S_active_; associative learning). Prediction error from associative learning influences latent state inference. **(B)** Redish et al. (2007): Observations are compared to stored prototypes of existing latent states S_x_. If similarity of any states exceeds a threshold *∂_s_*, the best-matching state is selected as active; otherwise, a new state S_y_ is created and selected as active. Sustained negative reward prediction errors (RPEs) across timesteps increase the likelihood of splitting into new states. **(C)** Gershman et al. (2017): The agent infers a posterior distribution over latent states and updates memory for all states in proportion to this posterior. Structural learning (posterior update) and associative learning (memory update) alternate, with additional iterations (‘rumination’) occurring during inter-trial interval (ITI). Although associative updates are distributed across states, each trial is still assumed to be generated by a single latent cause, S_active_. The state with the highest posterior at the end of each trial is selected and used as prior input for the next trial. Rumination due to replays does not change inferred state, S_active_, but it updates the memory associated with that state. This assumption is critical for explaining extinction-retrieval results (**fig. S14**). **(D)** Cochran et al. (2019): the agent infers posterior probability and estimates associability *α_x_* for each state given the observation. If unexpected uncertainty q(t) exceeds a threshold η, a new state is created. Memory is updated for all states, scaled by both the posterior and associability. These updates may iterate during ITI via rumination under the same posterior beliefs. **(E-M)** Redish model captures results from G1 and G2, but not G3 or G2 with optogenetic manipulation. **(E)** Simulated probability of inferring acquisition context, P(contextAcq), during CS-A extinction for each group across all parameter combinations (gray). Black dot indicates the best fit parameter combination (see Methods). **(F)** Simulated P(contextAcq) over trials, across phases, using the best fit. Lines indicate mean across iterations and shadings represent STD. **(G)** AUC of simulated CS-A value during reacquisition for each group across all parameter combinations. **(H)** Simulated CS-A value over trials using the best fit. **(I)** AUC of simulated CS-A value during acquisition vs. reacquisition for each group. Light dots indicate individual parameter combinations and darker dots mark the best fit combination. **(J)** AUC of simulated CS-A value during acquisition and reacquisition for each group, using the best fit. In box plot, the central line indicates the median of iterations; box edges mark the 25^th^ and 75^th^ percentiles; whiskers extend to the most extreme data points within 1.5× the interquartile range. **(K-L)** Same as I-J, but for Group3 (G3), which behaves like G1 and not G2. **(M)** Simulated CS-A value over trials during reacquisition in G2-opto. Reacquisition was slower with optogenetic manipulation (greys) compared to without manipulation (red). Increasing the magnitude of modeled negative RPE during optogenetic manipulation (darker grey) progressively slowed reacquisition.

**fig. S7.**
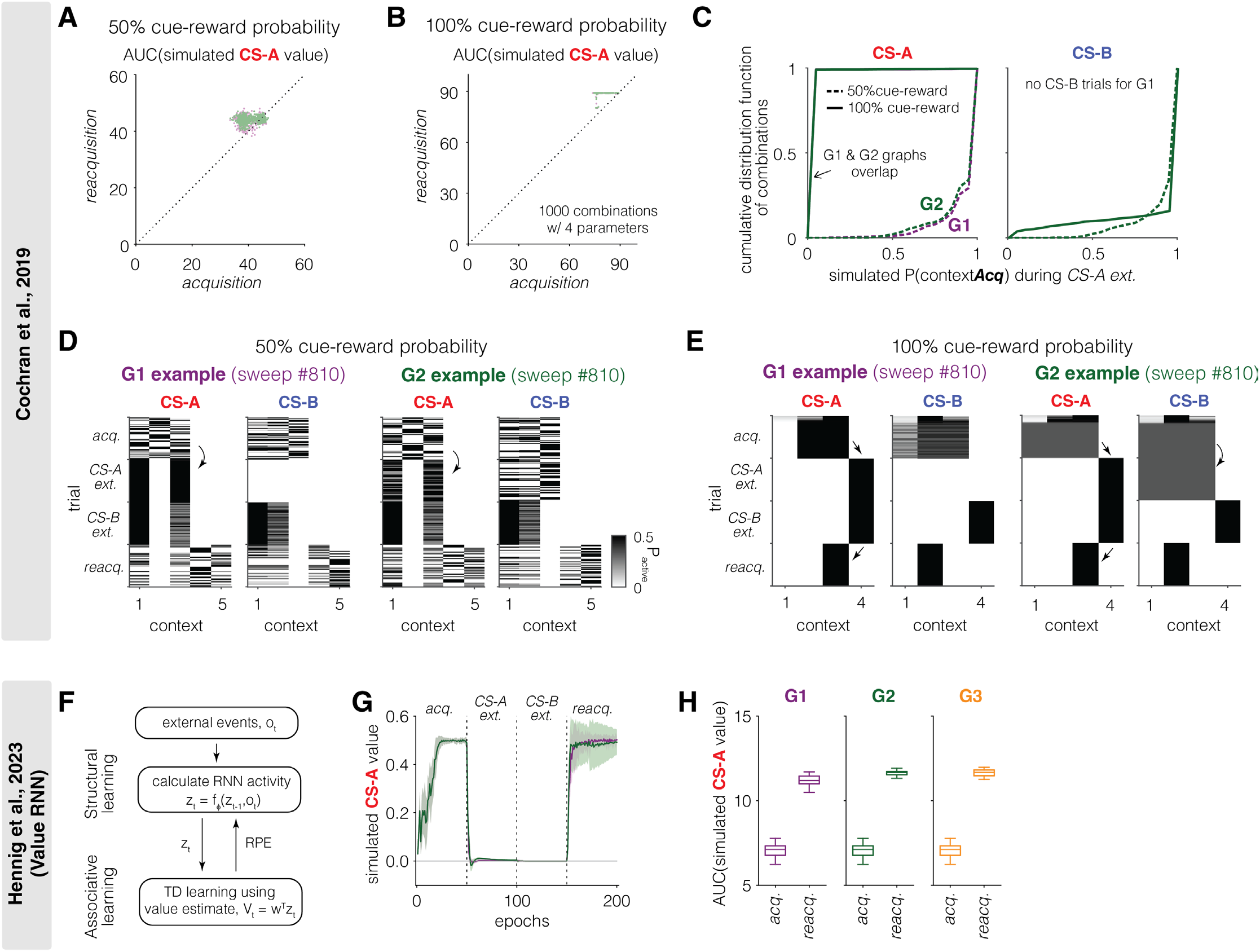
Additional simulation results from variants of latent cause model (A-E) Simulation results from Cochran et al. model. **(A-B)** Simulated AUC of CS-A value during acquisition vs. reacquisition for all parameter combinations under 50% (A) or 100% (B) cue-reward probability. In both conditions, the model does not predict different speed of reacquisition between groups. Scatters indicate individual parameter combinations (Green, G1; Purple, G2). **(C)** Cumulative distribution of simulated P(contextAcq) during CS-A extinction for CS-A (left) and CS-B (right) trials. Note that there are no CS-B trials for G1 during CS-A extinction. At 50% cue-reward probability (dotted), the model continues to infer contextAcq in both groups. At 100% cue-reward probability (solid), the model infers a new context for CS-A trials (left) in both groups but maintains contextAcq for CS-B trials in G2 (right). This shows that the presence of CS-B trials during CS-A extinction in G2 does not promote inference of contextAcq, even when cue-reward contingencies are deterministic. Instead, the model segregates CS-A and CS-B trials into distinct contexts. **(D-E)** Trial-by-trial simulated context inference across phases for an example parameter combination (#810) under 50% (D) or 100% (E) cue-reward probability. Each row is a trial. Color scale shows inferred probability for each context. **(F-H)** Simulation results from value RNN (Hennig et al., 2023). **(F)** Schematic of value RNN. Activity of RNN units represents inferred state (‘belief state’) given trial. The weights for neural network (structural learning) and value calculation (associative learning) are trained to minimize RPE. **(G)** Simulated CS-A value across trials. Value RNN predicts rapid reacquisition in both G1 and G2 and does not capture behavior results. **(H)** AUC of simulated CS-A value during acquisition and reacquisition for each group. Value RNN fails to distinguish groups. In box plot, the central line indicates the median of iterations; box edges mark the 25^th^ and 75^th^ percentiles; whiskers extend to the most extreme data points within 1.5× the interquartile range.

**fig. S8.**
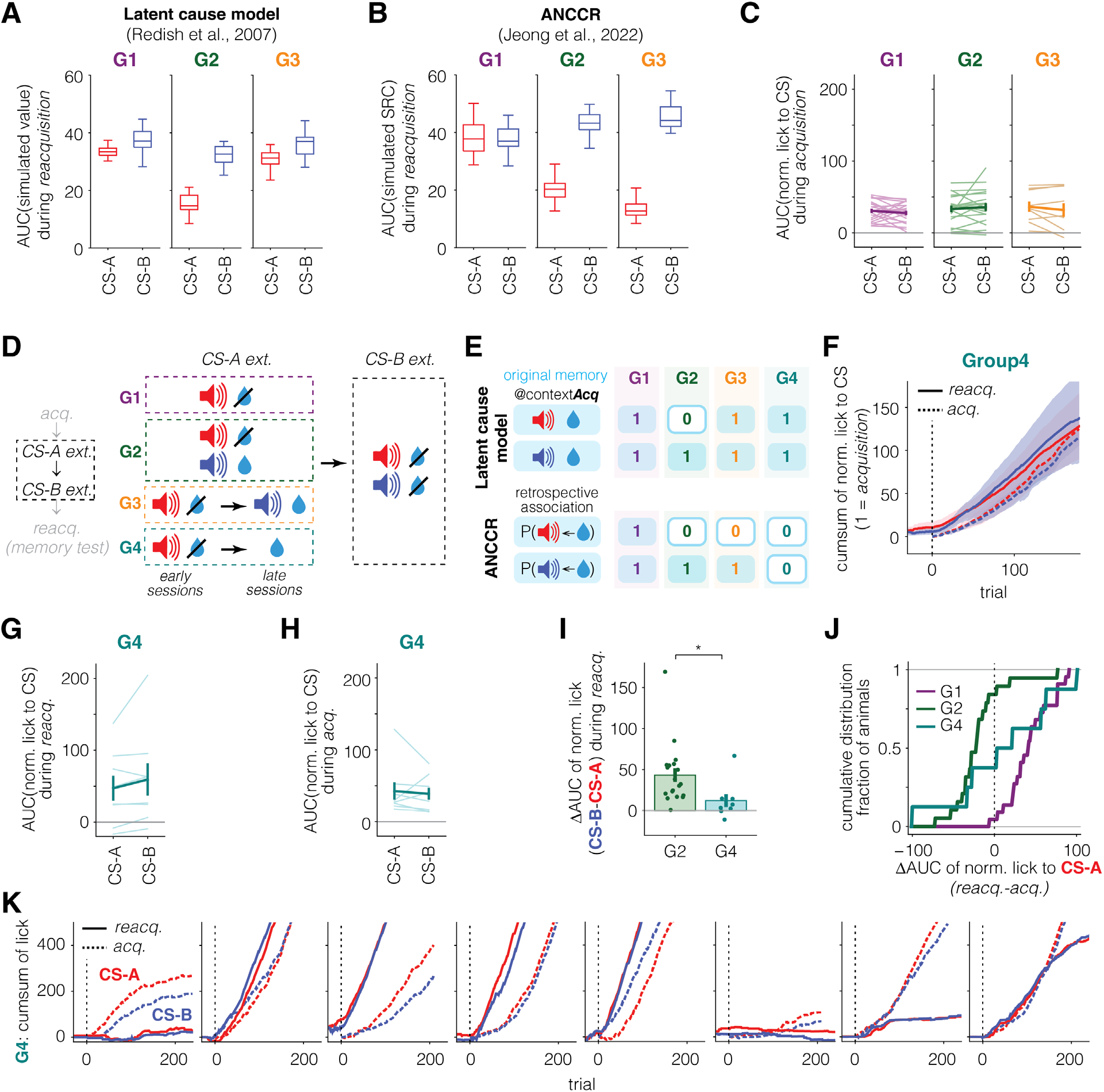
Presentation of uncued reward erases associated memories to the reward. **(A-B)** Simulations. AUC of simulated CS-A and CSB value (latent cause model, A) or SRC (ANCCR, B) during reacquisition, reflecting speed of reacquisition. **(C)** AUC of normalized lick to CS-A and CS-B during acquisition. **(D)** Task schematic for CS-A extinction. Group4 (G4) underwent CS-A extinction like G3, but uncued rewards were delivered instead of CS-B reward trials in late extinction sessions. **(E)** Model predictions. Latent cause model (top) predicts that original cue-reward memories are preserved in G4, as CS-A omission trials in early sessions drive context switch. In contrast, ANCCR (bottom) predicts unlearning of both CS-A and CS-B, as uncued rewards in late sessions degrade all retrospective associations paired with the reward. **(F-K)** Behavioral results from G4 support ANCCR prediction. **(F)** Group-averaged cumulative sum of normalized lick to CS-A (red) and CS-B (blue) during acquisition (dotted) and reacquisition (solid). On average, reacquisition was as slow as acquisition for both cues. Lines indicate mean across animals and shadings represent SEM. **(G-H)** AUC of normalized lick to each cue during reacquisition (G) and acquisition (H). Individual animals are shown as light-colored lines. Error bar indicates SEM across animals. **(I)** G4 shows significantly smaller difference in reacquisition speed between cues during reacquisition than G2. Individual animals are shown as circles. Error bar indicates SEM across animals. **(J)** Cumulative distribution of animals based on reacquisition vs. acquisition difference in learning speed. G1 and G2 show opposing biases (faster or slower reacquisition), while G4 shows a wide range of distribution resulting in no bias on average. This variability may depend on how consistently animals associate rewards during unpaired reward-only sessions with the same context as the rest of experiment. Because a physical context is nothing but a set of sensory cues, some animals may infer the unpaired reward-only sessions lacking any neutral cues present elsewhere in the task as a distinct context, potentially preventing degradation of retrospective associations stored in the original context and resulting in little or no impairment in reacquisition. In contrast, G3 (Fig. 3) had auditory cues during these sessions and showed strong impairment that was consistent across animals. These results suggest that sensory cues support inference of a physical context, and that their absence can lead to misassignment of context (see **Extended Data note 5** and the Discussion section in main text for more discussion about context dependency in ANCCR vs. latent cause models). **(K)** Cumulative sum of lick to CS-A (red) and CS-B (blue) during acquisition and reacquisition for individual animals.

**fig. S9.**
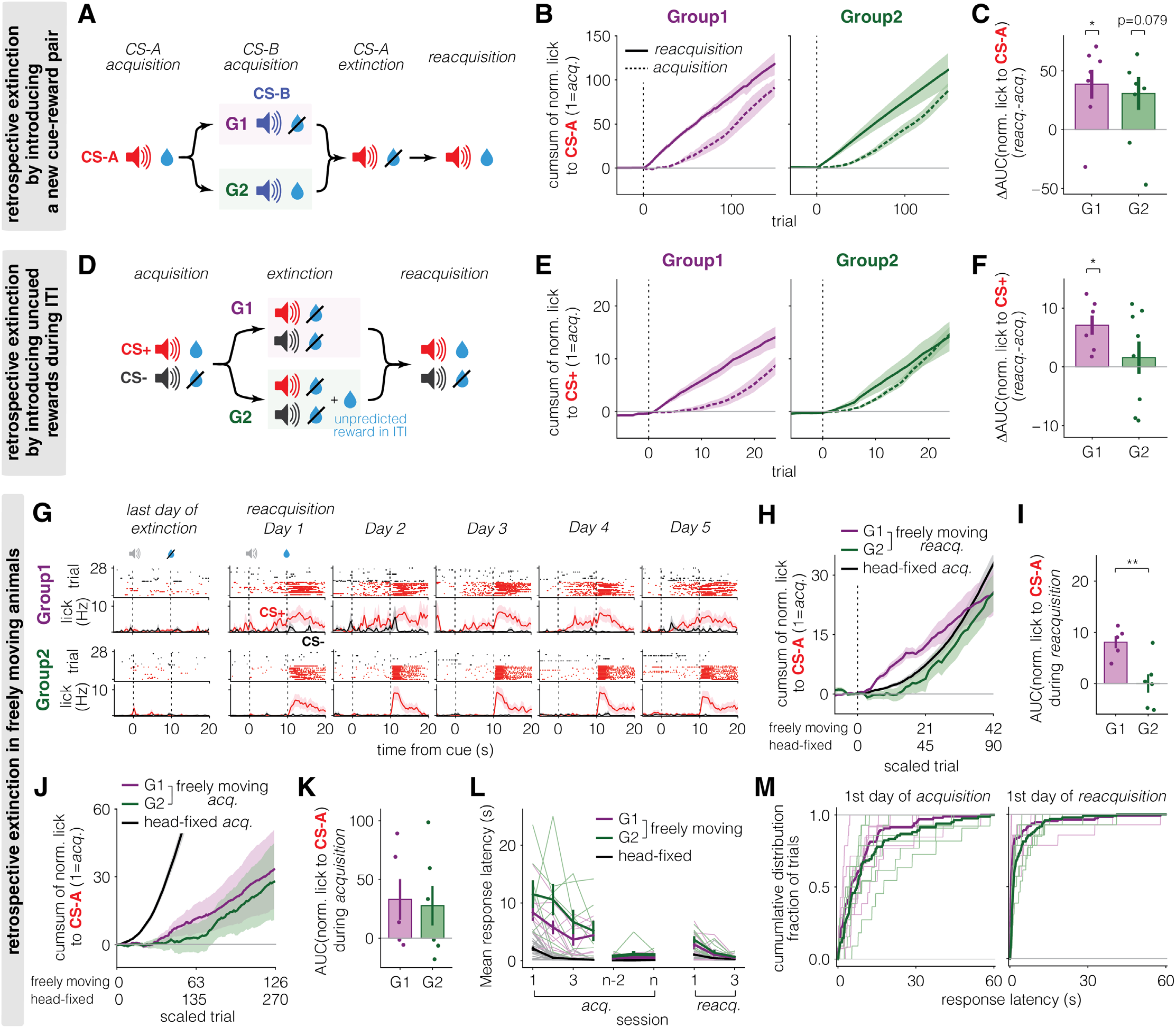
Boundary conditions for retrospective memory erasure. **(A-C)** Introducing a new cue-reward pair after learning the original cue-reward association does not induce retrospective memory erasure, consistent with the two experiences being inferred as separate physical contexts. **(A)** Experimental design. Following CS-A acquisition, mice experienced either presentation of CS-B alone (G1) or CS-B and reward pairs. This experiment tested whether learning a new cue-reward pair after learning the original cue-reward association erases the original memory. **(B, C)** Group-averaged cumulative sum of normalized lick to CS-A during acquisition and reacquisition (B) and difference in AUC of normalized lick to CS-A between phases (C). Both groups showed rapid reacquisition, indicating that rewarded CS-B trials did not erase the original CS-A memory when CS-A and CS-B were never experienced within the same session (i.e., same physical context). Because a physical context is defined by sensory information in the environment, the complete absence of CS-A and exclusive presentation of CS-B during CS-B acquisition phase can cause animals to infer that they have entered a different physical context from the one in which the original CS-A←reward association is learned. Consequently, rewards in CS-B trials cannot induce erasure of retrospective CS-A←reward association. This dependence on physical context in ANCCR is different from inference of an unobservable latent context in latent cause models. See Extended Data fig 8, Extended Data note 5, and the Discussion section for further discussion. Continuously presenting CS-A without reward while introducing a novel CS-B reward pair could potentially reduce these physical context-related issues. However, this paradigm is functionally very similar to the original G2 design, differing only in the inclusion of CS-B trials during acquisition. The original G2 paradigm therefore provides a cleaner test because nothing about CS-B reward contingency changes from *acquisition* to *CS-A extinction* phase, while CS-A memory is selectively degraded during *CS-A extinction* phase. **(D-F)** Introducing uncued rewards during inter-trial interval (ITI) induces retrospective memory erasure while avoiding physical context ambiguity. **(D)** Experimental design. The main experiments (Fig. 1-2) used two rewarded cues so that memory erasure could be manipulated in one cue while the second cue served as a within-animal control. A simpler implementation of retrospective extinction paradigm is to deliver uncued rewards during ITI following acquisition of a single cue-reward association. Accordingly, after acquisition, G1 underwent standard extinction in which the expected reward was omitted, whereas G2 underwent retrospective extinction in which the expected reward was omitted and uncued rewards were delivered randomly during ITI. Because uncued rewards were delivered within the same sessions in which the extinguished cue was presented, this design avoids the physical context ambiguity illustrated in A-C while preserving the critical feature of retrospective extinction. The same design was used for cue-alcohol (Fig. 4) and cue-shock (**fig. S10**) experiments. **(E, F)** Same analyses as in B, C. G1 reacquired behavior significantly faster than during initial acquisition, whereas G2 reacquired at a rate comparable to initial acquisition, consistent with memory erasure via retrospective extinction. The larger individual variability of G2 during reacquisition in this paradigm than in the main experiment (Fig. 1) is consistent with simulations (**fig. S1D**, *reacq.* vs. Fig. 1R, CS-A). **(G-M)** In freely moving animals, conditioned behavior develops through both learning of cue-reward association and learning of non-associative task structure, making acquisition speed of conditioned behavior a poor proxy for speed of associative learning strength. **(G)** Representative lick rasters (top row) and PSTHs (bottom row) from one mouse in each group (top, G1; bottom, G2). The first column shows the final extinction day (Day -1), and the remaining columns show the first five reacquisition days. G1 reacquired anticipatory licking to CS+ (red) immediately on Day 1, whereas G2 showed delayed reacquisition, with anticipatory licking emerging around Day 3. PSTHs show mean ± SEM across trials. **(H,I)** Group-averaged cumulative sum (E) and AUC (F) of normalized anticipatory licking to CS+ during reacquisition in freely moving animals (purple, G1; green, G2). Freely moving data are overlaid with acquisition data from head-fixed animals (black) shown in the main figures. Trial numbers were scaled so that equivalent training time was aligned across freely moving and head-fixed conditions. Reacquisition speed in freely moving G2 was similar to the speed of initial acquisition in head-fixed animals, consistent with memory erasure through retrospective memory. **(J,K)** Same analyses as in H,I, but during initial acquisition of freely moving animals. AUC was calculated across the first nine acquisition sessions. No difference in acquisition rate was observed between freely moving G1 and G2. However, acquisition of conditioned licking in both freely moving groups was substantially slower than either head-fixed acquisition (black) or freely moving reacquisition (E,F), which is likely due to the non-associative learning factor discussed below. **(L,M)** Mean response latency across acquisition and reacquisition sessions (I), and cumulative distribution of response latency on the first day of acquisition and reacquisition (J). During early acquisition, freely moving animals exhibited much longer and more variable response latencies (∼10s) than during reacquisition or in head-fixed animals. Unlike in the head-fixed condition, where animals can immediately detect and consume reward because the reward port is positioned directly in front of animal’s mouth, freely moving animals must first learn that reward is occasionally delivered at the reward port and subsequently adopt strategies that allow rapid detection and collection of reward (e.g., repeatedly checking the port or remaining nearby the port). Moreover, because reward collection in freely moving animals requires substantially greater effort than the simple licking response required in the head-fixed condition, the rate at which animals acquire efficient reward collection behavior varies considerably across individuals, likely reflecting difference in motivational level. Consequently, the development of conditioned behavior during initial acquisition reflects both learning of the cue-reward association and learning of efficient reward collection. Because reward collection behavior has already been acquired by the time reacquisition begins (as evidenced by substantially faster response latency during reacquisition compared to the early days of acquisition), comparing reacquisition speed to acquisition speed within the freely moving paradigm does not directly test memory erasure. Instead, we reason that the more appropriate comparison is the absolute time required for associative learning under conditions where reward collection does not constrain performance (i.e., the head-fixed preparation). Consistent with this interpretation, reacquisition in freely moving G2 required a similar amount of training time as initial acquisition in head-fixed animals. We previously showed that absolute amount of time for learning is maintained across different conditions by scaling learning rate with inter-reward interval (*45*). Assuming this principle also holds in freely moving animals, these results suggest that reacquisition in G2 proceeds at the rate expected for initial learning of cue-reward association, consistent with retrospective memory erasure.

**fig. S10.**
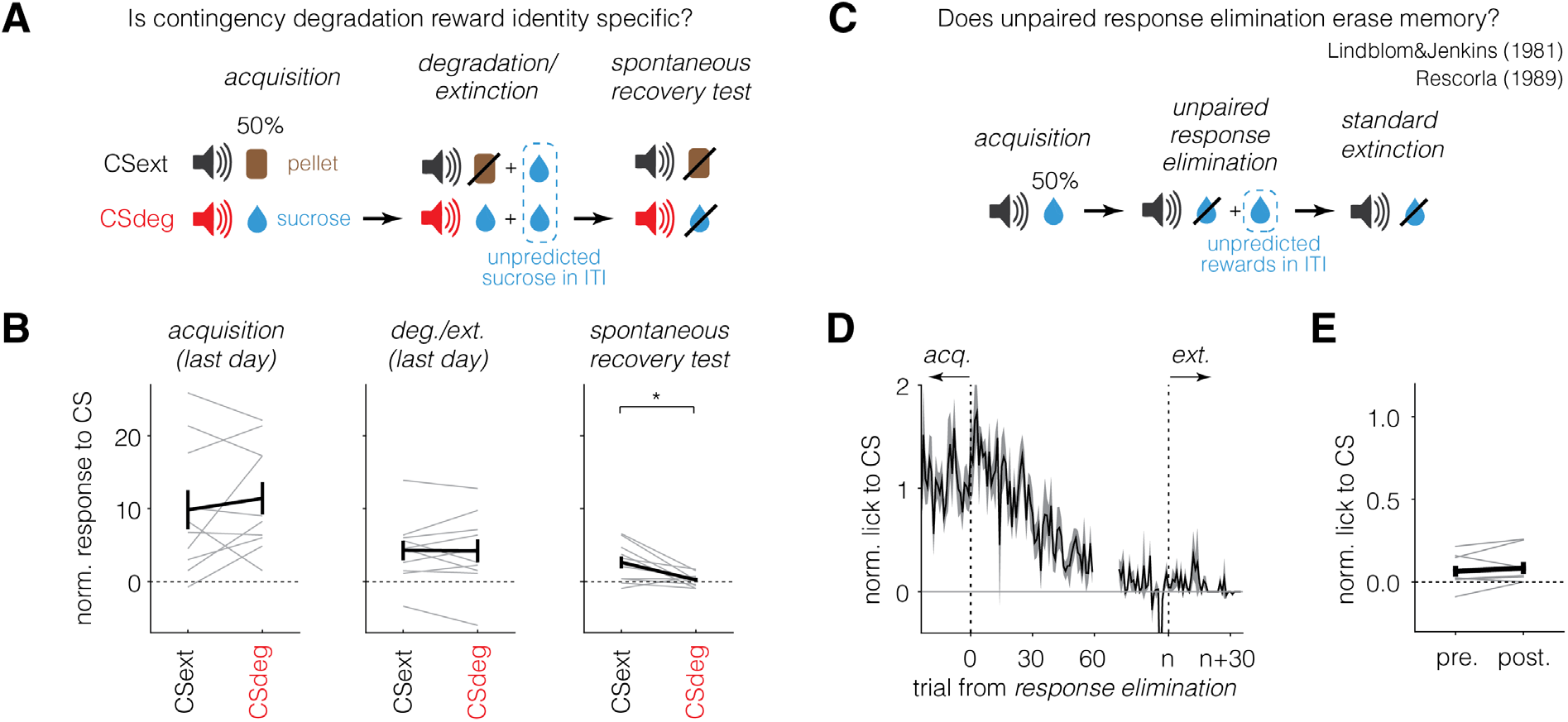
Additional data supporting retrospective learning. **(A-B)** Contingency degradation is reward-identity specific in freely moving rats. **(A)** Task schematic. After the acquisition of two cue-reward pairs (CSdeg-sucrose and CSext-pellet), one cue (CSdeg) underwent contingency degradation while the other cue (CSext) underwent standard extinction. Memory was tested in a spontaneous recovery test. **(B)** Normalized response to cues in each phase. More spontaneous recovery was observed to CSext than CSdeg (right), while no difference was found during acquisition (left) or degradation/extinction (middle). **(C-E)** Unpredicted rewards eliminate memory in head-fixed mice. **(C)** Task schematic. After cue-reward learning, response elimination was induced by omitting rewards after cue and delivering uncued rewards during ITI. Standard extinction followed to assess memory retention. **(D)** Average normalized lick to CS over trials. Lick to CS declined during response elimination and did not recover in extinction as opposed to previous findings from pigeon auto shaping (*52*, *53*). **(E)** Normalized lick to CS during pre- and post-transition from response elimination to standard extinction.

**fig. S11.**
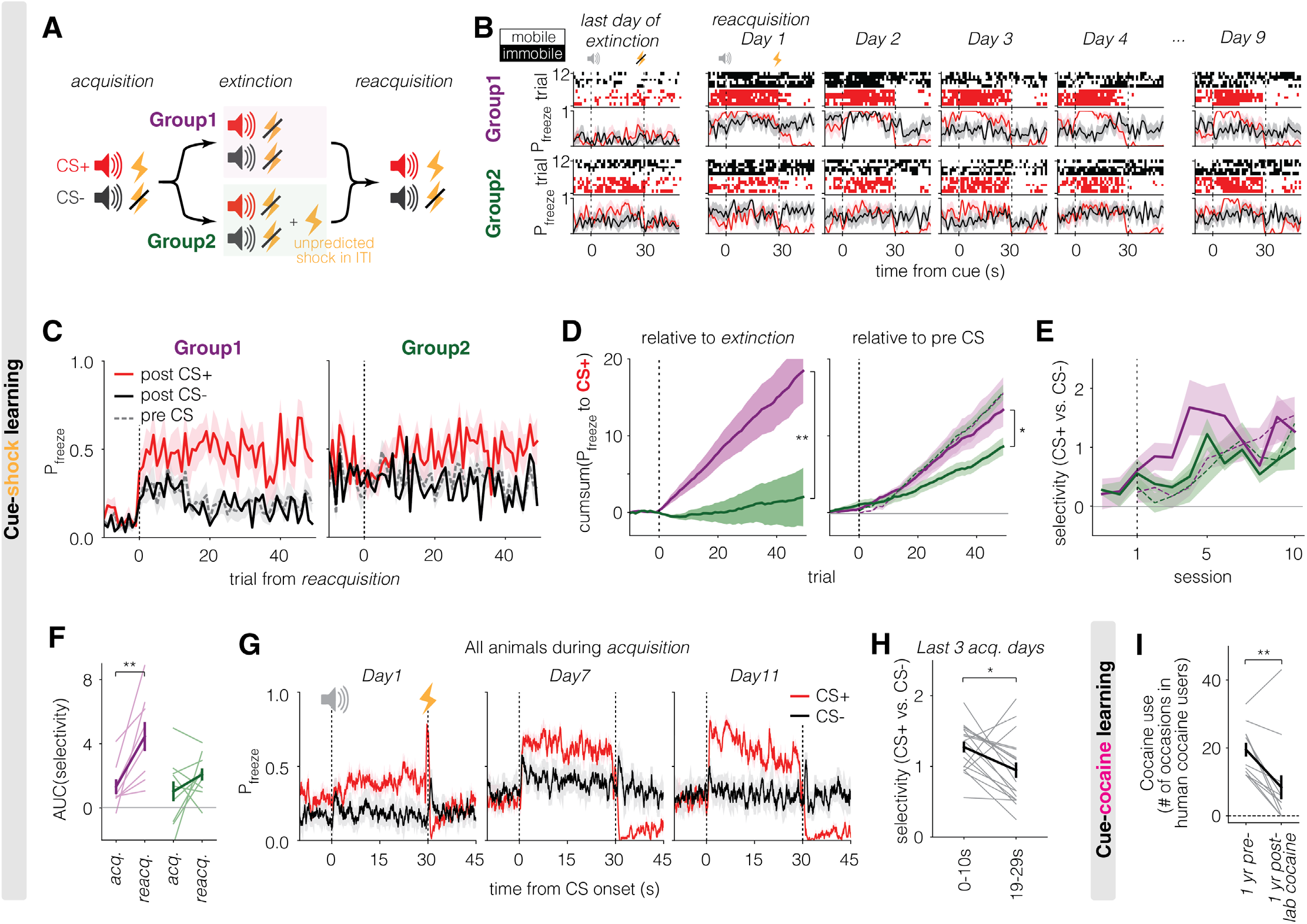
Retrospective extinction degrades memory in preliminary studies of cue-shock and cue-cocaine learning. **(A-F)** Cue-shock learning. Same format as Fig. 4A**-F**, using immobile probability (P_freeze_) during a 10 s window as behavior measure. **(A)** Schematic of cue-shock learning. Similar task structure as cue-alcohol learning, but with shocks instead of alcohol. On each trial, cue lasted for 30 s to allow a sufficient window for immobility analysis. **(B)** Example heatmap (top row) and PSTHs (bottom row) of immobility for representative mice. In heatmap, red and black indicate immobile states, while white indicates mobile states. In G1, selective immobility to CS+ (red) emerged by Day 3, with reduced immobility during CS-(black) and baseline. In contrast, G2 showed sustained immobility to both cue and baseline, with delayed selectivity emerging only by Day 10. **(C)** G1 rapidly develops immobility to CS+ during reacquisition, while G2 shows nonselective immobility to both cues and baseline during extinction and a delayed emergence of CS+ selectivity during reacquisition. **(D)** Cumulative sum of immobile probability to CS+ across trials during acquisition (dotted) and reacquisition (solid). When P_freeze_ to CS+ was calculated as the difference of P_freeze_ between pre and post CS+ delivery, reacquisition of immobility to CS+ alone was not faster than acquisition, even in G1 (right). However, selectivity to CS+ over CS-developed more rapidly during reacquisition in G1 (see **fig. S11E**), suggesting that learning involves both response to CS+ and suppression of response to CS-. Consistent with this, reacquisition appeared substantially faster in G1 than G2 when P_freeze_ to CS+ was instead calculated relative to the average post-CS+ immobility during the two days of extinction (left). **(E)** Group averaged selectivity (CS+ vs. CS-) based on anticipatory immobility during acquisition (dotted) and reacquisition (solid). G1, but not G2, shows rapid reacquisition. **(F)** AUC of selectivity over first 5 days of acquisition or reacquisition. **(G)** Average PSTHs of immobility in CS+ (red) and CS-(black) trials across acquisition. As learning progresses, the difference between CS+ and CS-immobility becomes more pronounced during the early portion of the cue period. **(H)** Selectivity index quantifying the difference in immobility between CS+ and CS-during early (0-10 s) vs. late (19-29 s) cue periods on the last 3 days of Acquisition (Day 9-11). Gray lines indicate individual animals. Error bar shows SEM across animals. **(I)** Human recreational cocaine users ingested cocaine in an atypical environment, i.e., a hospital research setting such as a brain imaging unit. During a follow-up interview, they reported their cocaine use during the preceding 12 months.

**fig. S12.**
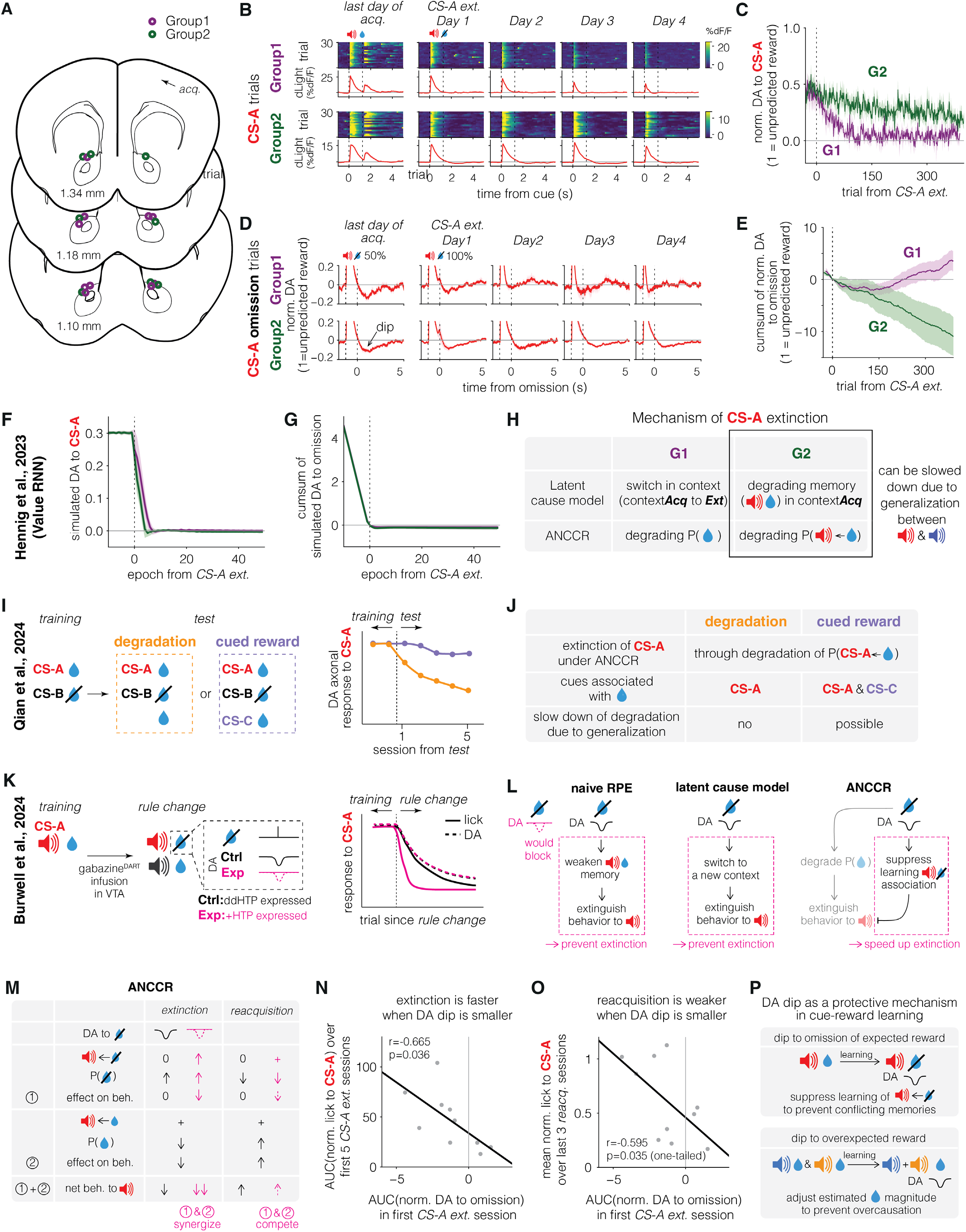
Potential mechanisms underlying dopamine dynamics during extinction. **(A-E)** Changes in dopamine response associated with extinction are slower in G2 than G1 during CS-A extinction. **(A)** Histological locations of optic fiber tips for animals from dopamine fiber photometry experiments. **(B)** Example heatmaps (top) and PSTHs (bottom) of dopamine release (dF/F) in the nucleus accumbens in response to CS-A, from a representative mouse each in G1 and G2. The graphs are aligned to the cue onset. The first column shows data from the last day of acquisition, and the remaining columns show Days 1-4 of CS-A extinction. **(C)** Group-averaged normalized DA to CS-A over trials during CS-A extinction. Lines indicate mean across animals and shadings represent SEM. **(D)** Group-averaged PSTHs for CS-A omission trials over days. Time zero indicates the expected reward delivery time. During acquisition, omission occurred in 50% of CS-A trials; during extinction (Day 1 onward), all CS-A trials were unrewarded. G1 shows a pronounced DA dip that rapidly disappears, while G2 shows a prolonged dip across days. **(E)** Group-averaged cumulative sum of normalized DA to omission across CS-A extinction trials. DA to omission was calculated as AUC of dF/F over a 1.25 s window after expected reward time. **(F-G)** Value RNN, a model for unbiased and automatic discovery of state spaces, does not capture group differences in dopamine extinction. **(F)** Simulated DA (RPE) to CS-A over epochs (see Methods) during CS-A extinction. Lines indicate mean across iterations and shading represent STD. The model predicts a similar dopamine extinction profile across groups. **(G)** Cumulative sum of simulated DA to omission. Again, no group differences are observed. **(H)** Summary of hypothesized mechanisms supporting group differences in CS-A extinction. Neither latent cause model nor ANCCR can account for slower extinction in G2 than G1 without additional assumptions. Assuming generalization between CS-A and CS-B can rescue both models. **(I-J)** Findings from Qian et al. can be explained by ANCCR by assuming generalization. **(I)** Task schematic and behavioral results. Animals that associated a single cue (CS-A) with reward during training received ITI reward either without any cue (degradation) or with a new cue (cued reward) during test. Dopamine response to CS-A decreased more slowly when ITI reward was cued. **(J)** ANCCR with generalization explains the behavior results, as when a new cue is introduced in cued reward condition, generalization between the existing cue (CS-A) and the new cue provides a protective mechanism that slows degradation of CS-A responses. **(K-L)** Findings from Burwell et al. could be explained by ANCCR. **(K)** Task schematic and behavioral result. Animals were trained to associate CS-A with reward. During the rule change phase, CS-A was no longer rewarded while a new cue was paired with reward. Experimental group (Exp), in which dopamine dip at omission was blocked, exhibited faster extinction of anticipatory lick, but not dopamine response to CS-A, compared to control group (Ctrl). Dopamine response to CS-A was not different between groups. **(L)** Summary of how dopamine dip at omission affects extinction in different models. In RPE-based theories (naïve RPE and latent cause model), extinction is triggered by a dip in dopamine at omission. In contrast, ANCCR posits that extinction arises from degradation of P(reward), independent of dopamine at omission. Instead, dopamine dip may reduce the estimated “meaningfulness” of omission event, weakening learning of a cue-omission (i.e., frustration) association. Thus, blocking the dopamine dip may paradoxically enhance learning of cue-omission association and speed up extinction. That said, the dynamics of behavior and dopamine are predicted to be dissociable. For conditioned behavior, the cue-reward association must contribute positively, whereas the cue-omission association must contribute negatively, because predicting reward promotes conditioned responding while predicting omission suppresses it. However, because dopamine response to an event reflects its meaningfulness, both cue-reward and cue-omission associations may contribute positively, as both render the cue informative about future events. Further, because omission (i.e., lack of reward) is meaningful only when reward is expected, strengthening the cue-omission association may have only a small contribution to cue-evoked dopamine, allowing the meaningfulness of the cue to remain primarily determined by the cue-reward association. This is likely because otherwise the cue-omission association would continue to maintain the cue’s meaningfulness even after the cue-reward association had been degraded, i.e., when omission is occurring in the absence of a reward prediction. Once the cue no longer predicts reward, omission should no longer constitute a meaningful event and therefore should not contribute to the cue’s meaningfulness. The observed data support this idea. **(M-O)** Preliminary results support the proposed role of dopamine dip in cue-omission learning in ANCCR. **(M)** Schematic illustrating how dopamine dip at omission operates in ANCCR. A negative response at omission prevents learning of cue←omission (i.e., cue←frustration) association by reducing “meaningfulness” of omission event. This mechanism ensures that animals do not form a separate cue-omission memory on top of the existing cue-reward memory, as recognition of omission itself depends on prior learning of the cue-reward contingency and cannot be treated as an independent event. ANCCR predicts that abolishing the dopamine dip allows learning of a cue-omission association, which facilitates behavioral extinction by introducing additional cue-omission (i.e., frustration) memory alongside the decrease in estimated reward rate. However, this newly formed cue-omission memory competes with the preserved cue-reward memory during reacquisition, resulting in slower relearning. **(N)** Correlation between magnitude of dopamine dip at omission on the first day of CS-A extinction and speed of CS-A extinction across animals. Across animals, a larger dopamine dip at omission on the first day of CS-A extinction correlated with slower extinction, consistent with ANCCR’s prediction that cue-omission learning facilitates extinction. **(O)** Correlation between magnitude of dopamine dip at omission on the first day of CS-A extinction and asymptotic level of reacquired behavior to CS-A. Animals with smaller dopamine dip at omission showed weaker asymptotic levels of reacquired behavior to CS-A, supporting the prediction that cue-omission memory competes with cue-reward memory during reacquisition. **(P)** Dopamine dip acts as a protective mechanism in cue-reward learning. During omission of an expected reward, dopamine dip prevents maladaptive learning by suppressing cue←omission association that could compete with existing cue←reward memory. During overexpectation, dopamine dip adjusts the estimated reward magnitude associated with each cue to prevent overexpectation. Although the omission-evoked dip may also reduce the perceived magnitude of the omission (i.e., magnitude of frustration) itself, this has little impact because cue←omission learning is simultaneously suppressed. Together, these mechanisms stabilize learning by preventing conflicting or excessive associations.

**fig. S13.**
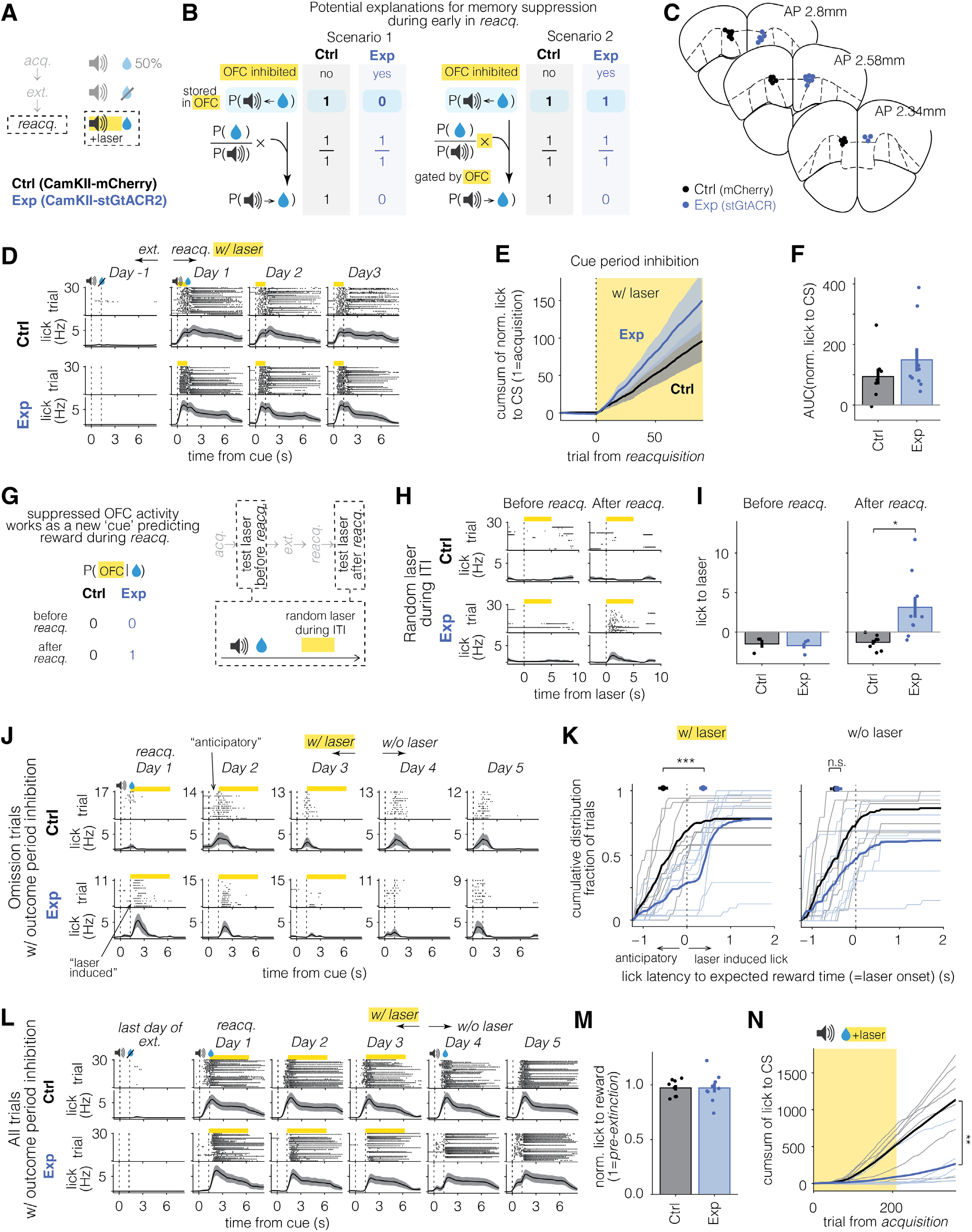
Suppressed OFC activity during cue works as a new cue predicting reward. **(A)** Experimental schematic. During reacquisition following standard extinction, OFC CamKIIα-expressing neurons were optogenetically suppressed for 1.25 s from cue onset to reward in experimental group (Exp). The control group (Ctrl) underwent the same procedure without expression of inhibitory opsin. **(B)** Two potential explanations for memory suppression via OFC inactivation. If retrospective association between cue and reward is stored in OFC, suppression of the memory can degrade reacquisition (scenario 1). Or if computation of prospective association is gated by OFC, preventing it would also result in a deficit in reacquisition (scenario 2). **(C)** Histological location of optic fiber centers for OFC optogenetic experiments. **(D)** Example lick raster plots and PSTHs for a representative mouse per group (top, Ctrl; bottom, Exp). Data are shown for the last day of extinction and the first three days of reacquisition (Day 1 to 3). Yellow bars indicate laser-on periods. Both Ctrl and Exp animals reacquire anticipatory lick on Day 1. **(E)** Group-averaged cumulative sum of normalized lick to CS. Yellow shading indicates trials with laser on. Both groups show rapid reacquisition compared to acquisition. **(F)** AUC of normalized lick to CS during reacquisition. OFC inhibition during the cue period did not slow reacquisition; if anything, Exp showed a non-significant tendency toward more licking, consistent with laser-evoked licking (**H–K**). **(G)** Experimental schematic to test if suppression of OFC works as a new cue. A 5 s long laser was randomly delivered during ITI before and after reacquisition to see if laser induces licking. During reacquisition, laser was delivered during cue-reward delay, allowing it to be associated with following reward. **(H)** Example lick raster plots and PSTHs for example animal per group in random laser test before (left) and after (right) reacquisition. In Exp, delivery of laser induces licking only after reacquisition, where laser was followed by reward, indicating that the suppressed activity of OFC induced by the laser works as a new cue predicting reward. **(I)** Comparison of lick to laser between groups before (left) and after (right) reacquisition. **(J)** Example lick raster plots and PSTHs for one representative animal per group, showing omission trials during reacquisition with outcome period inhibition. Yellow bars indicate laser on periods. In Ctrl, anticipatory lick appears by Day 2. However, Exp animal shows lick immediately following laser onset on Day 1. This laser-induced lick diminishes from Day 1 to Day 3, consistent with extinction of laser-reward association. When laser was omitted on Day 4, Exp animal begins to show anticipatory lick. **(K)** Cumulative distribution of first lick latency during omission trials, aligned to the expected reward time (and laser onset). Light traces represent individual animals. Bold traces show group averages. Ctrl shows anticipatory lick regardless of laser condition (left, with laser; right, without laser). Exp shows strong laser-induced lick in sessions with laser (left), which disappears when the laser is omitted (right). Horizontal error bar indicates group average of mean lick latency. **(L)** Example lick raster plots (top row) and PSTHs (bottom row) for one mouse per group (top, Ctrl; bottom, Exp). Data are shown across days including the last day of extinction and the first five days of reacquisition (Day 1 to 5). Laser was delivered on the first three days of reacquisition (Day 1 to 3). Yellow bars indicate laser-on periods. Ctrl shows rapid reacquisition on Day 1, while Exp fails to reacquire for all three days. When laser was turned off on Day4, anticipatory lick in Exp emerged, indicating the delay in reacquisition was due to suppression of OFC reward responses. **(M)** Normalized lick to reward does not differ between groups, suggesting that OFC suppression does not impair motor ability or motivation to reward. **(N)** Group-averaged cumulative sum of normalized lick to CS during acquisition when OFC reward response was inhibited (Exp in blue) or not (Ctrl in black). *, AUC of anticipatory licking across trials during acquisition with laser on was significantly lower when OFC reward response was inhibited than when it was not.

**fig. S14.**
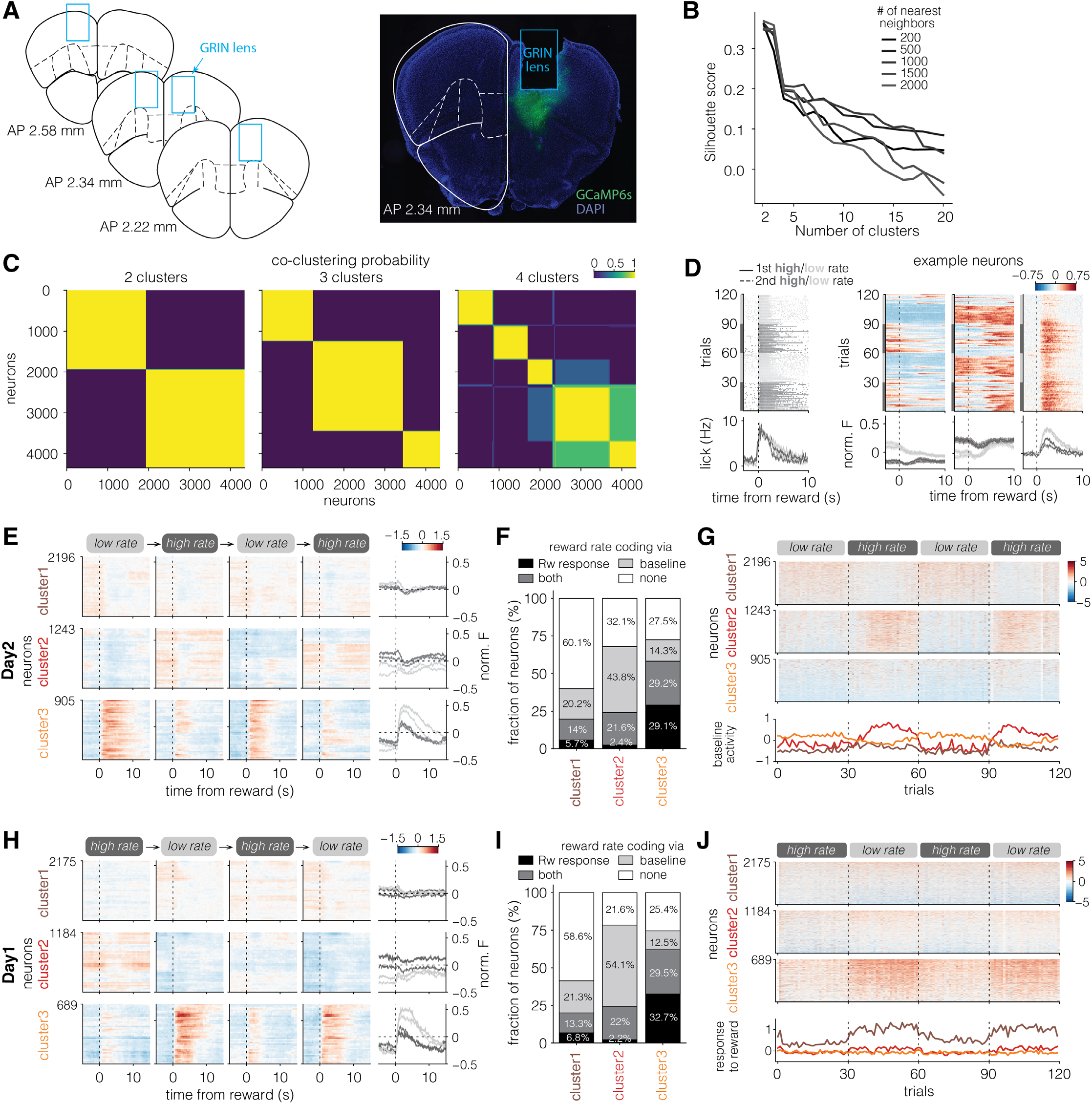
Reward rate coding by OFC neurons was consistently observed across days. **(A)** Histological location of the GRIN lens (left) and representative histological images (right). The left panel shows the position of the lens bottom centered along the anterior-posterior axis. **(B)** Silhouette score results from spectral clustering using varying number of clusters (k) and nearest neighbors. Both k = 2 or 3 produced similarly high silhouette scores, indicating that neurons within each cluster exhibit cohesive response patterns distinct from those in other clusters, regardless of number of nearest neighbors. **(C)** Co-clustering probability across 100 independent clustering iterations with different random seeds, indicating the stability of clustering results. Again, both k = 2 or 3 yielded highly stable clustering results. As k = 2 and 3 did not differ substantially in clustering validity (silhouette score) or stability (co-clustering probability), we selected k = 3 to allow finer subdivision of neuronal response patterns. **(D)** Lick rasters and PSTHs from a representative mouse (left) and heatmaps and PSTHs for five example neurons (right). Dark and light gray indicate low and high rate blocks, respectively. Solid line corresponds to the first low/high blocks (first and second blocks of the session), and dashed lines to the second low/high blocks (third and fourth blocks of the session). Time 0 marks the first lick after reward delivery. In case animal does not lick within 5 s after reward delivery, the trial was excluded from analysis. **(E)** Heatmaps and PSTHs of clustered neurons (see Methods) averaged across the last three trials of each block. Neurons are sorted by average reward response across blocks. **(F)** Fraction of neurons encoding reward rate via changes in phasic reward response, tonic baseline activity, or both, shown for each cluster. **(G)** Same as in Fig. 5H but for baseline responses. **(H-J)** Same analyses as in **fig. S14E,F and** Fig. 5H, but from the first recording day in which the order of reward-rate blocks was reversed. Clustering was performed separately for each day and yielded similar cluster structures. **(H)** Heatmaps and PSTHs of clustered neurons averaged across the last three trials of each block. Note that the first block (high rate) shows more distinct activity pattern from the second high rate block compared with the nearly identical patterns observed across blocks of the same reward rate on Day 2 (**fig. S14E**), particularly for cluster 2, whose baseline activity encodes reward rate. Because animals had not experienced sucrose rewards or the experimental context (e.g., head fixation box) before Day 1, the distinct activity in the first block likely reflects novelty-related effects in addition to reward rate. **(I)** Fraction of neurons encoding reward rate via changes in phasic reward response, tonic baseline activity, or both, shown for each cluster. **(J)** Reward responses of individual neurons across trials (top) and averaged response across neurons within each cluster (bottom).

**fig. S15.**
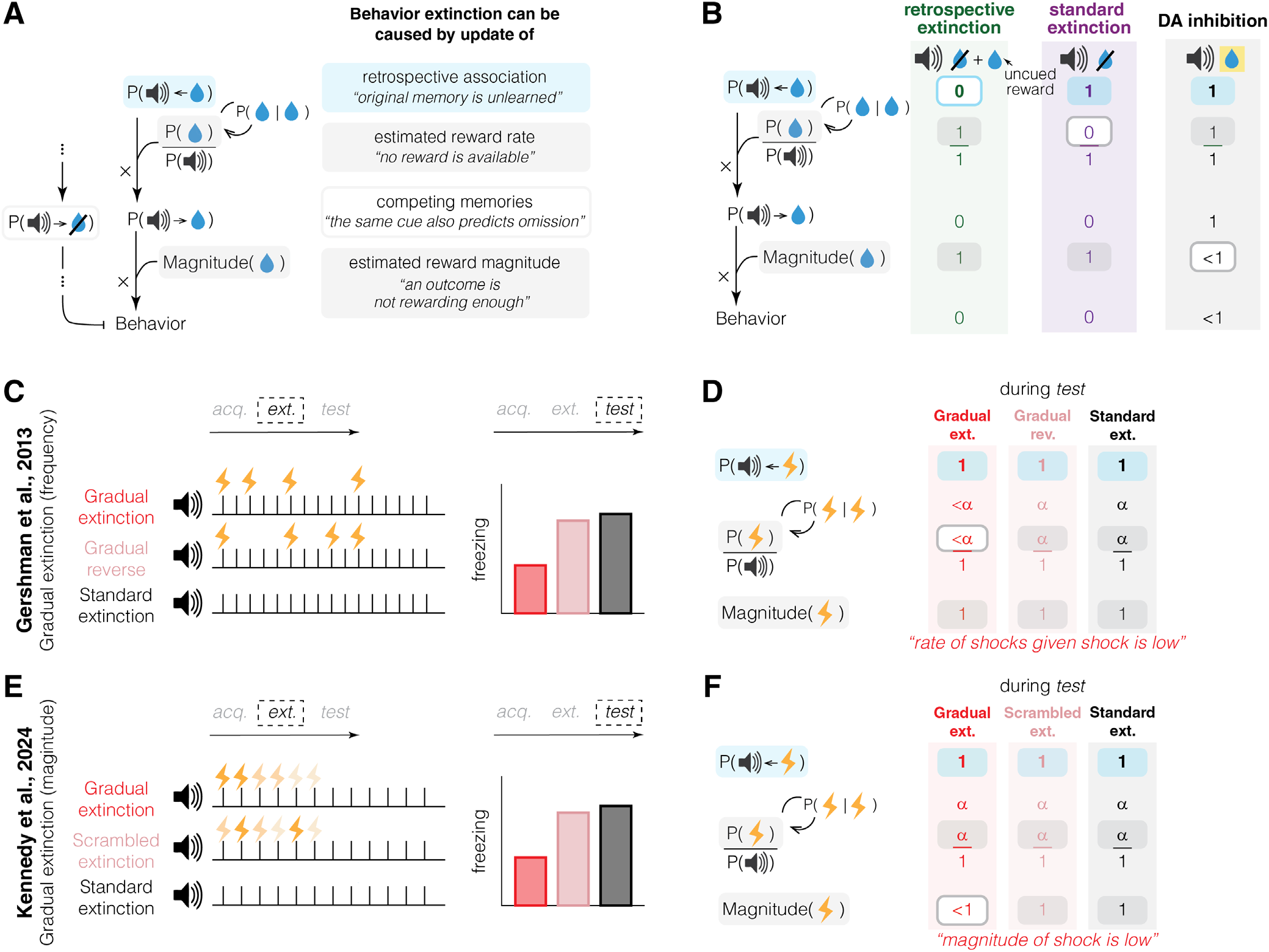
Low estimates of outcome rate or magnitude can induce behavior extinction despite preserved memory. **(A-B)** ANCCR predicts that degradation of three distinct computational variables can induce behavior extinction (filled box in A). Extinction via degraded retrospective association (i.e., retrospective extinction; left in B) results in unlearning of the original memory and leads to slow reacquisition. In contrast, extinction caused by reduced estimated reward rate (e.g., standard extinction; middle in B) or reward magnitude (e.g., DA inhibition; right in B) occurs despite the original memory being intact. Degraded component is marked as dark edge around the shaded area in B. Additionally, ANCCR predicts that strengthening of a competing memory (empty box in A), such as cue←omission association, can also induce behavior extinction, as observed in Burwell et al. (**fig. S12K-P**). This mechanism is separate from degradation of the computational variables underlying the cue←reward association. **(C-D)** Extinction through gradual reduction in frequency of shock can be explained by ANCCR. **(C)** Experimental design and finding from Gershman et al. (2013). When shock frequency gradually decreased during extinction (Gradual extinction), retention of behavior was weaker on following test than in Gradual reverse (increasing frequency) or Standard extinction (no shocks during extinction). **(D)** ANCCR prediction. We hypothesized that animals estimate the rate of outcome given outcome (i.e., the expected rate of outcomes following a single outcome) and this prior can facilitate rapid reacquisition of behavior when the original memory is intact. This rate is only updated at outcome delivery. In Gradual extinction, the final shock occurs after a long gap since the previous one, reducing the estimated rate of shock given shock. In contrast, in Gradual reverse, the final shock is preceded by another shock after a shorter interval, preserving a higher rate. In Standard extinction, the last shock is experienced during acquisition, when shocks were frequent, so the rate of shock given shock remains high. Thus, only in Gradual extinction, animals show lower level of behavior. **(E-F)** Extinction through gradual reduction in magnitude of shock can be explained by ANCCR. **(E)** Experimental design and finding from Kennedy et al., (2024). In Gradual extinction, shock magnitude decreased gradually over trials during extinction. In Scrambled extinction, shock magnitude was presented in a randomized order during extinction. Standard extinction involved no shocks during extinction. Weaker freezing was observed in Gradual extinction compared to other conditions during test. **(F)** ANCCR prediction. In Gradual extinction, estimated magnitude of shock decreased over time, leading animals to infer that the shock associated with cue is weak. In Scrambled extinction, stronger shocks may occur closer to the test, resulting in a higher estimated magnitude than Gradual extinction. In Standard extinction, the magnitude estimate remains as high as during acquisition.

**fig. S16.**
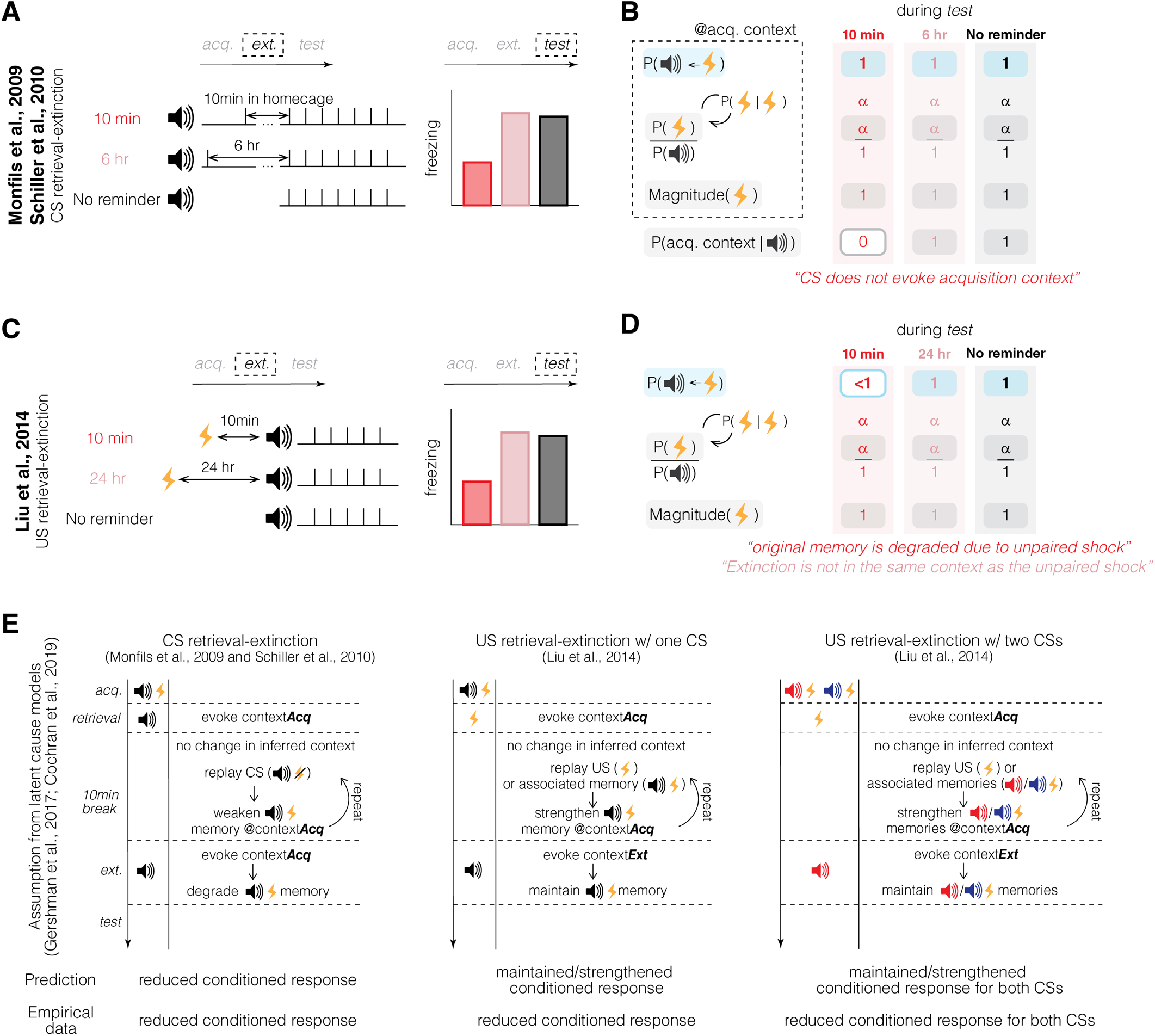
Potential mechanisms to explain retrieval-extinction reconsolidation results. **(A-B)** Conditioned stimulus (CS) retrieval-extinction. **(A)** Experimental schematic and findings from Monfils & Schiller. When a retrieval cue is presented shortly before standard extinction (10 min delay), recovery of conditioned behavior is suppressed. However, the effect disappears when the delay is extended to 6 hours. **(B)** Potential explanation under ANCCR framework. A context is nothing more than a set of cues. Therefore, animals may infer a context change when the properties of CSs change. In the 10 min condition, the unusually long delay between the retrieval trial and the first extinction trial (compared to ITIs during acquisition) may lead animals to infer a new context. Note that the retrieval trial is not operationally different from extinction trials. Because this new context was never paired with cue-shock memory, conditioned response is low at test. In contrast, when the delay is extended to 6 hours, animals may treat the retrieval trial and extinction session as separate episodes (e.g., different experimental sessions), thus avoiding the context shift during extinction. **(C-D)** Unconditioned stimulus (US) retrieval-extinction. **(C)** Experimental schematic and findings from Liu et al. (2014). Similar schematic to CS retrieval-extinction, but use US as a retrieval instead of CS. Having retrieval US 10 min before extinction can reduce recovery of behavior at test compared to when delay is long (24 hr) or no US was given (No reminder). **(D)** Potential explanation under ANCCR framework. In the 10 min condition, the retrieval US (unpaired shock) may degrade the retrospective association between cue and shock, weakening the original memory. This reduction could be facilitated by replay of unpaired US during 10 min delay. Supporting this idea, the same study showed that US retrieval followed by extinction of one cue reduced conditioned behavior to a different cue that had also been paired with the same shock. This suggests that memory degradation stems from the unpaired shock itself, rather than the extinction procedure. This is consistent with the ANCCR prediction that an unpaired outcome globally weakens retrospective association with its predictive cues. When the delay from US retrieval to extinction is long (24 hr), animals may fail to assign the US to the same context as the rest of experiment, preventing degradation of the retrospective association. A similar idea may explain inter-animal variability in G4 (**fig. S8**), where some animals showed no deficit in reacquisition after unpaired reward-only sessions. **(E)** Prediction from latent cause models. In CS retrieval-extinction (left), latent cause models assume that the retrieval trial evokes inference of the acquisition context (contextAcq), and that rumination of retrieval trial (CS alone), which is assumed to not alter the inferred context but produce associative learning, during the 10 min delay weakens the cue-reward association at contextAcq. This reduces RPE during subsequent extinction trial, preventing context switching and resulting in unlearning. When the delay is long (6 hr), the model infers a new context due to a time-based prior favoring different contexts for temporally distant events, thereby preserving the original memory. However, latent cause models cannot account for the effects of US retrieval-extinction (middle and right), since replaying US alone or the CS-US association would not weaken cue-reward memory under the RPE framework (*38*). Thus, extinction trials following US retrieval should evoke a new context and preserve the original memory, contradicting the empirical findings.

**fig. S17.**
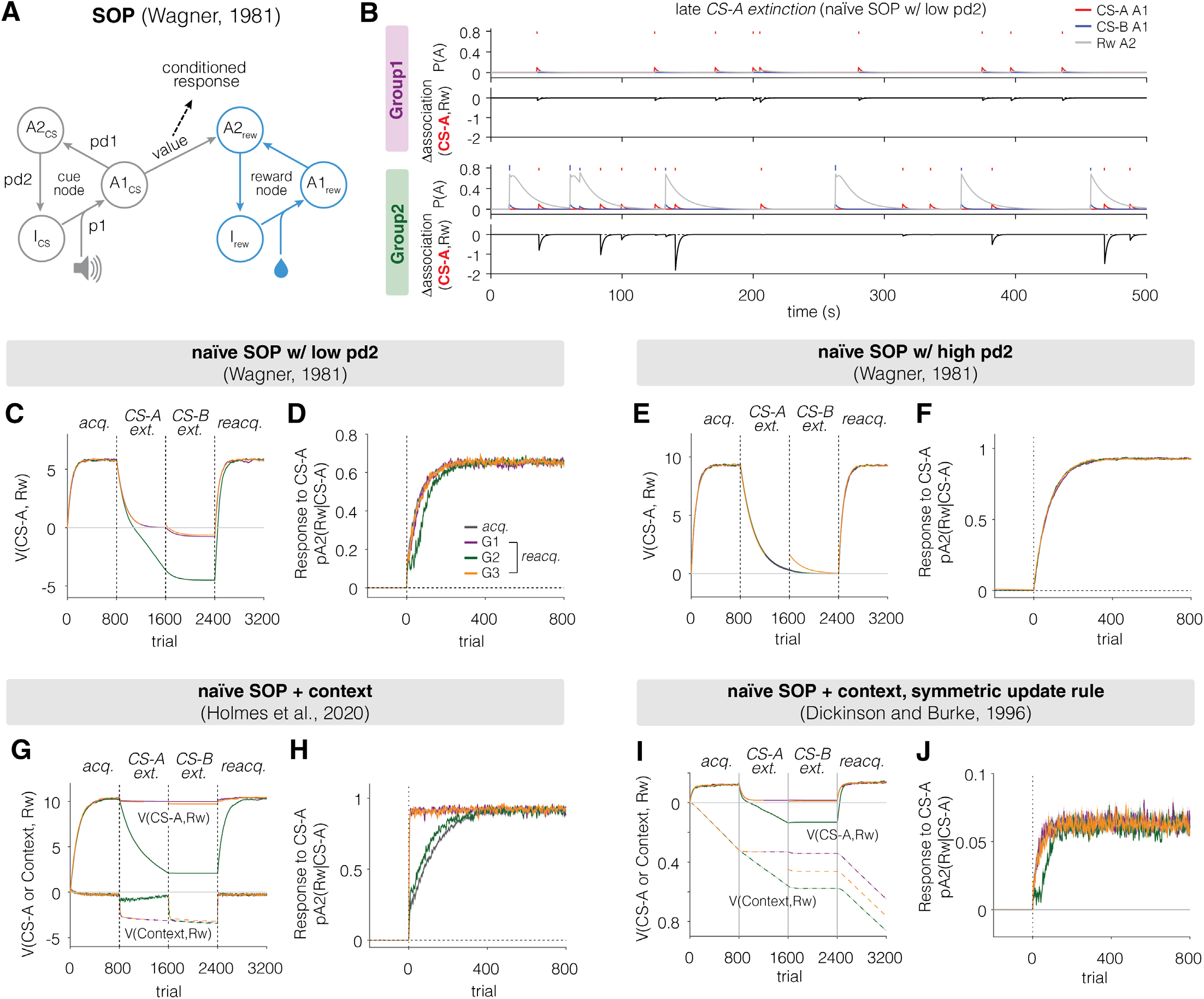
Simulation results from Wagner’s SOP model. **(A)** Schematic of Wagner’s SOP (Standard Operating Procedure or Sometimes Opponent Process; (*95*)) model. Stimulus presentations evoke mental representations composed of multiple informational elements. These stimulus representations are dynamic: presentation of a stimulus moves a portion of elements from the inactive (I) state into the primary active state (A1), then decay to the secondary active state (A2), and finally return to I. Transition of elements between states is determined by specified probabilities (p1, pd1, pd2). Cue-reward association (value) is strengthened when cue and reward elements are simultaneously in A1, and weakened when cue elements in A1 (A1_CS_) overlap with reward elements in A2 (A2_rew_). A2rew can be also evoked by A1CS, and the strength depends on the cue-reward association. **(B-F)** Simulation results from the naïve SOP. During early CS-A extinction, presentation of CS-A without reward induces overlap between A1CS-A and A2rew, causing CS-A value to reduce. Once CS-A value decreases sufficiently, CS-A no longer evokes A2_rew_ (red, B), leading to saturation of CS-A value in Group1 (C). In Group2, however, CS-B reward trials evoke A2_rew_ (grey, B), which can persist into subsequent CS-A trials. As a result, late in CS-A extinction (example traces shown in B) once the association of CS-A and reward decays enough, CS-A presentation (red ticks) stops evoking A2_rew_ (grey) as shown in example traces in both groups. In Group1, this causes a saturation in value function of CS-A (C, G1 and G3) as A1_CS-A_ no longer overlaps with A2_rew_. However, in Group2, presentation of CS-B and following reward triggers high A2_rew_, and when the transition of A2 elements to I is slow (low pd2; C,D). This residual A2_rew_ overlaps with A1_CS-A_ and drives CS-A value below zero, producing conditioned inhibition and slowing reacquisition. This effect disappears when transition of A2 elements to I is fast enough, because A2_rew_ decays before the next CS-A trial (E,F). Although SOP model can predict slower extinction in G2 than G1 in a parameter-dependent manner, it does not capture G3 result, because separating CS-A and CS-B trials into different blocks prevent overlap between CS-B evoked A2_rew_ and A1_CS-A_. **(G-J)** Simulation results from modified SOP models. Neither modeling context as an external cue (G,H) (*96*) nor generalizing the value-update rule (I,J) (*97*)—such that associations are strengthened when cue and reward are in the same state (A1 or A2) and weakened otherwise (either cue in A1 and reward in A2 as in original SOP, or the other way)—captured G3 results.

## Supplementary Tables

**Supplementary Table 1.**
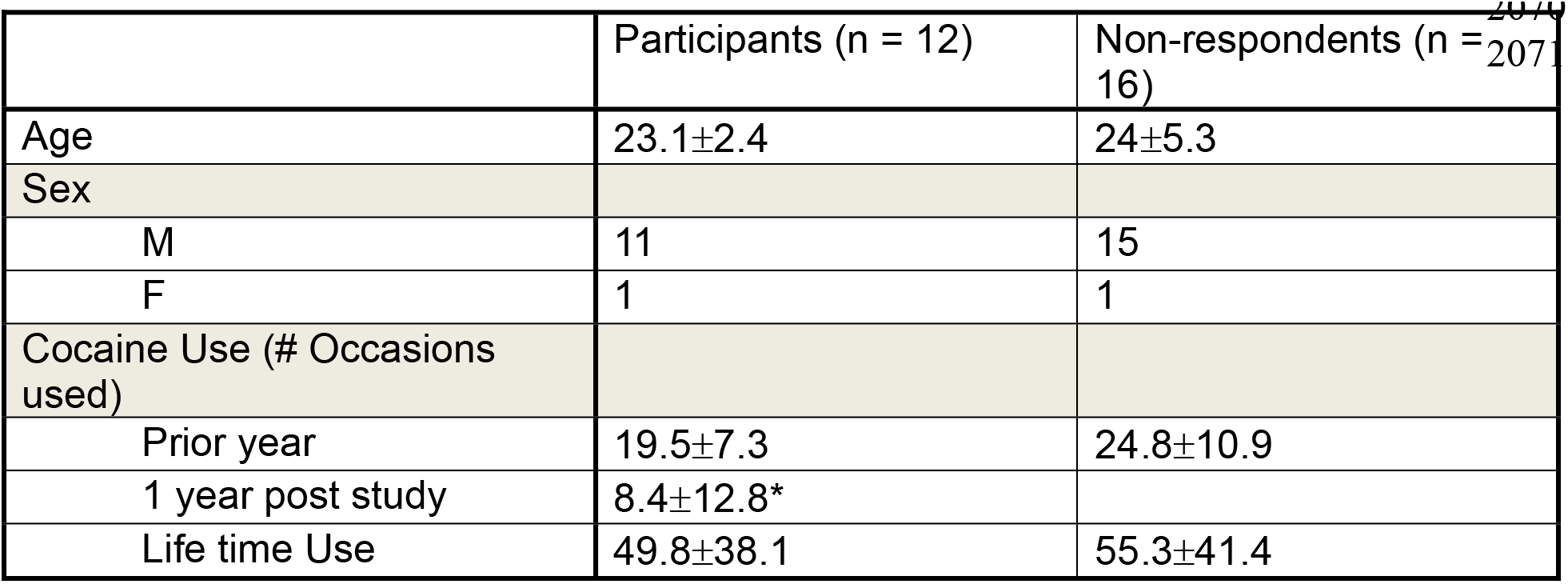
Demographics and cocaine use patterns (mean±STD) of participants. *Three out of 12 participants had not used cocaine during the year post study. Mean number of occasions used was based on all 12 participants.

**Supplementary Table 2.** Statistical results. Mean differences were calculated as [first condition] minus [second condition], with 95% confidence intervals. Effect sizes are Cohen’s d for independent comparisons and Cohen’s dᶻ for paired comparisons. All independent-samples t-tests were Welch’s tests. Tests were two-tailed unless a directional hypothesis based on previous findings justified a one-tailed test, as indicated in the P value column.

| Figure | Description | Test | Statistic, t (df) | Mean difference [95% CI] | Effect size, Cohen's d | p value | Number of samples |
| --- | --- | --- | --- | --- | --- | --- | --- |
| Fig. 1I | AUC(norm lick to CS-A); acq. vs. reacq. | Paired t test | G1, -7.562 (21);<br>G2, 2.865 (18) | G1, -42.82 [-54.60, -31.05];<br>G2, 20.38 [5.44, 35.32] | G1, -1.612;<br>G2, 0.657 | Two-tailed p value =<br>G1, 2.0×10<sup>-7</sup>;<br>G2, 0.010 | G1, 22;<br>G2, 19 |
| Fig. 1L | trials to learn CS-A; learners only; acq. vs. reacq. | Two sample t test | G1, 6.500 (39.04);<br>G2, -0.114 (23.16) | G1, 21.91 [15.09, 28.73];<br>G2, -1.07 [-20.33, 18.20] | G1, 1.960;<br>G2, -0.042 | Two-tailed p value =<br>G1, 1.0×10<sup>-7</sup>;<br>G2, 0.910 | G1, [22, 22];<br>G2, [19, 12] |
| Fig. 1N | AUC(norm lick to CS-A); G1 vs. G2 | Two sample t test | 3.483 (21.39) | 1.10 [0.44, 1.75] | 1.013 | Two-tailed p value = 0.002 | G1, 22;<br>G2, 19 |
| Fig. 1Q | AUC(norm DA to CS-A); acq. vs. reacq. | Paired t test | G1, -5.154 (9);<br>G2, -0.408 (7) | G1, -17.72 [-25.49, -9.94];<br>G2, -2.07 [-14.09, 9.94] | G1, -1.630;<br>G2, -0.144 | Two-tailed p value =<br>G1, 6.0×10<sup>-4</sup>;<br>G2, 0.696 | G1, 10;<br>G2, 8 |
| Fig. 2E | AUC(norm lick to CS); acq. vs. reacq. | Paired t test | Ctrl, 1.182 (8) ;<br>Exp, -3.371 (8) | Ctrl, 7.26 [-6.90, 21.42];<br>Exp, -24.83 [-41.82, -7.84] | Ctrl, 0.394;<br>Exp, -1.124 | Two-tailed p value =<br>Ctrl, 0.2710 ;<br>Exp, 9.8×10<sup>-4</sup> | Ctrl, 9;<br>Exp, 9 |
| Fig. 2F | AUC(norm lick to CS-A) | Two sample t test with Bonferroni correction | G1 vs. G2, 8.864 (37.99);<br>G1 vs. Ctrl, 8.640 (22.89);<br>G1 vs. Exp, 1.085 (16.96);<br>G2 vs. Ctrl, -0.543 (23.33);<br>G2 vs. Exp, -6.438 (18.11);<br>Ctrl vs. Exp, -6.139 (14.74) | G1 vs. G2, 60.67 [46.82, 74.53];<br>G1 vs. Ctrl, 56.93 [43.30, 70.57];<br>G1 vs. Exp, 8.52 [-8.04, 25.08];<br>G2 vs. Ctrl, -3.74 [-17.97, 10.50];<br>G2 vs. Exp, -52.15 [-69.17, -35.14];<br>Ctrl vs. Exp, -48.42 [-65.25, -31.58] | G1 vs. G2, 2.779;<br>G1 vs. Ctrl, 2.861;<br>G1 vs. Exp, 0.406;<br>G2 vs. Ctrl, -0.188;<br>G2 vs. Exp, -2.470;<br>Ctrl vs. Exp, -2.894 | Two-tailed p value =<br>G1 vs. G2, 5.2×10<sup>-10</sup>;<br>G1 vs. Ctrl, 7.0×10<sup>-8</sup>;<br>G1 vs. Exp, 1;<br>G2 vs. Ctrl, 1;<br>G2 vs. Exp, 2.7×10<sup>-5</sup>;<br>Ctrl vs. Exp, 1.2×10<sup>-4</sup> | G1, 22;<br>G2, 19;<br>Ctrl, 9;<br>Exp, 9 |
| Fig. 2G | Norm consummatory lick; Ctrl vs. Exp | Two sample t test | -1.609 (10.27) | -0.10, [-0.25, 0.04] | -0.759 | Two-tailed p value = 0.138 | Ctrl, 9;<br>Exp, 9 |
| Fig. 3L | AUC(norm lick to CS); reacquisition; CS-A vs. CS-B | Paired t test | G1, -2.077 (21);<br>G2, -5.100 (18);<br>G3, -2.742 (8) | G1, -6.21 [-12.43, 0.01];<br>G2, -43.35 [-61.20, -25.49];<br>G3, -27.51 [-50.65, -4.37] | G1, -0.443;<br>G2, -1.170;<br>G3, -0.914 | Two-tailed p value =<br>G1, 0.0502;<br>G2, 7.5×10<sup>-5</sup>;<br>G3, 0.0254 | G1, 22;<br>G2, 19;<br>G3, 9 |
| Fig. 3L-stat only | Difference in AUC(norm lick to CS) during reacquisition between CS-A and CS-B | Two sample t test | G1 vs. G2, -4.122 (22.44)<br>G1 vs. G3, -2.034 (9.46) | G1 vs. G2, -37.14 [-inf, -21.68];<br>G1 vs. G3, -21.30 [-inf, -2.21] | G1 vs. G2, -1.366;<br>G1 vs. G3, -1.075 | One-tailed p value = G1 vs. G2, $2.2 \times 10^{-4}$;<br>G1 vs. G3, 0.0355; testing if cue difference in G1 is smaller than that in G2 or G3 based on results of Fig. 3L | G1, 22;<br>G2, 19; |
| Fig. 4C | AUC(lick to CS+) | Two sample t test | 3.877 (6.82) | 176.71, [68.35, 285.08] | 2.073 | Two-tailed p value = 0.006 | G1, 7;<br>G2, 7 |
| Fig. 4F | AUC(selectivity) | Paired t test | G1, -3.990 (6);<br>G2, -0.406 (6) | G1, -14.59 [-23.54, -5.64];<br>G2, -0.84 [-5.89, 4.22] | G1, -1.508;<br>G2, -0.154 | Two-tailed p value = G1, 0.007;<br>G2, 0.699 | G1, 7;<br>G2, 7 |
| Fig. 5C | AUC(norm. lick to CS) | Two sample t test | 4.273 (14.73) | 47.17, [23.60, 70.74] | 2.078 | Two-tailed p value = $6.9 \times 10^{-3}$ | G1, 8;<br>G2, 9 |
| Fig. S2B | Averaged lick to CS-A after learning; learners only; acq. vs. reacq. | Two sample t test | G1, -1.269 (41.76);<br>G2, 2.463 (24.58) | G1, -0.50 [-1.30, 0.30];<br>G2, 1.08 [0.18, 1.98] | G1, -0.383;<br>G2, 0.896 | Two-tailed p value = G1, 0.211;<br>G2, 0.021 | G1, [22, 22];<br>G2, [19, 12] |
| Fig. S2C | Averaged lick to CS-A | Two sample t test | 1.153 (36.97) | 0.06 [-0.04, 0.16] | 0.351 | Two-tailed p value = 0.256 | G1, 22;<br>G2, 19 |
| fig. S2D | Averaged lick to CS-A | Two sample t test | 0.830 (36.27) | 0.02 [-0.03, 0.07] | 0.262 | Two-tailed p value = 0.412 | G1, 22;<br>G2, 19 |
| fig. S2G, right | AUC(norm lick to CS-A); session number matched; acq. vs reacq. | Paired t test | G1, -6.130 (11);<br>G2, 0.990 (10) | G1, -50.86 [-69.12, -32.60];<br>G2, 10.58 [-13.23, 34.39] | G1, -1.770;<br>G2, 0.299 | Two-tailed p value = G1, $7.4 \times 10^{-5}$;<br>G2, 0.345 | G1, 12;<br>G2, 11 |
| fig. S2H, right | AUC(norm lick to CS-A); behavior matched; acq. vs reacq. | Paired t test | G1, -4.961 (9);<br>G2, 5.360 (7) | G1, -33.19 [-48.32, -18.05];<br>G2, 33.85 [18.92, 48.78] | G1, -1.569;<br>G2, 1.895 | Two-tailed p value = G1, $7.8 \times 10^{-4}$;<br>G2, 0.001 | G1, 10;<br>G2, 8 |
| fig. S2J | AUC(norm lick to CS-B); acq. vs. reacq. | Paired t test | G1, -7.972 (21);<br>G2, -1.663 (18);<br>With analysis window restricted to the first 2 days:<br>G1, -7.840 (21);<br>G2, -2.305 (18) | G1, -52.01 [-65.58, -38.44];<br>G2, -20.26 [-45.86, 5.34];<br>With analysis window restricted to the first 2 days:<br>G1, -35.99 [-45.54, -26.45];<br>G2, -18.45 [-35.26, -1.63] | G1, -1.700;<br>G2, -0.381;<br>With analysis window restricted to the first 2 days:<br>G1, -1.672;<br>G2, -0.529 | Two-tailed p value = G1, $8.7 \times 10^{-8}$;<br>G2, 0.114;<br>With analysis window restricted to the first 2 days:<br>G1, $1.1 \times 10^{-8}$;<br>G2, 0.033 | G1, 22;<br>G2, 19 |
| fig. S2K | trials to learn CS-B; learners | Two sample t test | G1, 5.827 (41.27) ;<br>G2, 1.433 (32.28) | G1, 21.73 [14.20, 29.26]; | G1, 1.757;<br>G2, 0.482 | Two-tailed p value = G1, $7.5 \times 10^{-7}$; | G1, [22, 22]; |
|  | only; acq.<br>vs. reacq. |  |  | G2, 14.52 [-6.11, 35.15] |  | G2, 0.161 | G2, [19, 17] |
| fig. S2M | AUC(norm DA to CS-B); acq. vs. reacq. | Paired t test | G1, -4.635 (9);<br>G2, -2.876 (7) | G1, -20.27 [-30.16, -10.37];<br>G2, -23.41 [-42.66, -4.16] | G1, -1.466;<br>G2, -1.017 | Two-tailed p value =<br>G1, $1.2 \times 10^{-3}$;<br>G2, 0.024 | G1, 10;<br>G2, 8 |
| fig. S3D | AUC(norm lick to CS-A); G1 vs. G2 | Two sample t test | -4.286 (27.06) | -44.60 [-65.95, -23.25] | -1.397 | Two-tailed p value = $2.1 \times 10^{-4}$ | G1, 22;<br>G2, 19 |
| fig. S3H | AUC(norm selectivity); G1 vs. G2 | Two sample t test | 0.748 (10.93) | 0.89 [-1.73, 3.51] | 0.400 | Two-tailed p value = 0.470 | G1, 7;<br>G2, 7 |
| fig. S4D | AUC(norm lick to CS-B); G1 vs. G2 | Two sample t test | -3.273 (24.68) | -35.70 [-58.19, -13.22] | -1.076 | Two-tailed p value = 0.003 | G1, 22;<br>G2, 19 |
| fig. S4G | AUC(norm DA to CS-B); G1 vs. G2 | Two sample t test | -4.829 (9.94) | -33.90 [-49.55, -18.24] | -2.454 | Two-tailed p value = $7.1 \times 10^{-4}$ | G1, 10;<br>G2, 8 |
| fig. S4K | AUC(norm lick to CS); CS-A vs. CS-B | Paired t test | t = 4.074 (21) | 20.82 [10.19, 31.45] | 0.869 | Two-tailed p value = $5.4 \times 10^{-4}$ | 22 |
| fig. S4N | AUC(norm DA to CS); CS-A vs. CS-B | Paired t test | 1.586 (9) | 7.61 [-3.24, 18.46] | 0.502 | Two-tailed p value = 0.147 | 10 |
| fig. S5C | AUC(norm lick to CS) | Paired t test | Ctrl, CS-A, 1.182 (8);<br>Ctrl, CS-B, -2.444 (8);<br>Exp, CS-A, -3.371 (8);<br>G2, CS-B, -5.837 (8) | Ctrl, CS-A, 7.26 [-6.90, 21.42];<br>Ctrl, CS-B, -47.09 [-91.51, -2.66];<br>Exp, CS-A, -24.83 [-41.82, -7.84];<br>Exp, CS-B, -45.68 [-63.73, -27.63] | Ctrl, CS-A, 0.394;<br>Ctrl, CS-B, -0.815;<br>Exp, CS-A, -1.124;<br>Exp, CS-B, -1.946 | Two-tailed p value =<br>Ctrl, CS-A, 0.2710;<br>Ctrl, CS-B, 0.040;<br>Exp, CS-A, $9.8 \times 10^{-4}$;<br>Exp, CS-B, $3.9 \times 10^{-4}$ | Ctrl, 9;<br>Exp, 9 |
| fig. S5E | AUC(norm DA to CS) | Paired t test | Ctrl, CS-A, -2.562 (4);<br>Ctrl, CS-B, -4.973 (4);<br>Exp, CS-A, 0.044 (4);<br>Exp, CS-B, -1.077 (4) | Ctrl, CS-A, -10.98 [-22.88, 0.92];<br>Ctrl, CS-B, -49.47 [-77.10, -21.85];<br>Exp, CS-A, 0.18 [-11.48, 11.84];<br>Exp, CS-B, -7.58 [-27.11, 11.95] | Ctrl, CS-A, -1.146;<br>Ctrl, CS-B, -2.224;<br>Exp, CS-A, 0.019;<br>Exp, CS-B, -0.482 | Two-tailed p value =<br>Ctrl, CS-A, 0.062;<br>Ctrl, CS-B, 0.008;<br>Exp, CS-A, 0.967;<br>Exp, CS-B, 0.342 | Ctrl, 5;<br>Exp, 5 |
| fig. S5G | AUC(norm DA to CS) | Paired t test | Ctrl, CS-A, -3.650 (4);<br>Ctrl, CS-B, -8.654 (4);<br>Exp, CS-A, -4.906 (4);<br>Exp, CS-B, -8.083 (4) | Ctrl, CS-A, -25.99 [-45.75, -6.22];<br>Ctrl, CS-B, -64.34 [-84.98, -43.70];<br>Exp, CS-A, -14.50 [-22.71, -6.30]; | Ctrl, CS-A, -1.632;<br>Ctrl, CS-B, -3.870;<br>Exp, CS-A, -2.194;<br>Exp, CS-B, -3.615 | Two-tailed p value =<br>Ctrl, CS-A, 0.022;<br>Ctrl, CS-B, $9.8 \times 10^{-4}$;<br>Exp, CS-A, 0.008;<br>Exp, CS-B, 0.001 | Ctrl, 5;<br>Exp, 5 |
|  |  |  |  | Exp, CS-B, -<br>23.19 [-31.16, -<br>15.23] |  |  |  |
| fig. S5L | AUC(norm<br>DA to<br>reward) | Two<br>sample<br>t test | 6.428 (7.45) | 2.09 [1.33, 2.85] | 4.066 | Two-tailed p value<br>= $2.8 \times 10^{-4}$ | Ctrl, 5;<br>Exp, 5 |
| fig. S8C | AUC(norm<br>lick to CS);<br>acquisition;<br>CS-A vs.<br>CS-B | Paired t<br>test | G1, 0.965 (21);<br>G2, 0.638 (18);<br>G3, 1.098 (8) | G1, 2.98 [-3.44,<br>9.39];<br>G2, -2.71 [-<br>11.62, 6.21];<br>G3, 4.61 [-5.07,<br>14.30] | G1, 0.206;<br>G2, -0.146;<br>G3, 0.366 | Two-tailed p value<br>= G1, 0.345;<br>G2, 0.531;<br>G3, 0.304 | G1, 22;<br>G2, 19;<br>G3, 9 |
| fig.<br>S8G | AUC(norm<br>lick to CS);<br>reacquisiti<br>on; CS-A<br>vs. CS-B | Paired t<br>test | -1.469 (7) | -12.24 [-31.93,<br>7.46] | -0.519 | Two-tailed p value<br>= 0.185 | 8 |
| fig. S8H | AUC(norm<br>lick to CS);<br>acquisition;<br>CS-A vs.<br>CS-B | Paired t<br>test | 0.465 (7) | 3.88 [-15.84,<br>23.59] | 0.164 | Two-tailed p value<br>= 0.656 | 8 |
| fig. S8I | Delta<br>AUC(norm<br>lick); G2<br>vs. G4 | Two<br>sample<br>t test | 2.614 (20.51) | 31.11 [6.33,<br>55.89] | 0.920 | Two-tailed p value<br>= 0.016 | G2, 19;<br>G4, 8 |
| fig. S9C | Delta<br>AUC(norm<br>lick to CS-<br>A); each<br>group vs. 0 | One<br>sample<br>t test | G1, 2.897 (6);<br>G2, 2.056 (7) | G1, 38.59 [6.00,<br>71.18];<br>G2, 30.80 [-<br>4.63, 66.22] | G1, 1.095;<br>G2, 0.727 | Two-tailed p value<br>=<br>G1, 0.027;<br>G2, 0.079 | G1, 7;<br>G2, 8 |
| fig. S9F | Delta<br>AUC(norm<br>lick to CS-<br>A); each<br>group vs. 0 | One<br>sample<br>t test | G1, 3.959 (5);<br>G2, 0.537 (7) | G1, 7.09 [2.49,<br>11.70];<br>G2, 1.57 [-5.35,<br>8.50] | G1, 1.616;<br>G2, 0.190 | Two-tailed p value<br>=<br>G1, 0.011;<br>G2, 0.608 | G1, 6;<br>G2, 8 |
| fig. S9I | AUC(norm<br>lick to CS);<br>reacquisiti<br>on | Two<br>sample<br>t test | 3.420 (8.36) | 8.08 [2.67,<br>13.49] | 1.977 | Two-tailed p value<br>= $8.5 \times 10^{-3}$ | G1, 5;<br>G2, 6 |
| fig. S9K | AUC(norm<br>lick to CS);<br>acquisition | Two<br>sample<br>t test | 0.553 (8.68) | 2.70 [-8.40,<br>13.79] | 0.322 | Two-tailed p value<br>= 0.847 | G1, 5;<br>G2, 6 |
| fig.<br>S10B | Norm<br>response<br>to CS | Paired t<br>test | acquisition, -0.840<br>(9);<br>deg/ext, 0.104 (9);<br>test, 2.839 (9) | acquisition, -<br>1.58 [-5.81,<br>2.66];<br>deg/ext, 0.08 [-<br>1.56, 1.71];<br>test, 2.40 [0.49,<br>4.31] | acquisition, -<br>0.266;<br>deg/ext,<br>0.033;<br>test, 0.898 | Two-tailed p value<br>=<br>acquisition, 0.422;<br>deg/ext, 0.920;<br>test, 0.019 | 10 |
| fig.<br>S10E | Norm lick<br>to CS | Paired t<br>test | -0.916 (7) | -0.02 [-0.07,<br>0.03] | -0.324 | Two-tailed p value<br>= 0.390 | 8 |
| fig.<br>S11D<br>left | AUC(P<sub>freeze</sub><br>to CS+);<br>G1 vs. G2 | Two<br>sample<br>t test | 3.063 (13.50) | 8.86 [2.63,<br>15.08] | 1.487 | Two-tailed p value<br>= $8.7 \times 10^{-3}$ | G1, 8;<br>G2, 10 |
| fig.<br>S11D<br>right | AUC(P<sub>freeze</sub><br>to CS+);<br>G1 vs. G2 | Two<br>sample<br>t test | 2.708 (10.39) | 2.85 [0.52, 5.18] | 1.368 | Two-tailed p value<br>= 0.021 | G1, 8;<br>G2, 10 |
| fig.<br>S11F | AUC(selec<br>tivity); acq.<br>vs. reacq. | Paired t<br>test | G1, -3.702 (7);<br>G2, -1.378 (9) | G1, -3.13 [-5.12,<br>-1.13];<br>G2, -1.05 [-2.78,<br>0.68] | G1, -1.309;<br>G2, -0.436 | Two-tailed p value<br>=<br>G1, 0.008;<br>G2, 0.201 | G1, 8;<br>G2, 10 |
| fig. S11H | Selectivity; 0-10s vs. 19-29s | Paired t test | 2.629 (17) | 0.33 [0.06, 0.59] | 0.620 | Two-tailed p value = 0.018 | 18 |
| fig. S11I | Cocaine use; pre vs. post | Paired t test | 4.001 (11) | 11.08 [4.99, 17.18] | 1.155 | Two-tailed p value = 0.002 | 12 |
| fig. S13F | AUC(norm lick to CS); Ctrl vs. Exp | Two sample t test | -1.114 (13.84) | -54.93 [-160.82, 50.97] | -0.528 | Two-tailed p value = 0.284 | Ctrl, 8; Exp, 9 |
| fig. S13I, left | Delta lick to laser; pre-reacquisition | Two sample t test | 0.312 (3.76) | 0.23 [-1.84, 2.29] | 0.248 | Two-tailed p value = 0.772 | Ctrl, 3; Exp, 4 |
| fig. S13I, right | Delta lick to laser; post-reacquisition | Two sample t test | -3.095 (8.86) | -4.44 [-7.69, -1.19] | -1.421 | Two-tailed p value = 0.0131 | Ctrl, 8; Exp, 9 |
| fig. S13K, left | omission latency; w/ laser | Two sample t-test | -5.97 (13.89) | -0.91 [-1.24, -0.59] | -2.928 | Two-tailed p value = $3.6 \times 10^{-5}$ | Ctrl, 8; Exp, 9 |
| fig. S13K, right | omission latency; w/o laser | Two sample t-test | -0.187 (14.77) | -0.03 [-0.43, 0.36] | -0.091 | Two-tailed p value = 0.854 | Ctrl, 8; Exp, 9 |
| fig. S13M | Norm lick to reward; Ctrl vs. Exp | Two sample t test | 0.000 (13.02) | 0.00 [-0.11, 0.11] | 0.000 | Two-tailed p value = 1.000 | Ctrl, 8; Exp, 9 |
| fig. S13N | AUC(lick to CS); Ctrl vs. Exp | Two sample t test | 3.199 (10.83) | 430.40 [133.69, 727.12] | 1.737 | Two-tailed p value = 0.009 | Ctrl, 7; Exp, 6 |

## Methods

### Subjects

#### Mice

A total 182 adult wild type mice (C57BL/6; #000664; Jackson Laboratory; 78 males and 104 females) were used across cue-sucrose (Group1: n = 22 and Group2: n = 19, **Fig. 1**; Group2 Control for optogenetic dopamine inhibition: n = 10, **Fig. 2**; Group3: n = 9, **Fig. 3**; Group4: n = 8, **fig. S8**; Group1: n = 7; Group2: n = 8 in which CS-B acquisition was used to induce retrospective memory update, **fig. S9A-C**; Group1: n = 6; Group2: n = 8 with single CS+; **fig. S9D-F**; Group1: n = 5 and Group2: n = 6 in freely moving preparation, **Fig. S9G-M**), cue-alcohol (Group1: n = 7 males; Group2: n = 7 males; **fig. 4**), and cue-shock (Group1: n = 8 males; Group2: n = 10 males; **fig. S11A-H**), and response elimination (n = 8; **fig. S10C-E**)), standard extinction with optogenetic OFC inhibition (Ctrl: n = 15; Exp: n = 15; **Fig. 5A-D**), and random reward experiments (n = 4; **Fig. 5E-H**). In addition, 9 DAT-Cre heterozygous mice (B6.SJL-Slc6a3tm1.1(cre)Bkmn/J; #006660; Jackson Laboratory; 3 males, 6 females) were used as Group2 Experimental cohort for optogenetic dopamine inhibition **(Fig. 2)**. Animals that did not learn during acquisition phase were excluded from subsequent experiments (Group1/2: n = 13; Group3/4: n = 2; cue-alcohol: n = 3; optogenetic OFC inhibition: n = 5; optogenetic dopamine inhibition: n = 3; Group1/2 in **fig. S9A-C**: n = 1; Group1/2 in freely moving preparation: n = 7; see Behavioral analysis for cue-sucrose task section in Analysis to find detailed criteria for non-learners). Note that a comparable fraction of non-learners (∼20%) has been reported previously under similar 50% partial reinforcement condition (Fig 6B in Burke et al.(*45*)). Additionally, animals that failed to extinguish behavior during extinction phase were excluded (Group2: n = 1; Group3: n = 1; optogenetic OFC inhibition: n = 1; Group1 in **fig. S9D-F**: n = 2). Animals implanted with optic fibers for fiber photometry or optogenetic manipulation were single-housed following surgery, whereas behavior-only animals were group housed. All mice were maintained on a 12 h light/dark cycle. For all experiments except for cue-alcohol and cue-shock conditioning, daily water intake was adjusted to maintain animals at ∼85-88% baseline weight. Body weight was recorded daily and maintained above 82% pre-deprivation weight. All experimental procedures were performed in accordance with the National Institutes of Health Guide for the Care and Use of Laboratory Animals and approved by the UCSF Institutional Animal Care and Use Committee.

#### Rats

Ten Long-Evans rats (5 males, 5 females) were used. Rats were food deprived to 85% of their ad libitum weight throughout the experiment except for the first week between Extinction/Degradation and Recovery tests (See Behavioral tasks). Food deprivation resumed beginning one week prior to Recovery tests. All experimental procedures were performed in strict accordance with protocols approved by the Animal Care and Use Committee at Johns Hopkins University.

#### Humans

Twenty-eight non-dependent, non-treatment seeking stable cocaine users (**Supplementary Table 1**) were recruited from the community through advertisements in local newspapers. For all individuals, the primary and preferred route of cocaine self-administration was intranasal. Exclusion criteria included current or past Diagnostic and Statistical Manual of Mental Disorder, Fourth Edition (DSM-IV) (*105*) dependence (a moderate to severe substance use disorder) on any substance, a current non-substance related axis I disorder, and any cardiovascular, neurological or other medical condition that might be aggravated by study participation. Four participants met DSM-IV criteria for current (n = 2) or past (n = 2) cocaine abuse (a mild substance use disorder), whereas 6 met criteria for past or current substance abuse of another substance (alcohol: n = 4; cannabis: n = 3; ketamine: n = 1; heroin: n = 1). Participants were physically healthy as determined by a medical exam, electrocardiogram and standard laboratory tests. All participants gave written informed consent, and all procedures were carried out in accordance with the Declaration of Helsinki and were approved by the Research and Ethics Board of the Montreal Neurological Institute.

#### Surgeries

All surgeries were performed under aseptic conditions following procedures similar to those described previously (*42*). Mice were anesthetized with isoflurane (3-5% for induction and 1-2% for maintenance). A custom-designed head ring was implanted on the skull for head fixation in all mice except for animals used for cue-shock experiments. Mice used for cue-shock experiments or rats did not undergo surgeries. For optogenetic inhibition of VTA dopaminergic neurons (**Fig. 2**), 500 nL of Cre-dependent stGtACR2 (AAV1-hSyn1-SIO-stGtACR2-FusionRed, 6.3 x 10^12^ GC/ml) was injected bilaterally in VTA (AP -3.2, ML ±0.99, DV -4.51 with 5° angle). To prevent backflow, the injection needle was left in place for additional 10 min after injection. Optic fibers (NA 0.39, 200 μm, RWD) were implanted 100-200 μm above the virus injection sites. In a subset of animals (cue-sucrose experiment, Group 1: n = 10; Group 2: n = 8; Group 2 optogenetic dopamine inhibition, control (WT): n = 5; experimental (DAT-Cre): n = 5), 500 nL of dLight1.3b (AAVDJ-CAG-dLight1.3b, 2.4 x 10^12^ GC/ml) was injected into the nucleus accumbens core (AP 1.3, ML +/-1.4, DV -4.55) at 100 nL/min to allow simultaneous measurement of dopamine release. An optic fiber (NA 0.66, 400μm, Doric Lenses) was implanted 350 μm above the virus injection site. Animals used in the cue-sucrose, response elimination, and random reward experiments were water-deprived starting one week before behavioral training and remained water-restricted throughout the experiments. Animals recovered from surgery for at least one week prior to the onset of water deprivation. After completion of experiments, viral expression and optic fiber placements were confirmed using a Keyence microscope.

### Behavioral tasks

#### Cue-sucrose task (Fig. 1-4, fig. S2-10)

To compare two distinct types of extinctions (standard extinction and retrospective extinction) predicted from ANCCR in a way that dissociates predictions of the latent cause model, we designed a Pavlovian conditioning paradigm using two auditory cues. Mice were first trained to associate two 0.25 s tones (12 kHz and 3 kHz) with a delivery of a sucrose drop (∼5 ul) during the *Acquisition* phase. We refer to these reward-associated cues as CS-A and CS-B throughout the study; the cue identities of CS-A and CS-B were counterbalanced across animals. Sucrose was delivered 1s after cue offset with 50% probability. After a 3 s fixed delay following the outcome (reward or omission), the inter-trial interval (ITI) commenced. The ITI was randomly drawn from a truncated exponential distribution (mean = 60 s; maximum = 180 s). Each session consisted of 60 trials (30 CS-A and 30 CS-B). Once animals showed significantly greater anticipatory licking during the 1.25 s cue-reward delay compared to the 1.25 s pre-cue baseline (“lick to CS”), they received at least 3 further days of training to ensure stable asymptotic behavior prior to transitioning to the *CS-A extinction* phase.

Animals were then assigned to one of four groups that underwent distinct *CS-A extinction* procedures: <u>1) Group1 (Standard extinction of CS-A; CS-A omission only daily sessions).</u> Thirty CS-A trials were presented without subsequent reward, and no CS-B trials were delivered. To match total session duration with *Acquisition*, ITI was extended to 120 s (twice the *Acquisition* ITI). This allows animals to extinguish behavior to CS-A through degradation in estimated reward rate, while leaving CS-A←reward retrospective association intact. <u>2) Group2 (Retrospective extinction of CS-A; CS-A omission trials</u> <u>intermixed with CS-B trials over many daily sessions).</u> Thirty CS-A omission trials were intermixed with 30 CS-B trials that retained their 50% reward probability. Because rewards delivered on CS-B trials drive degradation of the CS-A←reward retrospective association, this paradigm enables retrospective extinction. ITI duration was kept identical to *Acquisition* (60 s). For a direct comparison between Group1 and Group2, we ran the first two cohorts (Group1: n = 12; Group2: n = 11) in which the number of *CS-A extinction* sessions was matched (14-16 days for the cohort 1, 18-20 days for the cohort 2). The trial at which the fitted sigmoid reached 25% of the (max-min) range was defined as the moment of extinction, and the session containing this trial was designated as the day of behavioral extinction. Extinction proceeded substantially more slowly in Group2 (6.64±2.9 days, mean±STD) than in Group1 (1.42±0.49 days) (**fig. S3A-D**). As a result, Group1 experienced a larger number of *CS-A extinction* sessions after their behavior had already reached asymptotic extinction than Group2. This difference in behavioral history could confound later memory test analyses by making animals with longer behavioral extinction produce slower reacquisition (although we did not find any evidence supporting that Group1 with longer behavioral extinction than Group2 produced faster reacquisition in our post-hoc analysis; **fig. S2**). To eliminate this confound, we ran the third cohort (Group1: n = 10; Group2: n = 8) in which extinction was matched by behavior rather than by number of sessions. Using the first two cohorts, we quantified the number of days from the moment of CS-A behavior extinction to the end of the *CS-A extinction* phase (15.67±2.36 days for Group1; 10.09±3.15 days for Group2). To identify the moment of behavioral extinction, we concatenated lick to CS values across extinction sessions and fit the trial-by-trial data with a sigmoid function. For the second cohort, we again fixed the number of *CS-A extinction* sessions for Group 1 (14 days). Group2, however, was trained until each animal experienced the same number of post-behavior extinction *CS-A extinction* sessions as the average number observed in Group2 (∼10 days) of the first cohort. To achieve this in real time while conducting the experiments, for each day, we tested whether lick to CS for both cues remained significantly above zero. Once licking to neither cue was significant, training continued for an additional 10 sessions, corresponding to the average number of post-extinction sessions in Group1 from the first cohort. When we later analyzed the second cohort using the same sigmoid-based curve-fitting method applied to the first cohort, we confirmed that there was no significant difference in number of additional days of *CS-A extinction* following their respective extinction days (Group1: 12.2±0.87 days; Group2: 14.63±5.19 days; two-sampled t-test, p = 0.190). In the first cohort animals, we tested whether increasing the reward amount on CS-B trials would accelerate extinction in Group2, given the hypothesis that retrospective updating depends on the magnitude of dopamine release at reward. However, extinction speed did not differ between animals that received larger reward (7.17±2.85 days to behavior extinction) and those that did not (5.92±2.81 days; two-sampled t-test, p = 0.695), thus these data were combined. Subsequent dopamine release measurement from the NAc showed that CS-B rewards elicited large dopamine responses (75.06±23.36% of the unpredicted reward response, mean±STD), likely reflecting partial reinforcement effect, suggesting that the standard sucrose reward already produced sufficient dopamine release for retrospective updating. Finally, comparisons with cue-alcohol experiments suggest that the slower extinction observed in cue-sucrose Group2 may reflect cross-cue generalization (see **Supplementary Note 3**). For optogenetic inhibition of dopamine neurons (**Fig. 2**), a 473 nm laser (8-11mW, Shanghai Laser & Optics Century Co.) pulse was delivered for 5 s following reward delivery. All optogenetic manipulations were initiated more than three weeks after surgery to ensure sufficient viral expression. <u>3) Group3 (Retrospective extinction of CS-A; CS-A omission only</u> <u>daily sessions followed by CS-B trials only daily sessions).</u> To dissociate predictions of ANCCR from those of the latent cause model, we separated CS-A omission trials and CS-B trials into different daily sessions. Animals were first trained on CS-A omission only sessions (30 trials per session; identical to Group 1 *CS-A extinction*) for 5 days (“early sessions”), followed by CS-B only session with 50% reward probability for 13 additional days (30 trials per session; “late sessions”). For one animal that did not extinguish lick to CS-A within the initial 5 early sessions, the early session phase was extended to 11 days to ensure at least two complete days of behavior extinction before transitioning to the CS-B only late sessions. In both early and late sessions, the ITI was maintained at 120 s (twice the *Acquisition* ITI) to match overall session duration to *Acquisition*. <u>4) Group 4 (Retrospective extinction of both CS-A</u> <u>and CS-B; CS-A omission only sessions followed by reward only sessions)</u>. To further test retrospective extinction under conditions that degrade memories in a non-cue selective manner, the CS-B trials only late sessions in Group3 were replaced with reward only sessions (30 rewards per session). Average inter-reward interval in reward only sessions was 120 s. Under the assumption that animals treat these sessions without CS-A or CS-B as belonging to the same physical context as acquisition, these reward only sessions degrade all retrospective associations linked to the reward, thereby producing retrospective extinction for both CS-A and CS-B.

After the *CS-A extinction* phase, animals in all groups underwent the same *CS-B extinction* phase for 8-10 days (but extended for 5 animals who were resistant to extinction), where 30 CS-A and 30 CS-B trials were intermixed and no rewards were delivered with 60 s mean ITI. This procedure does not modify the CS-B←reward retrospective association, while suppressing behavior through reduction in estimated reward rate; thus, CS-B←reward memory remains intact in Group1-3, whereas it remains degraded in Group4 due to non-selective retrospective extinction applied during *CS-A extinction* phase. After CS-B extinction phase, CS←reward memory was assessed in all groups during the *Reacquisition* phase, which was procedurally identical to the *Acquisition* phase. *Reacquisition* lasted for 7-9 days.

To further test whether physical context is inferred primarily from the sensory environment, including the set of neutral cues present during training, while preserving the standard versus retrospective extinction manipulation, we trained two additional groups (Group1 and Group2) in which CS-A and CS-B trials were completely separated into different sessions (**fig. S9A-C**). In this cohort, animals were first trained to associate CS-A with sucrose reward. Once animals acquired stable conditioned responding to CS-A after 8 days of training, they were divided into two groups. During the subsequent CS-B acquisition phase, Group1 animals experienced presentations of a novel auditory cue (CS-B) without reward, whereas Group2 animals experienced pairings of CS-B with sucrose reward. After >6 days of CS-B training, both groups underwent CS-A extinction, during which CS-A was presented without reward. Following 6 days of extinction, animals underwent 6 days of CS-A reacquisition training. This design temporally separated experiences involving CS-A and CS-B, such that the two cues were never encountered within the same behavioral sessions.

To test whether retrospective extinction remains effective at inducing unlearning in a simpler design with presentation of uncued rewards, instead of rewards following another reward-predictive cue, we trained two additional groups (Group1 and Group2) using a single CS+ paradigm (**fig. S9D-F**). During acquisition, animals learned to associate one auditory cue (CS+) with sucrose delivery, while a second cue (CS−) was never rewarded. Cue identities (3 kHz and 12 kHz) were counterbalanced across animals. Each cue was presented for 0.25 s, followed by a 1 s trace period; on CS+ trials, a drop of sucrose was delivered after the trace period. The ITI was drawn from a uniform distribution between 240 and 360 s (note that uniform distribution was used instead of exponential distribution to avoid extremely long ITI), and each session consisted of 6 CS+ and 6 CS− trials. Animals were moved to extinction after showing significant licking to the CS+ and no significant response to the CS− for >5 days. During extinction, Group1 received 6 CS+ and 6 CS− trials per session, with sucrose omitted on all CS+ trials. Group2 received the same cue presentations, but also received uncued sucrose rewards randomly during the ITI. The inter-reward interval for these uncued rewards was drawn from an exponential distribution with a mean of 625 s, matching the average reward interval during acquisition, calculated as 2 × (1.25 s cue–reward delay + 300 s ITI). After 16 days of extinction, animals were returned to reacquisition training for 7 days. If animals had not accumulated at least 8 days of non-significant licking to CS+ by the end of the initial 16 day extinction period, extinction training was extended until they reached this criterion. Animals that failed to meet the extinction criterion within 20 days were classified as non-extinguishers and excluded from subsequent investigation. A similar paradigm was tested in a freely moving preparation (**fig. S9G-M**) to determine whether retrospective memory erasure extends beyond the head-fixed preparation. Freely moving chambers were modified from standard home cages. A lick spout was mounted on one wall, and the floor was covered with copper tape connected to a lickometer to detect licking. Speakers were positioned outside the chamber, and the lid contained multiple slits to allow auditory cues to enter. The entire chamber was placed inside a sound-attenuating dark enclosure. Each cue was presented for 10 s to provide sufficient time for animals to approach the reward port and exhibit anticipatory licking. An ∼10 ul drop of sucrose reward was delivered at cue offset. A larger volume than in the head-fixed preparation was used to maintain motivation because reward collection requires substantially greater effort in freely moving animals. The ITI was randomly drawn from a truncated exponential distribution (mean = 120 s; maximum = 360 s). Note that a similar paradigm was used for cue-alcohol (**Fig. 4**) and cue-shock (**fig. S11**) experiments.

#### Reward identity specific contingency degradation task (**fig. S10A,B**)

We wanted to test whether retrospective contingency degradation is reward identity specific. To this end, we wanted to conduct experiments with highly sensorily separate rewards (e.g., sweet liquid vs non-sweet solid grain pellets). Because this was challenging in a head-fixed preparation, we performed these experiments in freely moving rats. During the *Acquisition* phase, two auditory cues were paired with different food rewards over eight daily sessions. The two food rewards (45 mg grain pellets and 0.1 ml of 20% sucrose) were preceded by a pure tone or white noise. Each auditory cue lasted for 20 seconds and was followed by a 50% probability of its associated reward. The ITIs were exponentially distributed with a mean of 4 minutes and a range of 21 s and 14 min. Each session lasted 70 minutes, and within a session, each trial type was presented eight times in pseudorandom order. Rats were guaranteed to experience four rewarded trials and four nonrewarded trials of each type. Cue-reward assignments were counterbalanced with sex. During the *Extinction/Degradation* phase, one cue underwent extinction while the other underwent contingency degradation over eight daily sessions. The extinguished cue terminated with a reward probability of 0, while the degraded cue offset was still associated with 0.5 reward probability. During the inter-trial intervals, the reward associated with the degraded cue was delivered every 20 seconds with a 50% probability except when the upcoming trial was less than 20 seconds away. This was to avoid noncontingent rewards being delivered during the 20 second pre-cue baseline. Cue-reward assignments, noncontingent reward identity, and sex were counterbalanced. Following the final session of *Extinction/Degradation*, rats stayed in their home cage for two weeks and were then placed in two spontaneous recovery tests. During each test, the auditory cues were presented with the same frequency and pseudorandom order as the prior phases, but no rewards were delivered.

#### Response elimination task (**fig. S10C-E**)

Classic autoshaping studies in pigeons have shown that presenting uncued rewards while omitting rewards on CS+ trials (‘negative contingency’ or contingency degradation in general) eliminates the conditioned response but does not erase the underlying cue-reward memory (*52*, *53*, *106*). Evidence comes from the observation that, after response elimination, subsequent standard extinction sessions often produce a recovery of conditioned responding, indicating that the original memory remains intact. This behavioral pattern directly contradicts a key prediction of ANCCR, which proposes that uncued outcomes should trigger retrospective updating that weakens the original association and produces true memory erasure. To test whether uncued rewards can erase memory in rodents under a head-fixed paradigm, we adapted the classic pigeon ‘negative contingency’ design (*52*). We used partial reinforcement conditions, because prior study (*106*) reported stronger recovery following response elimination when acquisition involves partial reinforcement. During *Acquisition*, mice learned that 0.25 s auditory cue (12 kHz tone) predicted a 5 ul sucrose reward with 50% probability, delivered after 1 s trace interval. The ITI was uniformly distributed between 240 s to 360 s, and each session contained 12 trials. Once lick to CS was significantly greater than zero for 5 days, animals advanced to *Response elimination* phase. In this phase, all cue-paired rewards were omitted, and instead six uncued rewards were delivered at random ITI times. This design directly mimics the ‘negative contingency’ paradigm used in the pigeon autoshaping literature (*52*). Response elimination continued until mice showed no significant CS-evoked licking for more than 5 consecutive days, or until a total of seven days without significant CS licking was reached. After response elimination, mice underwent three standard extinction sessions, where only the cue was presented without any associated or unassociated rewards. These sessions were used to determine whether behavioral recovery occurred.

#### Cue-alcohol task (**Fig. 4**)

To familiarize mice with alcohol, all animals underwent an intermittent access two bottle choice procedure prior to cue-alcohol conditioning task. Animals received three 24 h drinking sessions per week with continuous access to a 20% (v/v) ethanol bottle and a water bottle, separated by 24 h and 48 h withdrawal periods during weekdays and weekends, respectively. During withdrawal periods, animals had access to water only. The positions of alcohol and water bottles were alternated at the start of each drinking session to control for side preferences. A ‘spill cage’ without an animal was used to measure evaporation or spillage, and consumption amounts were corrected accordingly. Alcohol and water intake were measured for each 24 h drinking session. After 14 drinking sessions, animals consumed an average of 10.88±2.51 g/kg (mean±STD) of alcohol and displayed an average alcohol preference (over water) of 0.50±0.11, with no difference across groups (consumption, G1: 10.65±2.43, G2: 11.11±2.75, two-sampled t-test, p = 0.746; preference, G1: 0.51±0.10, G2: 0.49±0.13, two-sampled t-test, p = 0.744). Because this study was not intended to model severe alcohol dependence but rather to use alcohol as an appetitive outcome for cue-outcome conditioning, we included all animals regardless of their final alcohol intake or preference. Importantly, all mice exhibited robust consummatory licking for alcohol during the subsequent head-fixed cue-alcohol conditioning task.

To test standard vs. retrospective extinction of cue-alcohol memory, we used Pavlovian conditioning paradigm similarly to the cue-sucrose task. During the *Acquisition* phase, animals were trained to associate one auditory cue (CS+) with alcohol delivery and a second cue (CS-) with no outcome. Cue identities (3 kHz and 12 kHz) were counterbalanced across animals. Each cue was presented for 0.25 s, followed by a 1 s trace period, after which a drop of alcohol (∼9 μl) was delivered on CS+ trials. Following the outcome, the ITI commenced. The ITI was drawn from a uniform distribution between 240 s and 360 s. Each session consisted of 6 CS+ trials and 6 CS-trials. Behavior was quantified using a selectivity index, defined as 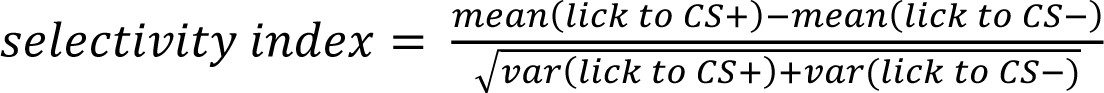, which measures the separation between CS+ and CS-licking responses. Animals were trained until they exhibited >6 days with a selectivity index greater than 1 before transitioning to the *Extinction* phase. Average selectivity index was not different across groups for the last 6 days of *Acquisition* (Group1: 1.77±0.57; Group2: 1.12±0.75; two-sample t-test, p = 0.121). For *Extinction*, animals were divided into two groups. <u>1)</u> <u>Group1 (Standard extinction).</u> All rewards in CS+ trials were omitted. Animals therefore experienced no alcohol during extinction in this group. This procedure suppresses behavior by reducing the estimated reward rate but leaves the original CS+←alcohol memory intact. <u>2) Group2 (retrospective</u> <u>extinction)</u>. Rewards in CS+ trials were omitted as in Group1, but animals additionally received 6 uncued alcohol rewards delivered at random times during ITIs (half following CS+ trials and half following CS-trials). To maximize separation between uncued rewards and cue presentations while preserving a sufficiently large window for random timing, the first and last 10 s of each ITI were excluded from the possible reward delivery windows. These uncued rewards drive retrospective degradation of CS+←alcohol memory. Extinction training continued until each animal exhibited >6 days with a selectivity index smaller than 0.2. Average selectivity index was not different across groups for the last 6 days of *Extinction* (Group1: 0.07±0.11; Group2: -0.04±0.24; two-sample t-test, p = 0.327). After extinction, memory was tested during *Reacquisition* phase, which was trained for 17 days and was procedurally identical to the *Acquisition* phase.

#### Cue-cocaine task (**fig. S11I**)

Human participants were recruited following a three-stage screening procedure. First, a semi-structured telephone screening was conducted. Second, those who were interested and tentatively met the entry criteria were invited for a face-to-face psychiatric interview, using the Structural Clinical interview for DSM-IV (SCID) (*105*). Third, volunteers underwent a medical examination, conducted by a physician, to ensure that it was medically safe for volunteers to participate in the study. Participants took part in one of three different studies involving intranasal cocaine self-administration (*107–110*). For each study, in-depth information was obtained from participants about their cocaine use histories using a timeline follow-back (TLFB) procedure (*111*), including number of occasions used per month during the past year, overall lifetime use, estimated amount (milligrams) used per occasion and routes of administration (see **Supplementary Table 1**).

Detailed characteristics of the cocaine self-administration procedures are described in previous reports (*107–110*). In each study participants self-administered between 1 mg/kg and 5.1 mg/kg cocaine, taken intranasally, on two to four separate occasions, in a hospital setting. Throughout testing, a project-dedicated nurse and emergency physician were on-site. Following testing, participants were kept overnight for observation with a nurse onsite and an emergency physician on call. There were no serious adverse effects. Since these two participants indicated no further interest in study participation, their information was excluded from this report.

For follow-up recruitment, participants were re-contacted via telephone or email. They were informed that additional information was required for a follow-up project of the initial study they participated in. They were not told that these self-reports would be compared to information obtained prior to the initial study. The sample consisted of 12 respondents (**Supplementary Table 1**). Sixteen participants could not be re-contacted, having either changed their telephone number or email address without notifying us, or did not respond to multiple re-contact attempts. Thus, the present findings need to be interpreted in light of the following considerations. First, the sample size of 12 (out of 28) respondents is modest. Although no differences were observed between respondents and non-respondents a sample bias cannot be excluded. The time interval between participation in the initial study and follow-up evaluation was on average 21.5±9.4 months (mean±STD). Two of the 12 respondents met criteria for at least 1 past substance use disorder (Subject 1: cocaine and cannabis abuse and Subject 2: alcohol and ketamine abuse) prior to study participation. Respondents were questioned about their cocaine use frequency and route of administration during the 12-month period immediately after study participation using a TLFB procedure (*111*) (**Supplementary Table 1**). In addition, the incidence of potential cocaine use disorders was assessed, using the same diagnostics (DSM-IV) as those used during the SCID interview prior to participation of the initial study. Eight participants were re-interviewed on the telephone, four in person. Since there were no differences in cocaine use between subjects from each interview medium, their data were combined.

#### <u>Cue-shock task</u> (**fig. S11A-H**)

Mice were handled for 2-5 minutes per day for the 5 days preceding cue-shock conditioning. Animals were placed into a sound-isolating chamber and received auditory cues and/or electric footshock (outcome) controlled by MED-PC (MED Associates). During *Acquisition*, animals were trained to associate one auditory cue (CS+) with a 0.35 mA footshock and a second auditory cue (CS-) with no outcome. Cues were 30 s tones (5 and 10 kHz), with CS+ and CS-identities counterbalanced across animals; on CS+ trials, the cue co-terminated with a 1-s shock. Each session consisted of 5 CS+ and 5 CS-trials. Trials were separated by an ITI drawn from a uniform distribution between 108 s and 134 s (mean of 120 s with a 10% jitter). Learning was assessed using the fraction of immobility during first 10 s of CS period. The fraction of immobility during the 10 s pre-cue baseline was also quantified. Animals were trained for 10 days, during which the population-averaged fraction of immobility during CS+ was significantly larger than that during the CS- and baseline periods for last 7 days. To quantify learning at the individual-animal level, we used the same selectivity index applied in the cue-alcohol task, substituting the average fraction of immobility during the CS for the lick to CS measure. Animals were then divided into two groups such that the average selectivity index on the last 3 days of *Acquisition* was not different across groups (Group1: 1.36±0.50; Group2: 1.39±0.45; two-sample t-test, p = 0.897) before transitioning to *Extinction*. During the *Extinction* phase, two groups went through different procedures. <u>1) Group1 (Standard extinction)</u>. All shocks in CS+ trials were omitted. <u>2) Group 2 (Retrospective extinction)</u>. Shocks in CS+ trials were omitted as in Group1, and animals additionally received 4 uncued shocks at random times during ITIs (2 following CS+ trials and 2 following CS-trials). The first and last 12 s of each ITI (10% of mean ITI) were excluded from the possible shock delivery windows. During the first two extinctions sessions, 10 uncued shocks (5 following CS+ trials and 5 following CS-trials) were mistakenly delivered. Because animals completed 15 total *Extinction* sessions (i.e., 13 correctly configured sessions following the first two), it is unlikely that the effects of these initial sessions persisted into the later stages of *Extinction*. After 15 days of *Extinction* with the comparable immobility fraction across cues for the last 2 days (Group1: 0.20±0.68; Group2: 0.22±0.46; two-sample t-test, p = 0.96), memory was tested during 14 days of *Reacquisition*.

Note that we used ABB design, in which *Acquisition* occurred in context A, and *Extinction* and *Reacquisition* occurred in context B. This design was adopted to dissociate the contributions of retrospective memory and reward-history-based priors to the speed of reacquisition. In ANCCR, animals estimate event rate (e.g., reward/shock rate) in a context-specific manner, and these estimates are restored quickly at the beginning of each session (or during reacquisition after long absence of outcomes) once any outcome is experienced in that context possibly using memory of probability of outcomes given an outcome (e.g., P(reward|reward) or P(shock|shock)). Thus, rapid reacquisition in Group1 during cue-sucrose/alcohol experiments could, in principle, be attributed to a high prior of P(reward|reward) as it does not update during extinction due to absence of rewards and remains as high as in *Acquisition*. By switching to a novel context B for extinction and reacquisition, Group1 enters *Reacquisition* with P(shock|shock) = 0 because no shocks were ever experienced in context B, eliminating any inflated prior of estimated shock rate. In contrast, Group2 receives uncued shocks in context B, resulting a nonzero P(shock|shock) at the start of *Reacquisition*. Therefore, if reacquisition were driven by shock-history priors (higher in Group2 than Group1) rather than retrospective memory (erased in Group2, while preserved in Group1), Group2 should reacquire faster than Group1. Using ABB design thus isolates retrospective-memory effects from confounding priors based on outcome statistics. Context 1 consisted of entirely grey siding, a rectangle arena shape, purple light, and vanilla scent, while context 2 consisted of vertical, high contrast stripes, a hexagonal arena, green light, and lemon scent. Both contexts have a shock grid floor. Context identities were counterbalanced across animals.

#### Simple extinction task with OFC optogenetic inhibition (**Fig. 5A-D, fig. S13**)

During *Acquisition*, 0.25 s auditory cue predicted delivery of a 5 ul sucrose reward with 50% probability following a 1 s trace interval. The ITIs were drawn from a truncated exponential distribution (mean = 120 s, maximum = 360 s) and began after a 3 s fixed post outcome delay. Once animals showed significantly elevated cue-evoked licking for more than 5 days, they advanced to the *Extinction* phase, in which rewards were omitted. After >2 days in which lick to CS was no longer significant, animals proceeded to *Reacquisition*. OFC activity was optogenetically inhibited during one of two task epochs: 1) cue-reward delay period (1.25 s; 473 nm, 8-11mW, Shanghai Laser & Optics Century Co.), or 2) outcome period (5 s). Each animal experienced both inhibition conditions, achieved by running two sequential extinction-reacquisition cycles (e.g., *Acquisition* → *Extinction* → *Reacquisition with OFC inhibition #1* → *Extinction* → *Reacquisition with OFC inhibition #2*). The order of inhibition periods was counterbalanced across animals. During the outcome period inhibition condition, laser delivery was terminated on Day 4 of *Reacquisition* to test whether animals rapidly reacquired the conditioned response once OFC inhibition was removed. In a subset of animals (n = 3/15 for Ctrl, 4/15 for Exp), thirty 5 s laser pulses were delivered at random times during ITI to test whether laser alone could elicit licking. This test was performed twice: once after animals had learned the task but before *Extinction* (‘test laser before *reacq*.’ in **fig. S13G**), and once after *Reacquisition* (‘test laser after *reacq*.’). Following the ‘test laser before *reacq*.’ session, animals underwent one additional acquisition session without laser before proceeding to extinction. To test the effect of OFC outcome period inhibition on initial acquisition, a separate cohort underwent *Acquisition* with 5 s OFC inhibition during outcome period (**fig. S13N)**. These animals did not undergo *Extinction* or *Reacquisition*.

#### Random reward task with OFC imaging (**Fig. 5E-H, fig. S14**)

Low (75 s inter-reward interval) and high (15 s) reward rate blocks were alternated twice within each session. Each block consisted of 30 rewards (∼3 μl sucrose). The experiment was conducted over two days with reversed block order across days (Day 1: high→low→high→low; Day2: low→high→low→high). Day 2 was the primary data analyzed due to the animals already being habituated to reward delivery on the previous day.

### Photometry recording

After three weeks from the virus injection, fiber photometry recordings were performed using an open-source system (PyPhotometry). In PyPhotometry, two LED wavelengths were emitted alternatively at 130 Hz through a fluorescence mini cube (Doric Lenses) using time-division sequence. Fluorescence signals were collected by a separate photodetector (Doric Lenses). Light intensity at the tip of the patch cord was maintained at 40 µW (470 nm) and 20 µW (405 nm) across sessions. Raw fluorescence signals were bandpass filtered (0.01-20 Hz) to remove low-frequency drift caused by photobleaching or motion artifacts and to suppress high-frequency electrical noise. The 405 nm signal was then linearly regressed onto the 470 nm signal to obtain a fitted isosbestic reference. Fractional fluorescence signal (ΔF/F) was computed as (470 nm signal – fitted 405 nm signal)/(fitted 405 nm signal).

### Two-photon microscopy

Calcium activity of OFC neurons was imaged using two-photon microscopy after expressing the calcium indicator (GCaMP6s) under CamKIIα promoter. A minimum of 6 weeks was allowed for sufficient viral expression before imaging began. Imaging was performed using Bruker Ultima 2pplus two-photon microscope equipped with a Cousa 10x long working distance objective lens (*112*) and a 920 nm excitation laser (SpectraPhysics). A resonant scanner combined with an electrically tunable lens was used so simultaneously image three planes at ∼8Hz per plane. Imaging planes were spaced >50 μm apart to minimize overlap of cells across adjacent planes. Laser power at the sample was maintained at 25-35 mW throughout sessions. Animals were positioned on a tip, tilt, and rotation stage (Thorlabs) to precisely align the surface of lens perpendicular to the excitation path. Goniometer readings were recorded on each day, and imaging planes were compared to the reference plane acquired on the first imaging day to ensure that the mouse was consistently positioned at the same head angle across all imaging sessions.

### Analysis

#### Behavioral analysis for cue-sucrose task

For the cue-sucrose task, memory was assessed by quantifying the speed and strength of reacquisition using two complementary measures. First, we computed the area under curve (AUC) of lick to CS vs. trial plot (e.g., **Fig. 1I**; used throughout the paper). Lick to CS was defined as the difference in lick counts between post-cue and pre-cue 1.25 s windows. For freely moving preparation, Lick to CS was defined as the difference in lick counts during the last 5 s of cue presentation and pre-cue 5 s window. Lick to CS values were normalized to the average lick to CS measured during last 3 days of acquisition for each animal to account for variability in baseline lick rates across animals. AUC was calculated over the first 3 days of acquisition or reacquisition. Because acquisition and reacquisition exhibited markedly different temporal dynamics in the freely moving preparation due to additional learning of reward collection strategy during acquisition, AUC was calculated over the first 9 days of acquisition (**fig. S9K**) and the first day of reacquisition (**fig. S9I**). Importantly, AUC was compared only between groups within each phase and was not compared directly between acquisition and reacquisition in this case. The AUC was used as the primary measure because it allows inclusion of all animals, including those that did not show clear evidence of behavioral reacquisition. We used the same AUC metric to measure the speed of extinction (**fig. S3,4**). However, because AUC reflects both the speed and strength (i.e., asymptotic level of licking) of reacquisition, we additionally used cumulative sum-based analysis to dissociate these components (e.g., **Fig. 1J-L, fig. S2A,B**). For cumulative-sum analysis, we plotted the cumulative sum of lick to CS across trials and identified the learned trial as the first trial at which the distance between the cumulative-sum curve and the unity (diagonal) line exceeded 80% of the maximum distance. Because this threshold is relative, it can identify a learned trial even when the cumulative-sum trace remains close to zero. To prevent classifying such cases as learners, we computed the average lick to CS using trials after the detected learned trial (‘average lick to CS after learning; **fig. S2B**) and excluded any animals with <0.5 licks during 1.25 s (or <2 licks during 5 s window for freely moving preparation) on average as non-learners. We included 2 extinction sessions and 6 reacquisition sessions for this analysis. To directly compare acquisition and reacquisition learned trials directly, we prefixed the cumulative-sum vector with an arbitrary list of 30 zeros (30trials/session*2 sessions) corresponding to extinction trials, followed by 6 acquisition days when performing cumulative-sum analysis during acquisition.

Memory was additionally assessed by measuring spontaneous recovery during extinction (**Fig. 1M,N**). Because the number of extinction days experienced after behavior had extinguished differed across groups—and spontaneous recovery is known to weaken over the course of extinction—we selected extinction sessions in a way that matched behavioral history across groups. In Group2, extinction of CS-A occurred in the presence of a non-zero reward rate (due to ongoing rewards in CS-B trials), which could contribute spontaneous recovery and confound the interpretation of memory strength. Therefore, for Group2, we used the 8 days of CS-B extinction phase, during which no rewards were delivered, as the comparison window. We first computed, across Group2 animals, the average number of days with no significant lick to CS-A (13±4.7 days) during the CS-A extinction phase. For each Group1 animal, we then identified the extinction day on which it had experienced the same number of days without significant CS-A licking as the Group2 average. The eight subsequent extinction days in each animal were used for comparison with the corresponding CS-B extinction window in Group2. Because spontaneous recovery is mostly prominent at the beginning of each session, we quantified it by computing the AUC of lick to CS vs. trial plot using only the first 5 trials of each session.

#### Analysis of dopamine response for cue-sucrose task

Dopamine learning was similarly assessed as behavioral learning by substituting ‘lick to CS’ with ‘DA to CS’—defined as the difference in area under the dLight fluorescence trace between post-cue and pre-cue 1.25 s window (**Fig. 1Q**). DA to CS value was normalized to the maximum of the mean dopamine responses to rewards during the first 3 days in learning, measured as the difference in fluorescence AUC between the post-reward and post-cue 1.25 s windows. This normalization enables comparison across animals by accounting for differences in overall fluorescence that may arise from variability in viral expression or in the alignment between fiber tip and the expression site. For **fig. S5**, we showed the DA to CS normalized to asymptotic DA to CS for each phase (acquisition or reacquisition) as well. To quantify the effectiveness of optogenetic inhibition of dopaminergic neurons in **Fig. 2D**, DA to reward with optogenetic inhibition was compared to that without inhibition. For this specific analysis, we used 5 s analysis window as inhibition lasted for 5 s. Dopamine dip at reward omission (DA to omission) was similarly measured as difference in fluorescence AUC between post-omission (when reward was expected) and post-cue 3 s windows (**fig. S12E,N,O**).

#### Behavioral analysis for cue-alcohol and cue-shock task

Learning was quantified using two complementary approaches. First, conditioned behavior was measured on trial-by-trial basis as in cue-sucrose task. For cue-alcohol task, lick to CS was defined as the difference in lick counts between post-cue and pre-cue 1.25 s windows. For cue-shock task, P_freeze_ to CS was defined as the difference in immobility fraction between post-cue and pre-cue 10 s windows. We used a 10 s window in the early cue period because immobility differences were largest early in the cue (**fig. S11G,H**), consistent with a previous report that showed a transition from freezing to early cue to flight behavior to late cue when two cues serially precede a foot shock (*113*). Second, because trial-by-trial analysis does not account for baseline behavioral differences across group (e.g., during CS- and pre-cue periods; this difference is strongest for cue-shock task; see **fig. S11C**), we computed a session-by-session selectivity index (equation below; **Fig. 4E, fig. S11E,F**). Behavioral readouts were licks to CS in cue-alcohol task and P_freeze_ to CS in cue-shock task.

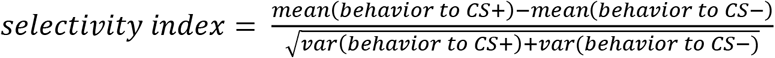

On a few sessions, recording files were corrupted and could not be accessed, and therefore excluded from analysis.

#### Behavioral analysis for reward identity specific contingency degradation task

Throughout the experiment, conditioned responding was measured as the mean rate of anticipatory reward port entries during the 20 second cues minus the entry rate during a 20 second pre-cue baseline (**fig. S10B**). Conditioned responding during the two recovery tests were averaged together.

#### Behavioral analysis for response elimination task

Lick to CS was computed in the same manner as in cue-sucrose task. To test whether behavior recovered during standard extinction following unpaired response elimination, we compared the average lick to CS during the last three days of unpaired response elimination phase with the average lick to CS on the first day of standard extinction (**fig. S10E**).

### Two-photon imaging data analysis

Raw imaging files acquired using Bruker Prairie View were motion-corrected, and regions of interests (ROIs) were detected using Suite2p (*114*). All detected ROIs were visually inspected by experimenters to confirm that their size and shape were consistent with neuronal soma and not dendritic segments or any other noise. Raw fluorescence signals (*F*) from each ROI were corrected for neuropil signal (*F_neu_*) as *F*^′^ = *F* − 0.7 × *F_neu_*. Then we calculated normalized fluorescence signal as the ratio (*F*^′^ − *F*^′^*_median_*)/(*F*^′^*_max_* − *F*^′^*_min_*) and normalized fluorescence was used for all subsequent analyses. If two ROIs recorded from adjacent planes (>50 μm apart) exhibited a cross correlation greater than 0.8 in their normalized fluorescence traces and were separated by less than 50μm in the x-y plane, the ROI with the lower variance in normalized fluorescence (presumed to have lower signal-to-noise ratio) was excluded.

Neurons recorded on each day were pooled across four animals (4048 on Day1, 4344 on Day2) for clustering analysis. For each neuron, pre-reward and post-reward activity were quantified as AUC of the normalized fluorescence signal within a 5 s window before and after the first lick following each reward delivery, respectively. If an animal did not lick within the first 10 s after reward delivery, that reward was excluded from analysis. Pre-reward and post-reward activity across trials (30 trials * 4 blocks) for all neurons were then concatenated and subjected to principal component analysis (PCA) for dimensionality reduction. The number of principal components retained was determined using an elbow detection method on the scree plot, which identifies the point of maximum deviation from a straight line connecting the first and last components. Neuronal activity was then projected onto the retained principal component axes, and neurons were clustered using the spectral clustering algorithm. To determine the optimal clustering parameters (number of clusters (k) and possible n nearest neighbors), we performed clustering across all parameter combinations for 10 iterations each, sweeping k from 2 to 20 and the number of nearest neighbors from [200,500,1000,1500,2000]. Clustering quality was evaluated using two independent metrics: 1) silhouette score and 2) co-clustering probability across iterations. Both measures were comparably high at k=2 and k=3 (**fig. S14B,C**); we therefore selected k=3 to obtain a finer classification of neuronal populations. At k=3, the silhouette score was maximized when number of nearest neighbors was 2000. Using these optimized parameters (k=3 and number of nearest neighbors=2000), we performed a final spectral clustering to obtain the clustering result. This clustering pipeline followed the procedure previously reported in (*80*).

To test whether individual neurons encoded reward rate through their phasic reward response, we compared the difference between post-reward and pre-reward activity across two distinct reward rate blocks (low vs. high) using a t-test. Neurons with a p value < 0.01 were classified as reward rate coding via reward response. Similarly, to identify neurons encoding reward rate through baseline activity, we compared pre-reward activity alone between the low- and high-reward rate blocks using the same statistical criterion (**fig. S14H, I**). Response to reward was defined as the difference between post-reward and pre-reward activity (**Fig. 5H, fig. S14J**) and baseline activity was defined as pre-reward activity (**fig. S14G**).

### Simulations

#### Tasks

Each experiment was simulated for 20 iterations. For ANCCR and latent cause models, we simulated 300 trials per CS type in each phase. For value RNN, each iteration consisted of 50 training epochs, with 5000 trials per CS type in each phase and a batch size of 12 episodes.

#### ANCCR

We previously proposed a learning model, ANCCR, based on learning of retrospective associations. ANCCR identifies cues that consistently precede meaningful events such as rewards, thereby learning a retrospective cue←reward association. This retrospective association can then be converted into a prospective cue→reward association through a Bayes’ rule: 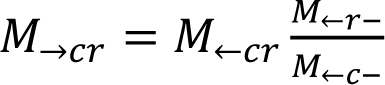, where *M*_←*r*−_ is proportional to the baseline rate of a reward in the environment, and *M*_←*c*−_is proportional to the baseline rate of a cue. Here, *M*_→*cr*_ corresponds to the successor representation (a measure of prospective association that is not baseline-corrected), while *M*_←*cr*_ represents the predecessor representation (a measure of non-baseline-corrected retrospective association). For intuitive clarity throughout the paper, we denoted these as P(cue→reward) and P(cue←reward), respectively. Because ANCCR is fundamentally driven by retrospective association, it predicts true learning and unlearning occur only when rewards are experienced. Consequently, the retrospective association functions as the original memory, providing an explanation for the rapid reacquisition observed after standard extinction.

We used the same ANCCR model described previously (*45*) but introduced one modification: the meaningfulness of an event was re-evaluated each time the event occurred, rather than treated as permanently meaningful once it first crosses a threshold. In the original implementation, meaningfulness became permanent when the sum of an event’s innate meaningfulness and its learned meaningfulness (reflected in the dopamine response) exceeded a threshold. This simplification was appropriate for scenarios in which meaningfulness does not change after learning—for example, rewards are innately meaningful, and cues become permanently meaningful after cue-reward association is formed. However, optogenetic inhibition of dopamine neurons at the time of reward suppressed the dopamine response below baseline, which in turn should reduce the total meaningfulness (innate+learned) of the reward and thereby suppress retrospective update. To capture this effect, we modified the model so that the meaningfulness of an event is computed dynamically on every occurrence, allowing meaningfulness of events (even the innately meaningful events like reward) to reduce when dopamine responses are arbitrarily reduced. In addition, we assumed that animals use their prior P(reward|reward) to rapidly restore the estimated reward rate when they returned to the reward-rich environment (e.g., reacquisition) after prolonged absence of rewards (e.g., extinction). For simplicity, we implemented this prior-based inference only on the first trial of the reacquisition phase for all groups, although the actual biological implementation may differ. For all simulations, we used the following parameters: w = 0.5, *α*_0_=4×10^-5^, M_opt_ = 0.5, dt = 0.2 s. T was defined as *M_opt_* ∗ ∑*_x_ λ_x_* / ∑*_x_ λ_x_*^2^, where *λ_x_* is an inverse of average inter-event interval for each event *x*. This ensures that the eligibility timescale matches the timescales operating in the environment. *α* was defined as 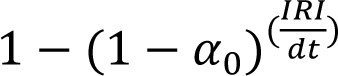 where IRI is inter-reward interval. This scaling ensures that learning of baseline event rates (e.g., *M*_←*r*−_ and *M*_←*c*−_), which are updated using the learning rate *α*_0_, and learning of retrospective association (*M*_←*cr*_), which is updated using learning rate *α*, proceed on comparable timescales. For additional details, see Methods and Supplementary Note of Burke et al (*45*). Dopamine response during optogenetic inhibition of dopaminergic neurons was modeled as -1.5 (1∼simulated dopamine response to an unexpected reward).

Learning speed was quantified using simulated baseline-corrected prospective association (successor representation contingency, SRC). Specifically, we computed the area under the SRC vs. trial curve for the first 90 trials of acquisition or reacquisition.

#### Latent cause models

We simulated three different latent cause models—Redish (2007) et al. (*40*), Gershman (2017) et al. (*38*), and Cochran (2019) et al (*37*)—as illustrated in **fig. S6A-D**. For each model, we systematically swept the model parameters most relevant to late state inference, as described below. All simulations for latent cause models were performed by modifying the scripts provided by Cochran et al. on Github (https://github.com/cochran4/OnlineLatentStateLearning). The modifications included the task inputs to match the experiments performed here and the parameter sweep described below.

Redish (2007) et al. (*40*)

- P*_P_* (state-creation threshold): determines when a new state is created based on the activation of existing states; lower values increase the likelihood of creating new states; default, 10^-8^; swept, [10^-10^,10^-8^,10^-7^, 10^-6^,10^-5^,10^-4^]

- *ξ*_0_ (decay rate of *δ*^―^, the running average of negative RPEs): controls how long past negative RPEs influence *δ*^―^; higher values integrate history over longer timescales, increasing evidence for new states; default, 0.9999; swept, [0.9999,0.99,0.9,0.8]

- *ξ*_1_ (scaling of *δ*^―^ updates): sets how strongly each negative RPE contributes to *δ*^―^; higher values increase the impact of each negative RPE, promoting state splitting; default, 1.5; swept, [1.5,1,0.5]

- *ξ_DB_* (attention squashing parameter): controls the steepness of the tanh transformation linking *δ*^―^ to cue attention; lower values make cue-attention changes more sensitive to *δ*^―^, increasing splitting; default, 1; swept, [1,0.5,0.25]

Gershman (2017) et al. (*38*)

- *α* (concentration parameter): governs the prior probability of assigning the current trial to a new latent cause; higher values increase the likelihood of inferring new causes; default, 0.1; swept, [0.01,0.1,0.5,1,1.5,2]

- *κ* (temporal kernel exponent): determines how strongly recent trials dominate the prior; higher values make the model favor the most recent latent cause, reducing splitting; default, 1; swept, [0.1,0.5,1,1.5,2,5,10]

- σ^2^_*x*_ (cue variance under each latent cause): prior uncertainty in cue observations; lower values increase sensitivity to small cue deviation, promoting an inference of new cause; default, 1; swept, [0.001,0.05,0.01,0.1,0.2,0.5,1]

- σ^2^_*r*_ (reward variance under each latent cause): prior uncertainty in reward outcomes; lower values make reward prediction errors more diagnostic, increasing state splitting; default, 1; swept, [0.001,0.05,0.01,0.1,0.2,0.5,1]

Cochran (2019) et al. (*37*)

- *γ* (latent state transition probability): controls the chance of switching states between consecutive trials; higher values increase trial-to-trial state transitions and new state inference; default, 0.05; swept, [10^-10^,0.001,0.05,0.01,0.1,0.2,0.5,1]

- *υ* (threshold for creating new state): determines how much mismatch is required to trigger a new state; lower values result in more frequent creation of new states; default, 0.2; swept, [0.1,0.2,0.5,10,1000]

- *δ* (unexpected-uncertainty threshold): controls how large prediction errors must be to start accumulating evidence for a state change; lower values make the model highly sensitive to deviations, increasing splitting; default, 0.6; swept, [0.05,0.3,0.6,1.5,5]

- *σ*_0_ (initial expected uncertainty): initial standard deviation of prediction error; lower values make early prediction errors more informative, facilitating new-state inference; default, 0.5; swept, [0.01,0.1,0.5,1,5]

The best-fitting parameter set was defined as the sweep combination that produced 1) the largest difference between groups in the probability of inferring the acquisition context, P(contextAcq), during CS-A trials in the CS-A extinction phase, and 2) the largest P(contextAcq) during CS-A trials in reacquisition for Group1. The acquisition context was defined by ranking all inferred contexts based on their total inferred probability during Acquisition and selecting the smallest set of top-ranked contexts whose cumulative inferred probability exceeded 90% of acquisition trials. Learning speed was quantified using the same procedure as for ANCCR, but substituting SRC with the value estimates generated by each model. For the optogenetic dopamine inhibition experiment, we simulated only Redish et al., model, as it was the only latent cause model that captured the behavioral results of Group1 and Group2. Optogenetic inhibition was modeled as a negative RPE with varying magnitudes [-10, -5, -2, -1.5, -1, -0.5, -0.3], and results with RPE=-1.5 (same magnitude as used in ANCCR simulation) were used in the main figure (1=modeled positive RPE evoked by an unpredicted reward).

For the Cochran et al. (*37*) model, we additionally simulated a 100% cue–reward probability version of the task to test whether the model could capture the behavioral patterns of Groups 1 and 2 under deterministic conditions. We focused on simulating the Cochran et al. model because it is the most recent latent-cause framework and is conceptually aligned with the Gershman et al. (*38*) model (both rely on Bayesian inference over latent causes). We did not re-simulate the Redish et al. (*40*) model for this condition, as we had already confirmed that it successfully captures the Group1 and Group2 results under the 50% probabilistic condition (although it does not capture either the Group3 pattern or Group2 with optogenetic manipulation). The same parameter sweep combinations were used as in the probabilistic (50%) condition. For **fig. S7D,E**, we selected the representative parameter set that captured the differential behavior between the 50% and 100% conditions (but see **fig. S7A-C** for full sweep results): the sweep in which no new state was inferred for CS-A trials in either Group1 or Group2 under the 50% condition, whereas a new state was inferred for CS-A trials in both groups under the 100% condition.

#### Value RNN

We simulated value RNN (*51*, *54*) as it shares a similar architectural logic with latent cause models—one component infers the latent state, and another learns the value associated with that inferred state—while recovering the correct state structure without requiring manually curated state labels as in traditional RPE models. We performed simulations by modifying the scripts provided by Hennig et al. (*51*) and Qian et al. (*54*) on Github (https://github.com/mhburrell/Qian-Burrell-2024). Briefly, we trained recurrent neural network models in PyTorch to estimate state-dependent value. Each Value-RNN consisted of 50 gated recurrent unit (GRU) cells, previously shown to capture belief-state representations (*51*, *54*), followed by a linear value readout. We used the same parameters as Qian et al, except that we reduced the time step from 0.5 s to 0.25 s to accommodate the 1.25 s cue duration in our task, and adjusted γ to 0.912 to match the 0.67 per s discount rate used by Qian et al (*54*). We did not jitter reward timing, as animals’ internal timing uncertainty was not a focus of this study.

Because weight updates occur at batch level rather than on a trial-by-trial basis in value RNN, and value learning evolved slowly, learning curves were plotted as a function of training epoch (one complete pass through the dataset, 21 batches), not trial. Similarly, quantification of learning was performed by measuring AUC of value vs. epoch curve. Dopamine response was measured as simulated reward prediction error at reward.

#### SOP

We simulated three different versions of the model (**fig. S17**): (1) naïve SOP (*95*), naïve SOP with context implemented as an additional cue (*96*), and (3) modified SOP that uses a symmetric update rule (*97*), in which cue-reward association is strengthened when both are in the same state (either in A1 or A2), and weakened when they are in different states. This modified update rule has been proposed as a mechanism of retrospective revaluation. For naïve SOP, we used the parameter set reported in Holmes et al. (*96*) (**fig. S17C,D**), which was extensively tested across different Pavlovian conditioning paradigms including extinction and reacquisition: p1_CS_ = 0.1, p1_US_ = 0.9, pd1 = 0.1, pd2 = 0.02, L_e_ = 0.9, L_i_ = 0.09, r1 = 1, and r2 = 0.1. To test whether pd2 drives the conditioned inhibition in G2 and thus slower reacquisition, we additionally simulated the same model but with an increased pd2 value (pd2 = 0.1; **fig. S17E,F**). When applying the Holmes et al. parameter set to SOP model with context implemented or the one with modified update rule, cue values did not reach stable asymptotes. We therefore explored parameter regimes that yield stable value representations for each model while producing results that mostly closely matched the empirical findings. For SOP with context implementation, we used the Holmes parameter set with increased pd2 (pd2 = 0.1; **fig. S17G,H**). For the SOP with modified update rule (**fig. S17I,J**), the following parameters were adjusted relative to the Holmes et al. parameter: p1_CS_ = 0.5, p1_US_ = 0.5, pd2 = 0.1, L_e_ = 0.04, L_i_ = 0.04, r1 = 1, r2 = 0.5. Notably, achieving stable value representation with the generalized update rule required a substantially lower excitatory learning rate (L_e_ = 0.04) compared to the original formulation (L_e_ = 0.9). Conditioned response was quantified as the proportion of reward elements in the A2 state during CS-A presentation.

## Notes

### Competing Interest Statement

The authors have declared no competing interest.

### Summary of Updates

Multiple new supplementary figures have been added showing additional analyses and experiments

